# Diatom Centromeres Suggest a Novel Mechanism for Nuclear Gene Acquisition

**DOI:** 10.1101/096016

**Authors:** Rachel E. Diner, Chari M. Noddings, Nathan C. Lian, Anthony K. Kang, Jeffrey B. McQuaid, Jelena Jablanovic, Josh L. Espinoza, Ngocquynh A. Nguyen, Miguel A. Anzelmatti, Jakob Jansson, Vincent A. Bielinski, Bogumil J. Karas, Christopher L. Dupont, Andrew E. Allen, Philip D. Weyman

**Author notes:** Present address: Designer Microbes Inc. London, Ontario, N6G4X8, Canada. Correspondence: Philip D. Weyman.

## Abstract

Centromeres are essential for cell division and growth in all eukaryotes, and knowledge of their sequence and structure guides the development of artificial chromosomes for functional cellular biology studies. Centromeric proteins are conserved among eukaryotes; however, centromeric DNA sequences are highly variable. We combined forward and reverse genetic approaches with chromatin immunoprecipitation to identify centromeres of the model diatom *Phaeodactylum tricornutum*. Diatom centromere sequences contain low GC content regions and an abundance of long contiguous AT windows, but lack repeats or other conserved sequence features. Native and foreign sequences of similar GC content can maintain episomes and recruit the diatom centromeric histone protein CENP-A, suggesting non-native sequences can also function as diatom centromeres. Thus, simple sequence requirements enable DNA from foreign sources to incorporate into the nuclear genome repertoire as stable extra-chromosomal episomes, revealing a potential mechanism for bacterial and foreign eukaryotic DNA acquisition.

## 1. Introduction

Centromeres play a crucial role in the cellular biology of eukaryotes by acting as a genomic site for kinetochore formation and facilitating effective transmission of replicated nuclear DNA to new cells. Centromere associated proteins are functionally conserved among eukaryote species (Cheeseman and Desai, 2008; Pluta et al., 1995; Westermann et al., 2003). Nearly all eukaryotes studied to date possess a version of a specialized centromeric histone protein known as centromere protein A (CENP-A, also described as CENH3), which binds to centromeric DNA and replaces the histone H3 at the site of kinetochore assembly (Earnshaw et al., 2013; McKinley and Cheeseman, 2016; Westhorpe et al., 2014). Conversely, the centromeric DNA sequences themselves are extremely variable and appear to evolve rapidly, even among similar organisms (Henikoff et al., 2001).

There are 3 general types of eukaryotic centromeres: point centromeres, holocentromeres, and regional centromeres. Point centromeres are uniquely characterized by specific conserved DNA sequences, and are found in limited fungal species including the budding yeast *Saccharomyces cerevisiae* and close relatives (Cleveland et al., 2003; Cottarel et al., 1989; Smith et al., 2012). In holocentromeric organisms, the kinetochore forms along the entire length of each chromosome; a notable example is the model organism *C. elegans* (Albertson and Thomson, 1982). Most eukaryotes have regional centromeres, which are commonly found as a single large DNA region on each chromosome (Reviewed in Sullivan et al., 2001 and Torras-Llort et al., 2009). Regional centromeres are variable in length and sequence even among closely related species; however, there are often predictable genetic features. For example, human centromeres contain large stretches of repetitive satellite DNA, ranging in size from hundreds of kilobases to megabases (Sullivan et al., 2001; Tyler-Smith et al., 1993; Willard, 1998). Centromeres of several plants and the insect model *Drosophila melanogaster* contain large arrays of satellite repeats interspersed with or adjacent to retrotransposons, which can vary substantially in copy number and organization (Ma et al., 2007). A common feature of many eukaryotic centromeric DNA is low GC content. Centromeres of *Schizosaccharomyces pombe* and other yeast species feature an unconserved core of AT-rich DNA sequence often surrounded inverted repeats (Clarke et al., 1986; Kapoor et al., 2015; Lynch et al., 2010; Nakaseko et al., 1986). The centromeres of the protist *Plasmodium* have no apparent sequence similarity besides being 2-4-kb regions of extremely low GC content (<3%)(Bowman et al., 1999; Iwanaga et al., 2010). Likewise, centromere regions of the red algal species *Cyanidioschyzon merolae* contain 2-3-kb of relatively low GC content but manifest no other apparent pattern (Kanesaki et al., 2015; Maruyama et al., 2008).

Centromere identification can also be useful for synthetic biology, enabling further discoveries and biotechnology applications. Artificial chromosomes provide a stable platform for introduction and maintenance of multigene constructs necessary for expression of biosynthetic pathways and large complex proteins (Coudreuse, 2009; Kouprina et al., 2014; Monaco and Larin, 1994; Yu et al., 2016). The experimental identification of eukaryotic centromeres has been extremely useful for developing molecular biology tools, particularly in the creation of artificial chromosomes. Circular and/or linear artificial chromosomes based on native centromeres, origins of replication, and in some cases telomeres have been developed for yeast (Murray and Szostak, 1983), mammalian cells including human cell lines (Harrington et al., 1997), plants (reviewed in Liu et al., 2013), and recently the protist *Plasmodium* (Iwanaga et al., 2012). Despite the great potential for eukaryotic algae in biotechnology, very little is known about algal centromeres and few resources are available to control gene expression from introduced autonomously replicating genetic constructs. In 1984, autonomously replicating plasmids utilizing chloroplast DNA were described for the green alga *Chlamydomonas reinhardii* (Rochaix et al., 1984). However, these vectors were not maintained stably and have not been commonly used. More recently, centromeres have been identified and characterized in the red alga *Cyanidioschyzon merolae* (Kanesaki et al., 2015; Maruyama et al., 2008), where each of the 20 chromosomes were found to contain one distinct region recruiting CENP-A. However, to our knowledge, these sequences have not yet been utilized for the construction of artificial chromosomes.

Identifying centromere composition and optimizing artificial chromosome construction would be particularly valuable for diatoms, which are an abundant group of eukaryotic phytoplankton with important ecological significance. Diatom research has facilitated major discoveries in algal physiology and genetics, and several species have been cultivated and genetically manipulated for the development of valuable bioproducts (Bozarth et al., 2009; Fu et al., 2015; Lopez et al., 2005). In our previous work, we discovered that a region of *S. cerevisiae* DNA containing low GC content enabled the stable maintenance of autonomously replicating episomes in diatoms (Diner et al., 2016; Karas et al., 2015). The DNA was introduced into the diatoms *Phaeodactylum tricornutum* and *Thalassiosira pseudonana* by bacterial conjugation, also suggesting a previously unexplored mechanism for horizontal gene transfer from bacteria. Diatom nuclear genomes contain large amounts of DNA derived from non-nuclear sources, including foreign sequences such as bacteria and viruses, and prokaryotic and eukaryotic DNA obtained from endosymbiotic events (e.g., mitochondria, chloroplasts, and additional secondary endosymbioses) (Armbrust, 2009; Armbrust et al., 2004; Bowler et al., 2008; Timmis et al., 2004). This genetic complexity and rapid evolution contributes to the ecological success of diatoms. Thus, elucidating mechanisms that may facilitate nuclear gene acquisition and episomal maintenance will advance our knowledge of diatom evolution and enable biotechnological innovation.

Here, we identify centromeric regions of diatom chromosomes using forward and reverse genetics approaches, and observe that diatom centromeres are characterized by a simple low-GC signal which is also found in the previously described synthetic diatom episomes (Diner et al., 2016; Karas et al., 2015). Furthermore, we show that non-nuclear diatom DNA and foreign DNA from a variety of sources with similarly low-GC content can mimic a diatom centromere, suggesting a permissive mechanism for nuclear gene acquisition. We conclude with a model suggesting that the frequency of contiguous A+T regions is important in addition to having a small region with average GC of <˜33%. This study significantly advances the understanding of diatom genomic features, facilitates the development of diatom molecular tools, and suggests a new mechanism for diatom acquisition of foreign genetic material.

## Results

### Identification of putative diatom centromeres in *Phaeodactylum tricornutum* chromosomes 25 and 26

We hypothesized that a centromeric region of a diatom chromosome would support maintenance of a nuclear episome, as this is a useful experimental method of confirming centromere function for other organisms (Clarke and Carbon, 1980; Iwanaga et al., 2012). To identify a diatom centromeric region we first examined the shortest *P. tricornutum* chromosomes with telomere-to-telomere assembly (25 and 26, (Bowler et al., 2008)), which were each previously cloned as five overlapping ˜100 kb DNA fragments (Karas et al., 2013). In our prior studies (Diner et al., 2016; Karas et al., 2015), sequences supporting episome maintenance in *P. tricornutum* were characterized by improved ex-conjugant colony yield compared to plasmids incapable of episome maintenance. Thus, we predicted 100-kb fragments from a single *P. tricornutum* chromosome that supported episome maintenance would yield similarly increased colony numbers in our standard conjugation assay. Out of the 5 large fragments spanning each chromosome, one fragment from each chromosome produced increased ex-conjugant diatom colonies: plasmid Pt25-100kb-1 (containing the 1^st^ 100-kb fragment of Chromosome 25) (Figure 1A, 1C) and plasmid Pt26-100kb-5 (containing the 5^th^ fragment of chromosome 26) (Figure 1B, 1D). The Pt25-1 plasmid resulted in 14-32-fold more colonies than plasmids containing other 100-kb fragments from chromosome 25 (Figure 1A, 1C), and plasmid Pt26-5 resulted in 26-100-fold higher colony numbers than other chromosome 26 fragments (Figure 1B, 1D).

**Figure 1:**
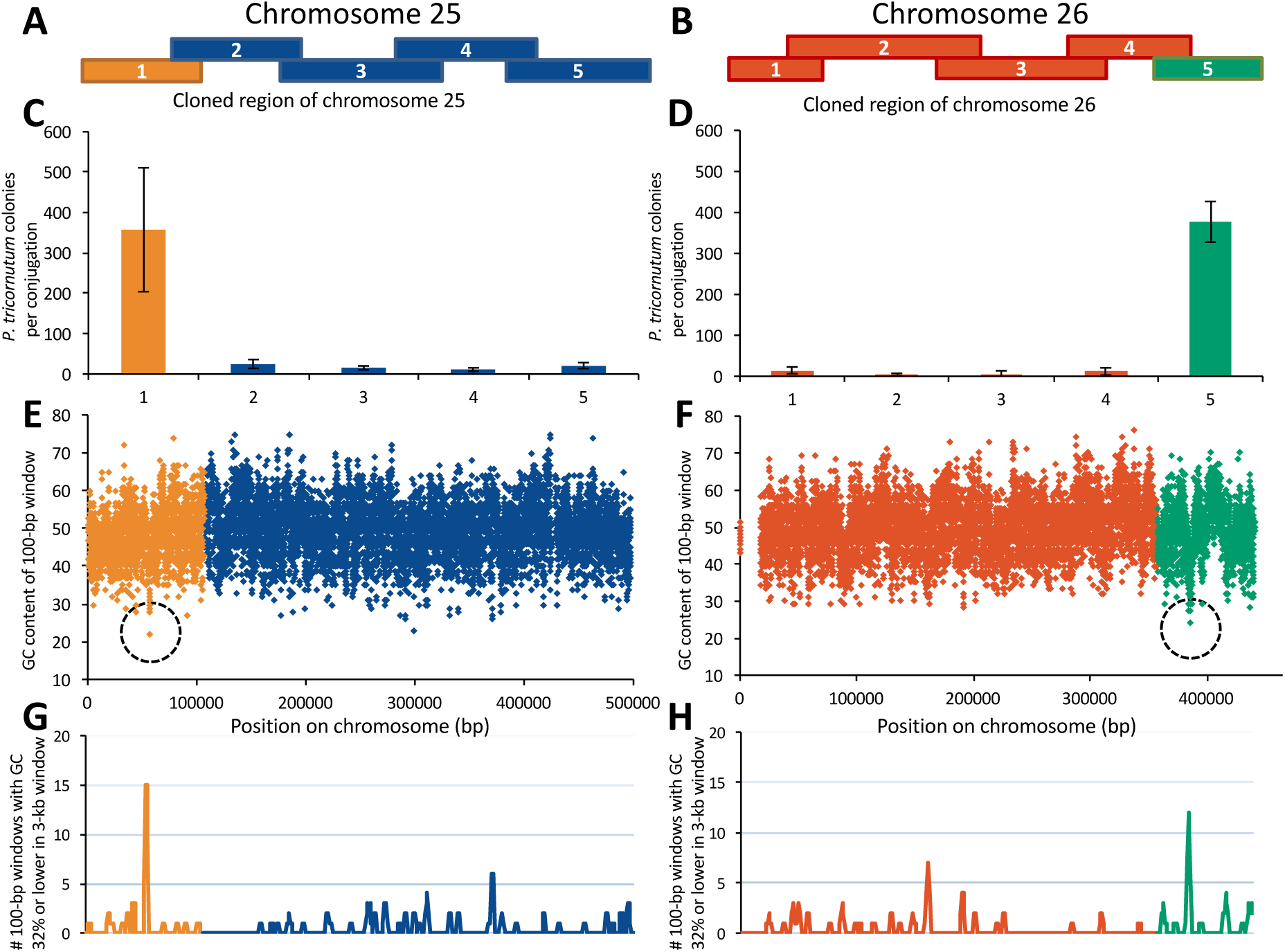
Regions of *P. tricornutum* chromosomes enriched for low GC support episomal maintenance. **A and B**. Chromosomes 25 and 26 were cloned as 5 overlapping ˜100-kb fragments. **C and D**. Number of resulting *P. tricornutum* colonies per conjugation for episomes containing indicated region of chromosome 25 or 26. Error bars indicate standard deviation of four independent conjugation reactions for each fragment. **E and F.** GC content was calculated for chromosomes 25 and 26 in 100-bp sliding windows that overlapped by 50 bp. Dashed circles indicate the lowest GC content for the chromosome in a 100-bp window. **G and H.** Number of 100-bp windows with GC content of 32% or lower within a larger sliding 3-kb window that advanced by 1 kb each step.

Both Pt25-100kb-1 and Pt26-100kb-5 fragments encompass regions of low GC content. We calculated the GC content of the genome in 100-bp windows overlapping by 50-bp, and found that windows with the lowest GC content were found on fragments enabling episome maintenance (Figure 1E, 1F). When calculating GC percentage with larger window sizes (10-kb to 0.5-kb), an obvious dip in GC content was not apparent on chromosomes 25 and 26 (Supplementary Figure 1); this was also true for the other chromosomal scaffolds (data not shown). We quantified the number of 100-bp windows less than or equal to 32% GC within a 3-kb larger window, and observed clear peaks for chromosomes 25 and 26 (Figure 1G, 1H).

To clarify whether these specific chromosomal regions enriched in low GC content enabled episome maintenance, three 10-kb DNA subsequences of Pt25-100kb-1 were cloned into plasmids otherwise incapable of maintenance (pPtPBR2, Diner et al., 2016): 1 sequence encompassing the bioinformatically identified low GC region (Pt25-10kb-12) (Figure 1E), and 2 other randomly selected sequences (Pt25-10kb-6 and Pt25-10kb-9) (Supplementary Figure 2). Pt25-10kb-12 conjugation led to 85-fold more colonies than the negative control, while the other plasmids showed no increase (Supplementary Figure 2). We further tested the low GC region found on Pt25-10kb-12 by assembling a 1-kb sub-region containing the lowest GC content region of chromosome 25 into pPtPBR2. This plasmid, Pt25-1kb, yielded 27-fold more colonies than the empty vector control (Supplementary Figure 2). Another plasmid containing the 1-kb region encompassing the lowest GC content region of chromosome 26, Pt26-1kb, resulted in 68-fold more colonies than the empty vector control. Thus, for chromosomes 25 and 26, regions containing the lowest GC content were the only regions supporting episome maintenance. To confirm that these plasmids were maintained in the diatoms over extended periods of time, two clones of Pt25-1kb were passaged for 30 days without selection. Antibiotic resistant colonies were recovered at percentages similar to prior studies (Supplementary Table 1) (Diner et al., 2016; Karas et al., 2015), and plasmids were recovered after the passaging period, demonstrating the stable maintenance of episomes in these lines (i.e., not integrated into genomic DNA).

### Identification of diatom centromeres using ChIP-seq and reverse and forward genetics

*P. tricornutum* genomic DNA sequences enabled episome maintenance in the diatom, suggesting these regions were functioning as centromeres. Nearly all eukaryotes previously studied incorporate the centromeric histone CENP-A (CENH3) into centromeric nucleosomes, and we tested this in *P. tricornutum* to confirm centromere functionality. We constructed an episome containing the *CEN6-ARSH4-HIS3* maintenance sequence and a translational fusion of *P. tricornutum* CENP-A and yellow fluorescent protein (YFP) regulated by a *P. tricornutum* promoter and terminator. After transfer to *P. tricornutum* using bacterial conjugation (see Materials and Methods), we performed chromatin immunoprecipitation (ChIP) assays on ex-conjugant lines using GFP epitope antisera, followed by high-throughput DNA sequencing to identify all sequences *P. tricornutum* genome sequences that recruit the centromeric histone.

ChIP-seq analysis revealed 25 regions that were enriched for sequence reads (peaks) among the previously reported 33 nuclear chromosome scaffolds (Bowler et al., 2008) (Figure 3, Supplementary Figure 3). Of the 12 chromosome scaffolds with telomere-to-telomere assembly, all but one (chromosome 11) had ChIP-seq peaks, including chromosomes 25 and 26. Two regions recruiting CENP-A were also found within the non-scaffold assemblies (“bottom drawer” sequences)(Obtained from the JGI *P. tricornutum* genome website: http://genome.jgi.doe.gov/Phatr2/Phatr2.home.html) (Supplementary Figure 3). A ChIP-seq peak was also identified within the *S. cerevisiae CEN6-ARSH4-HIS3* region on the episome used to express the YFP-CENP-A fusion protein (Figure 3). No mitochondrial or chloroplast sequences recruited CENP-A, which was expected as these genomes do not contain nucleosomes. Most ChIP-seq peaks on a genome-wide scale co-localized with the presence of at least ten 100-bp windows with GC content less than or equal to 32% GC in a larger 3-kb region (Supplementary Figure 3).

**Figure 2:**
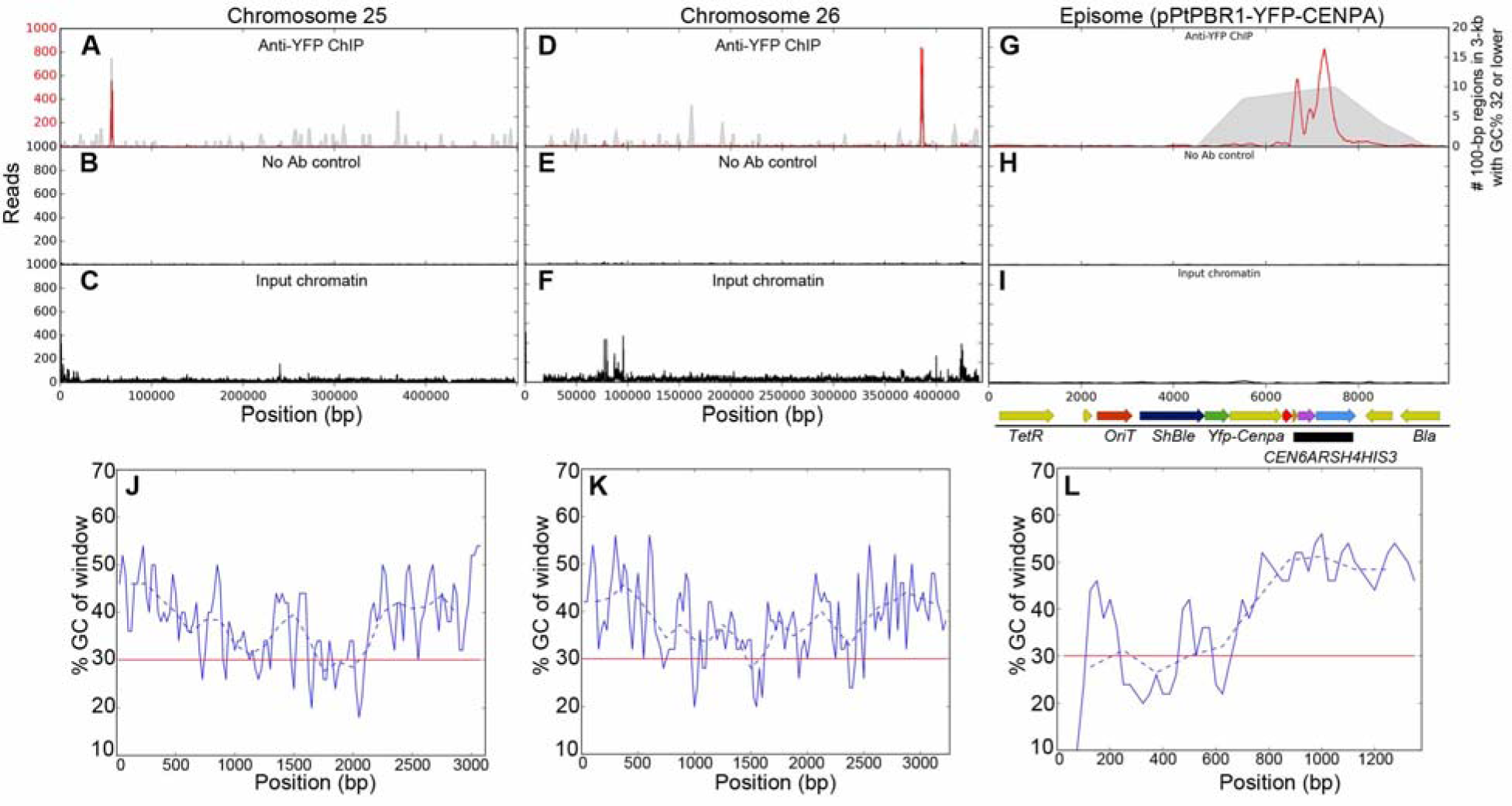
ChIP-seq and GC data for chromosomes 25, 26, and the episome. For each of chromosomes 25, 26, and the episome, ChIP-seq reads at each position for treatments with the YFP antibody (red) were plotted on the same graph as the number of 100-bp windows with GC 32% or lower in a larger 3-kb window (gray) (**A, D, G**). Graphs of the number of reads for the no-antibody ChIP-seq control (**B, E, H**) and input chromatin (**C, F, I**) were plotted using the same position scale as the anti-YFP ChIP-seq. For the episome, the positions of the genetic features are indicated below the input chromatin (the black bar indicates the *CEN6-ARSH4-HIS3* region). **J, K, L.** For the peaks identified by the CENP-A-YFP ChIP-seq in chromosomes 25, 26, and the episome, GC content for 100-bp windows (50 bp overlap, solid blue line) or 250-bp windows (125-bp overlap, dashed blue line) was plotted with a reference line at 30% in red.

**Figure 3:**
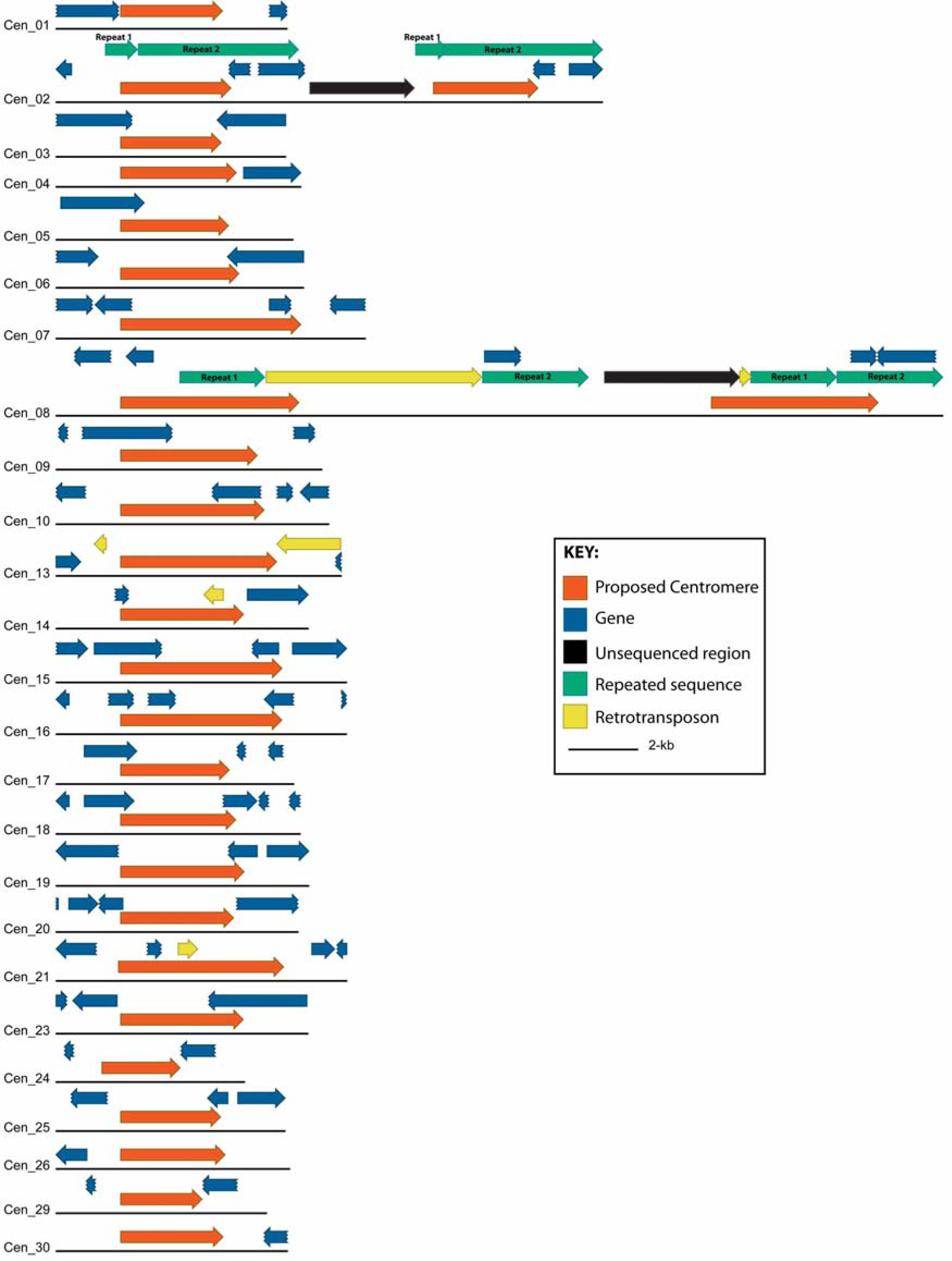
Location and genomic features of all *P. tricornutum* centromeres identified in chromosomal scaffolds. Centromeres identified by ChIP-seq (crimson) are plotted along with PHATR2 gene models (blue) for the regions spanning 2-kb upstream and 2-kb downstream of the annotated centromere regions (See Supplementary Table 3 for centromere genomic coordinates). For chromosomes 2 and 8, large direct repeated regions (teal) are plotted with regions of unresolved sequence (black) and retrotransposon annotations (yellow). Scale bar, 2-kb, noted in legend.

To verify the ChIP-seq data, we conducted ChIP-qPCR on two regions with ChIP-seq peaks, one in the genome (Pt25-1kb) and one in the episome (*ARSH4*), and a region of genomic and episomal DNA without ChIP-seq peaks as a control (See materials and methods) (Supplementary Figure 4A, 4B). DNA from the low-GC *ARSH4* episomal region was in greater abundance by >50-70-fold compared to the negative control (Supplementary Figure 4C, 4D, 4E). Similarly, the Pt25-1kb region was enriched >200-500-fold compared to the genomic DNA negative control (Supplementary Figure 4C, 4D, 4E). Thus, ChIP-qPCR confirmed the ChIP-seq results for the CENP-A enriched regions of both episomal and native *P. tricornutum* chromosomal targets.

Of the 25 chromosome scaffolds with ChIP-seq hits, 23 had only one associated ChIP-seq peak that was between 2.4-5.6 kb (Figure 3, Supplementary Figure 3). Chromosomes 2 and 8 each had two adjacent ChIP-seq peaks (Figure 3). Both putative centromeres on chromosome 2 (2a and 2b) are contained within a larger direct repeat and separated by a sequencing gap (indicated by Ns in the *P. tricornutum* genome sequence) (Figure 3). These sequences were highly similar to each other, with ˜2.9 kb aligning along the 3.4 kb sequence at >99% sequence identity. The 2 putative centromeres on chromosome 8 (8a and 8b, respectively) are each partially contained within long direct repeats at the 3’ end of the centromere. The 5’ end of 8b is adjacent to a region of unknown sequence (Figure 3). The 8a and 8b centromere sequences were also highly similar, with alignment across about half of the centromere sequence at 96.5 % identity.

Apart from these potentially tandem centromere cases, most *P. tricornutum* centromeres were unique, having no similarity to other centromere sequences, with two exceptions. Predicted centromeres from chromosomes 24 and 29 shared 99.2% sequence identity over the entire 2.4 kb region and differed by only 14 mismatches. Additionally, the centromere from chromosome 30 shared a 1.6 kb region of high identity (97%) to a bottom drawer sequence bd23x34, which was one of the 2 bottom drawer sequences with an associated ChIP-seq hit. Centromeres in *P. tricornutum* were mostly located in intergenic spaces (Figure 3). Direct repeats were detected in approximately one-third of the centromeres, but the repeat number was low (usually a single sequence found twice) and the repeat period was variable and small (16-400 bp) (Supplementary Table 2). Genomic coordinates of all predicted centromeres, including ChIP-seq read regions and bioinformatically predicted regions containing low GC content are noted in Supplementary Table 3.

We used forward genetics to test whether sequences in the *P. tricornutum* genome including and in addition to those identified by ChIP-seq could support episomal maintenance. We prepared a *P. tricornutum* genomic library with 2-5 kb inserts using a non-episome vector (pPtPBR2) and conjugated the library into *P. tricornutum* cells. Episomes were identified by extracting plasmids from antibiotic resistant *P. tricornutum* colonies and transforming *E. coli*; only DNA maintained as circular episomes in *P. tricornutum* was expected to yield *E. coli* colonies. We amplified and sequenced *P. tricornutum* genomic library inserts from *E. coli* colonies and identified 35 unique insert sequences from 99 recovered plasmids (Supplementary Table 4). Of these 35 unique insert sequences, 10 mapped to the nuclear genome chromosomal scaffolds and 1 mapped to the un-scaffold “bottom drawer” assemblies. Eighteen sequences mapped to the chloroplast genome and 6 mapped to the mitochondrial genome.

Reverse genetics was used to functionally test whether the sequences identified by ChIP-seq and the *P. tricornutum* forward genetics library could maintain episomes. Forty sequences, including all ChIP-seq peaks, potential ChIP-seq artifacts, and *P. tricornutum* forward genetic library sequences including selected mitochondrial and chloroplast DNA sequences were cloned into the non-episomal plasmid pPtPBR2 (See materials and methods). Most plasmids containing ChIP-seq identified sequences resulted in 7-162-fold more diatom ex-conjugant colonies than the pPtPBR2 negative control (Supplementary Figure 5). We also tested random regions of chromosome 1 as negative controls (Test-37, −38, and −39) and regions suspected to be ChIP-seq mapping artifacts based on high read counts in both input and anti-YFP immunoprecipitation treatments (Test-4, −10, and −16). Both classes of sequences were unable to support episome maintenance; ex-conjugant numbers were similar to the negative control and much lower than the positive control pPtPBR1 (Supplementary Figure 5). Ex-conjugant colony numbers following conjugation with the pPtPBR1 positive control (containing *CEN6-ARSH4-HIS3)* were not notably different from the episomes containing putative *P. tricornutum* centromeres. One insert sequence from chromosome 11 contained a region of GC content similar to, but slightly higher than, the centromeres (Test 40). However, this region contained no ChIP-seq peak and was unable to maintain an episome (Supplementary Figure 5).

We also tested the *P. tricornutum* regions recovered from the forward genetic screen for the ability to maintain episomes. All chloroplast and mitochondrial DNA sequences, the “bottom drawer” sequence, and 8 of the 10 nuclear genome sequences contained low GC content of 28-41% (Supplementary Table 4) across the entire insert region. These 8 nuclear genome sequences and the “bottom drawer” sequence mapped to identical regions as the ChIP-seq peaks (Supplementary Figure 3). The two remaining inserts (Test 18 and Test 20) from the nuclear genome had GC content typical of the *P. tricornutum* nuclear genomic DNA (47%), and did not map to a ChIP-seq peak (Supplemental Table 3). We re-tested whether the two high GC nuclear genome inserts as well as two sequences each from the chloroplast (Test 33 and Test 34) and the mitochondrion (Test 35 and Test 36) could support episome maintenance. Both mitochondrial and both chloroplast sequences supported episomes (Supplementary Table 4); however, the high GC nuclear sequences did not, and we predict that their appearance in the library was likely due to plasmid carry-over from the initial conjugation (Supplementary Table 4)

### Foreign DNA sequences examined for episome maintenance

Since the *CEN6-ARSH4-HIS3* sequence from *S. cerevisiae* supported episome maintenance in *P. tricornutum*, we hypothesized that other foreign DNA sequences with similarly low GC composition could as well. Deletion analysis of the *CEN6-ARSH4-HIS3* region previously revealed that low GC regions of >˜500 bp enabled maintenance. To test this pattern in the present study, we examined 24 sequences from *Mycoplasma mycoides* JCVI Syn1.0 (NCBI accession #CP002027) of various sizes (0.5-1 kb) and GC content (15-50%) for their ability to maintain diatom episomes. All sequences of less than 28% GC content regardless of the size resulted in high numbers of ex-conjugant colonies consistent with episome maintenance (Figure 4). Most sequences of 28% and 30% GC also resulted in large numbers of *P. tricornutum* ex-conjugant colonies with two exceptions that produced colony numbers similar to the negative control: a 500-bp 28% GC fragment (1.3 fold below control), and a 500-bp 30% GC fragment (1.2 fold above control) (Figure 4). Additionally, one 700-bp 30% GC fragment produced only 3.3-fold more colonies than the control, a relatively low colony increase. The fragments containing either 40% or 50% GC content sequences produced ex-conjugant colony numbers similar to the negative control. Thus, with a few exceptions (discussed below), DNA sequences of ˜30% GC or lower were required and sufficient to support *P. tricornutum* episomes.

**Figure 4:**
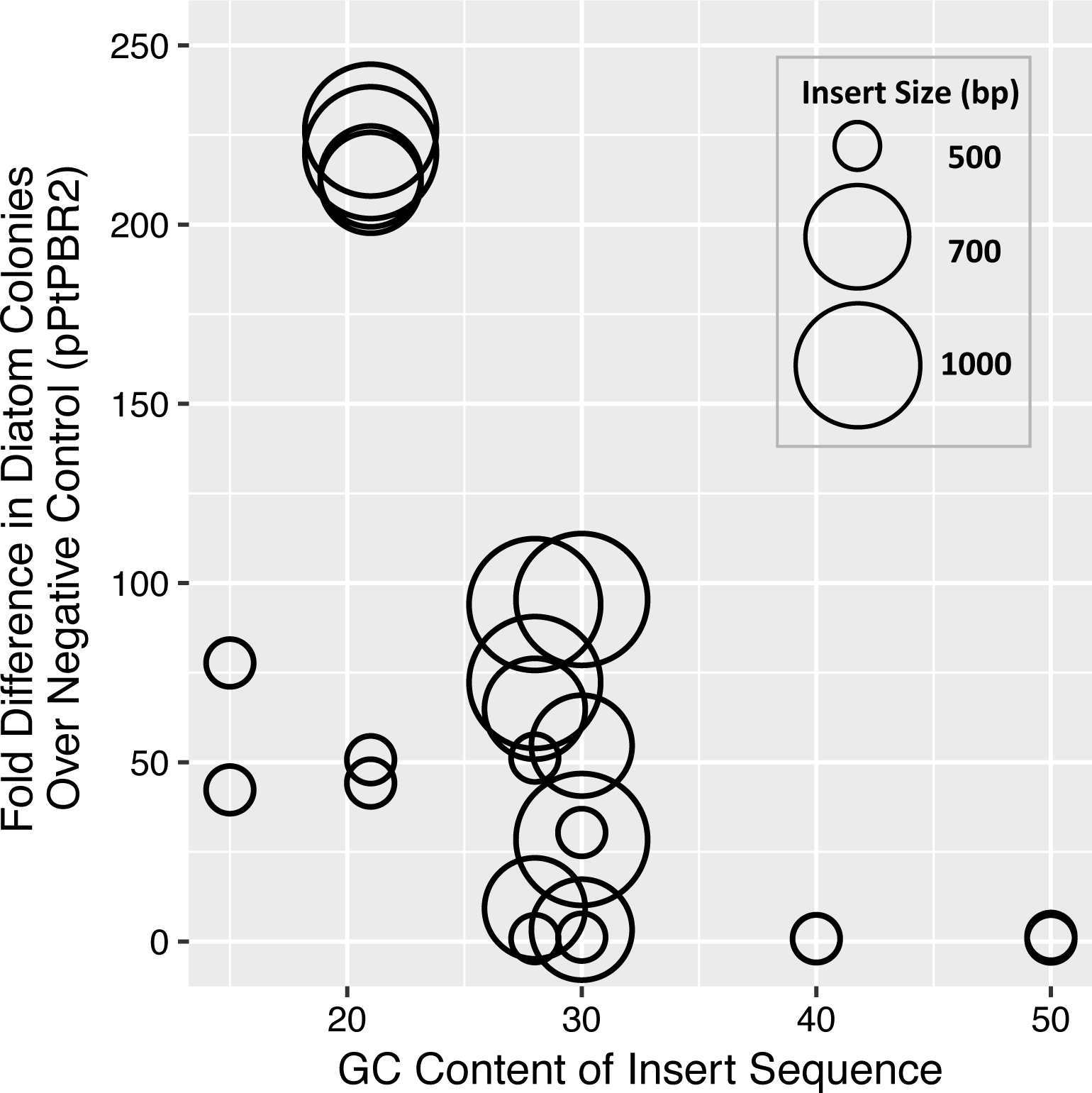
Maintenance of episomes containing *Mycoplasma mycoides* DNA sequences. GC content of *M. mycoides* DNA sequences tested for maintenance ability and number of diatom exconjugant colonies obtained after conjugation shown as fold increase in colony numbers over the pPtPBR2 negative control. The size of each circle represents the size of the insert sequence tested: large circles = 1000 bp, medium circles = 700 bp, and small circles = 500 bp.

The above results suggest that many sequences of at least 500-bp (the smallest fragment tested) of low GC DNA could maintain an episome in *P. tricornutum*, including sequences with environmental relevance. We examined whether a marine bacterial conjugative plasmid could support episome maintenance by searching the *Alteromonas macleodii* conjugative plasmid pAMDE1 for low GC content regions (Figure 5A). We then identified and cloned two 500-bp regions, AM-1 and AM-2, with 26.2% and 28.8% GC, respectively; conjugation of plasmids containing either region yielded 6-17-fold more ex-conjugant *P. tricornutum* colonies than the pPtPBR2 negative control with no maintenance sequence elements (Figure 5A). We also tested whether regions of plasmids previously isolated from the diatom *Cylindrotheca fusiformis* (Hildebrand et al., 1992; Jacobs et al., 1992) could support episomes in *P. tricornutum.* Two plasmids, pCF1 and pCF2, containing low GC, 560-bp regions (28.9 % and 28.4 % GC, respectively) were constructed (see materials and methods) and each yielded 7-12-fold more *P. tricornutum* ex-conjugant colonies than the pPtPBR2 negative control (Figure 5).

**Figure 5:**
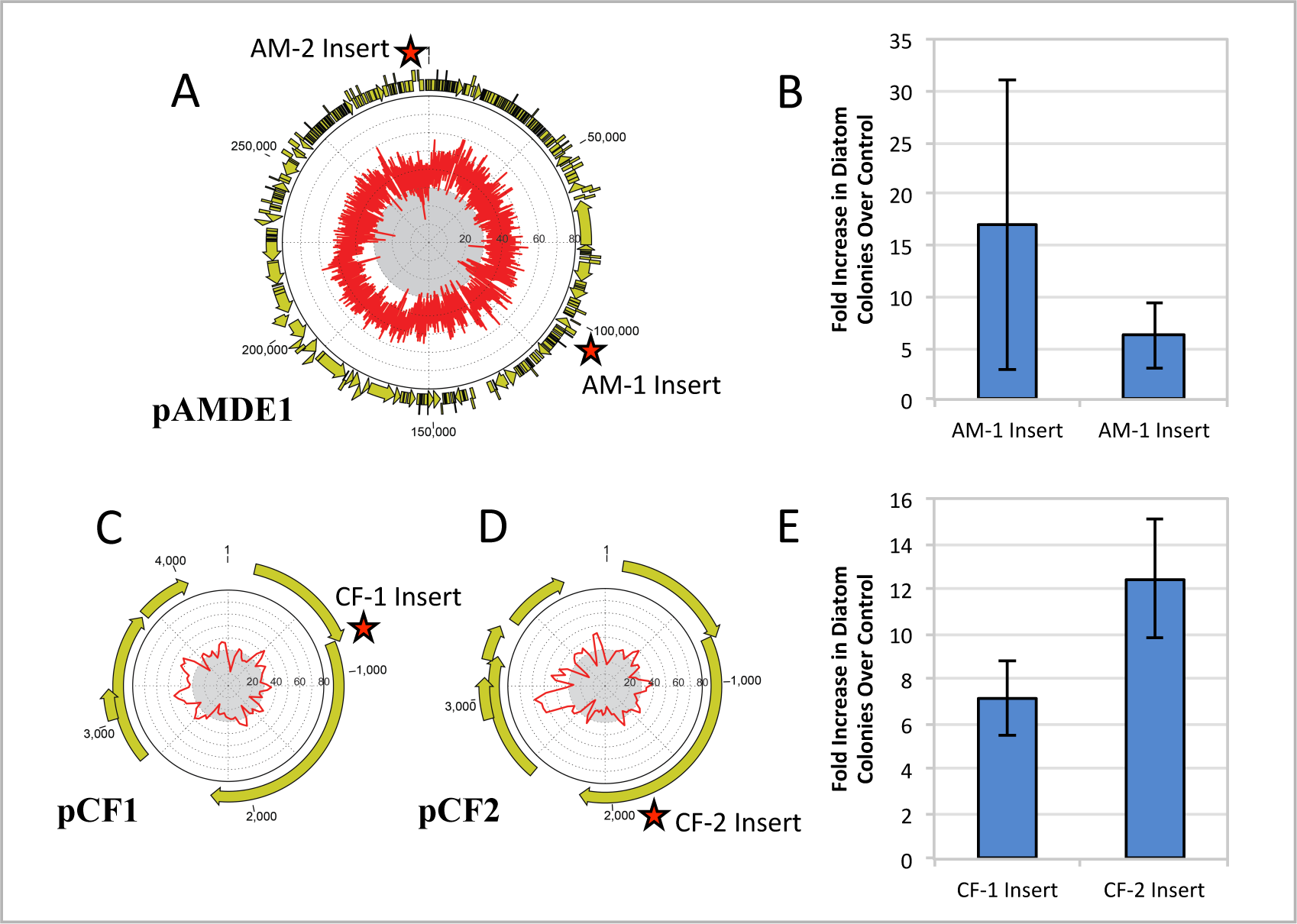
Maintenance of episomes containing *A. macleodii* and *C. fusiformis* DNA sequences. **A.** Map of the *Alteromonas macleodii* plasmid pAMDE1 with GC content mapped on the interior. Stars (*^AM-1 Insert^ or *^AM-2 Insert^) represent the regions used to test episome maintenance in plasmid pPtPBRAM-1 and pPtPBRAM-2, respectively. GC content was calculated for 100-bp windows that overlapped by 50-bp, and GC content below 30% in the plasmid figures is indicated by the central gray shaded circle. **B.** Ex-conjugant colony yield from conjugation of AM-1 and AM-2 Inserts, shown as fold increase in colony numbers over the pPtPBR2 negative control. Error bars show standard deviation from three biological replicates. **C.** Map of the *Cylindrotheca fusiformis* plasmid pCF1, *^CF-1 Insert^ represents the region used to test episome maintenance in plasmid pPtPBR-CF1. **D.** Map of the *Cylindrotheca fusiformis* plasmid pCF2, *^CF-2 Insert^ represents the region used to test episome maintenance in plasmid pPtPBRCF-2. **E.** Ex-conjugant colony yield from conjugation of episomes containing CF-1 and CF-2 Inserts are shown as fold increase in colony numbers over the pPtPBR2 negative control. Error bars show standard deviation from three biological replicates.

We examined maintenance properties of episomes supported by foreign DNA sequences. *P. tricornutum* lines containing plasmids with two different *Mycoplasma* inserts (Myco-15-500bp-2 and Myco-21-500bp-2) were examined for episomal maintenance in the absence of antibiotic selection. After 30 days of serial passage without antibiotic, between 45% and 75% of cells maintained the episome, similar to maintenance of episomes containing the *CEN6-ARSH4-HIS3* sequence (Diner et al., 2016; Karas et al., 2015) and the native *P. tricornutum* centromere sequence from chromosome 25 (Supplementary Table 1). We also tested maintenance of the *A. macleodii* and *C. fusiformis* sequences, and with the exception of colony 8 of the AM-2 plasmid, all episomes were maintained in the absence of antibiotic selection with retention rates between 24-84%, similar to previous reports (Diner et al., 2016; Karas et al., 2015). In colony 8 containing the AM-2 episome, only 3% of cells retained the episome after 30 days of serial passage without antibiotic. For all clones, as well as all other experiments where conjugations resulted in a high number of ex-conjugant diatom colonies relative to the negative control, episomes were successfully recovered in *E. coli*, confirming their stable extra-chromosomal maintenance. Thus, foreign low GC DNA sequences support episomal maintenance in a manner similar to native *P. tricornutum* centromeric sequences in the absence of antibiotic selection.

### Bioinformatic analysis of episome-supporting sequences

A common theme among sequences supporting episome maintenance regardless of source was their low GC content. ChIP-seq and forward genetic screening identified sequences longer than the minimal length required for episome maintenance; our results indicate that 500-bp sequences can maintain episomes. Thus, we searched within these sequences for the 500-bp sub-region with the lowest GC content (Supplementary Table 5). When viewed together based on ability to maintain an episome, all inserts from the native diatom “Test” series and all foreign DNA inserts examined (including *M. mycoides*, *C. fusiformis*, and *A. macleodii* plasmid pAMDE1 source DNA) indicated a clear pattern of low GC content supporting episome maintenance regardless of whether the source was foreign or native (Figure 6). However, three *M. mycoides* DNA sequences (28-500-2, 30-500-2, and 30-700-2) that were predicted to be maintained based on low average GC content produced low numbers of ex-conjugant colonies after conjugation. DNA sequencing confirmed the identity of the non-functioning *Mycoplasma* inserts and other critical plasmid features (data not shown).

**Figure 6:**
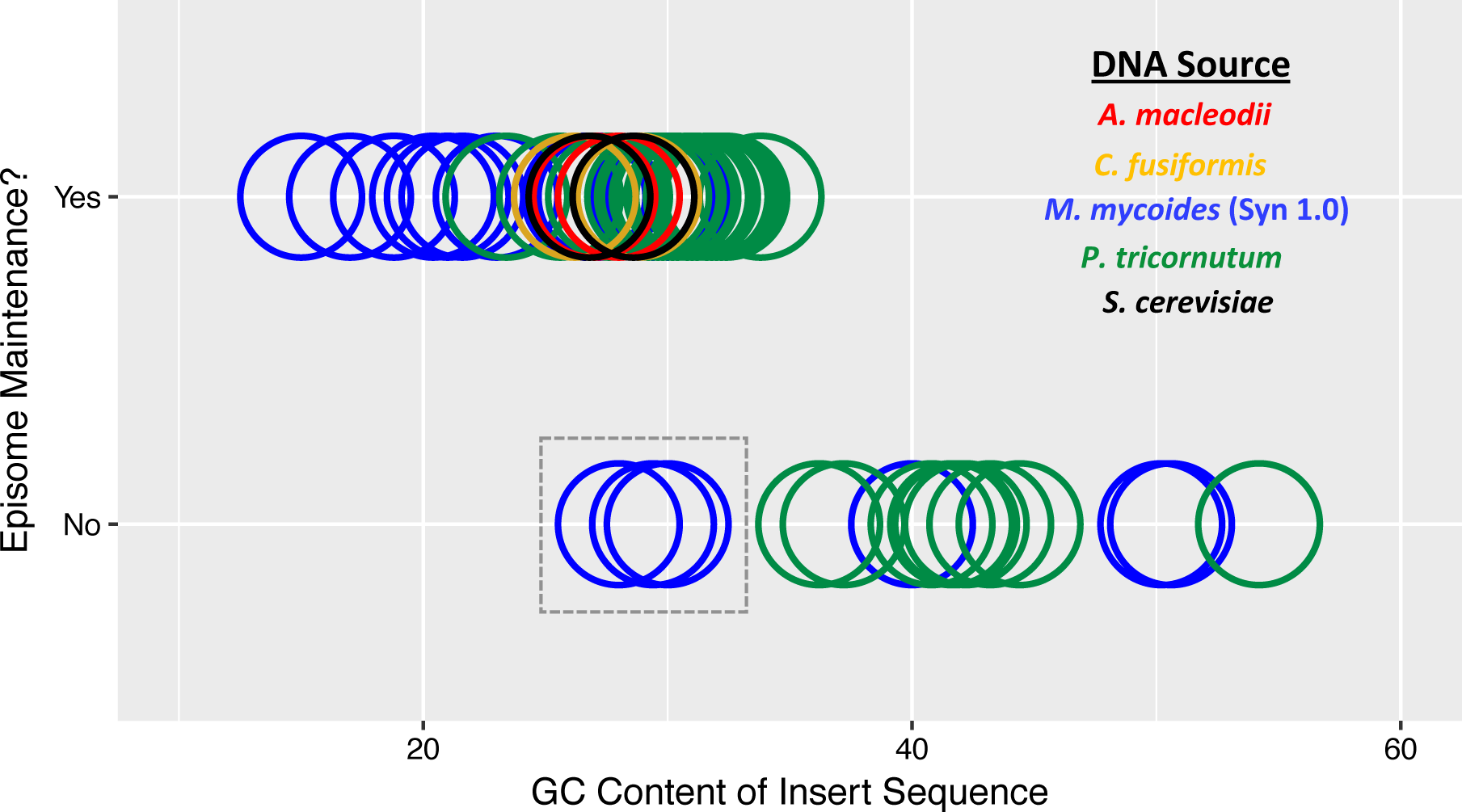
Relationship between GC content and episome maintenance. The 500-bp sub-region with the lowest GC content was identified for each insert sequence in the *P. tricornutum* Test series (which includes ChIP-seq peaks, forward genetic library sequences, designed negative controls, and potential ChIP-seq artifacts) and foreign DNA inserts (including *M mycoides, C. fusiformis*, and *A. macleodii* plasmid pAMDE1 source DNA). See Supplementary Table 5 for data included in the figure. This lowest GC content sub-region was plotted as a function of whether the DNA could support episomal maintenance in *P. tricornatam.* DNA sequences are colored by the organism from which each insert sequence originated.

Because these sequences were predicted to support episome maintenance based on average GC content, their failure to maintain episomes suggested that an additional signal besides average GC is required. Because native *P. tricornutum* centromeres do not have repeats or other structures and attempts to identify a conserved sequence motif using BLAST (Altschul et al., 1990) and MEME (Bailey et al., 2009) were unsuccessful, we examined k-mer usage to determine if very short sequences were over-represented in DNA fragments supporting episomes. We chose a k-mer length of 6 because it was the longest string that could still be well-represented in a sequence of 500-bp. We identified unique 6-mers over-represented in native *P. tricornutum* centromeres by requiring their retention to be statistically significant (P<0.001) when compared to randomly-selected *P. tricornutum* genomic sequence (47% GC) and randomly generated sequences of 47% GC. Because the overall GC content is lower for centromeric ChIP-seq peaks (39% GC average) compared to the genomic regions (47% GC), we also required the 6-mers to be significantly over-represented in the centromeres relative to a randomly generated set of 39% GC sequences. This allowed us to identify 6-mers over-represented in the *P. tricornutum* centromeres that were unexplained by GC content difference from the genomic DNA (Supplementary Table 6). We then examined the recruitment of this set of centromere-enriched 6-mers in two sets of *Mycoplasma* fragments. One set contained the two 28% GC sequences and one 30% GC sequence that did not support episome maintenance despite having a sufficiently low average GC content (“Myco-No” set). The second set comprised the remaining 9 *Mycoplasma* sequences with 28% and 30% average GC that successfully supported episome maintenance (“Myco-Yes” set). Unique 6-mers that were over-represented in the Myco-Yes set were characterized by very low GC content (i.e., the most abundant 6-mers in the “Myco-Yes” set were composed entirely of A+T bases) (Figure 7). When examining the number of consecutive A+T nucleotides in the *Mycoplasma* sequences that supported episome maintenance compared to those that did not, stretches of 6 or more consecutive A+T bases were more frequent in the *Mycoplasma* fragments that supported episome maintenance (i.e., “Myco-Yes”, Supplementary Table 7). The lower distribution of consecutive A+T bases in the “Myco-No” set was also observed when compared to a set of randomly generated sequences of 30% GC (Supplementary Table 7). Thus, the “Myco-No” samples that failed to support episome maintenance appear to have fewer long stretches composed of A+T residues despite having the same average GC content as fragments that supported episome maintenance in *P. tricornutum*. This observation is a valuable advancement in understanding the minimal sequence requirements for diatom centromeres and episomal maintenance.

**Figure 7:**
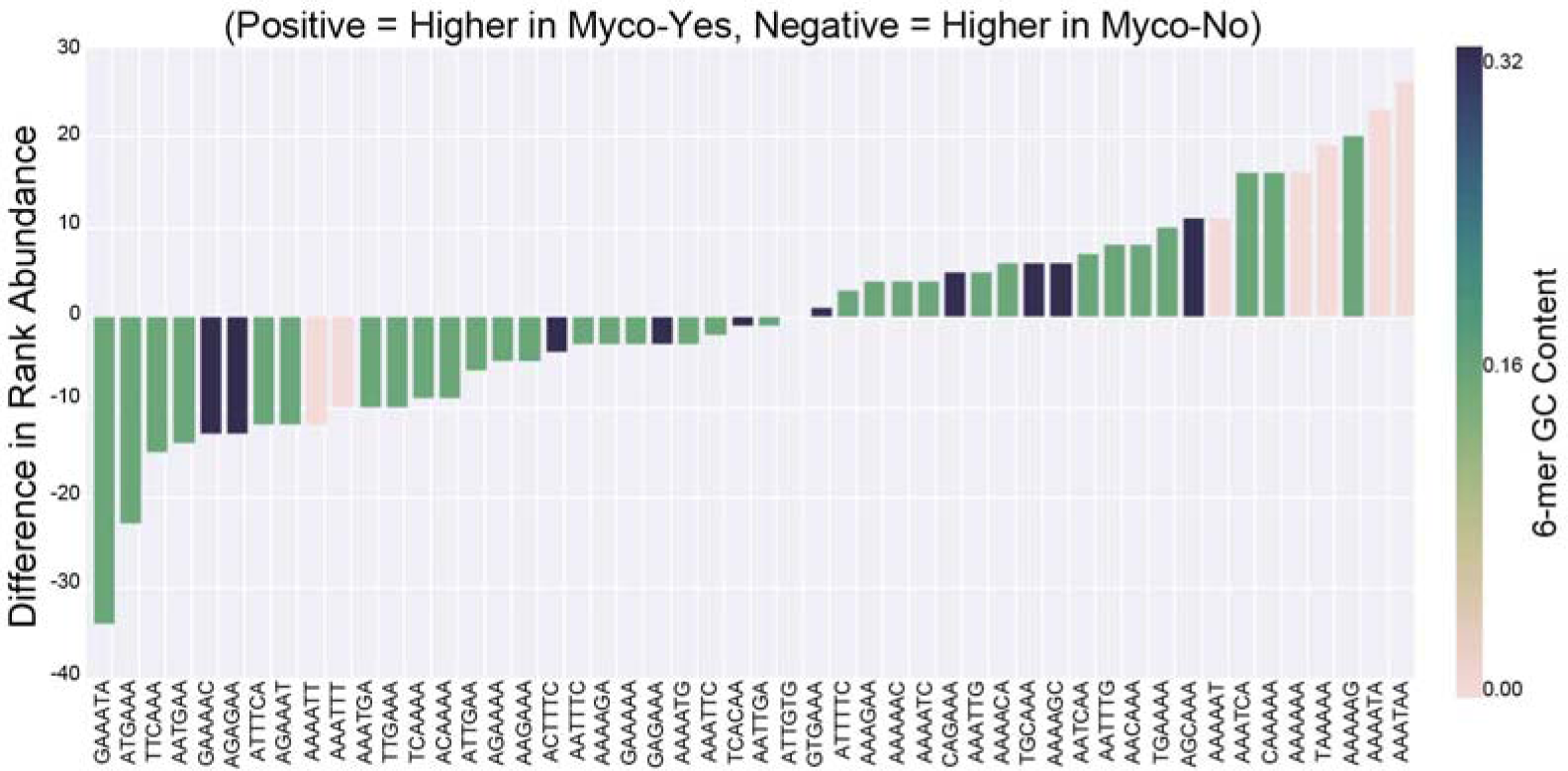
Unique 6-mers over-represented in the Myco-Yes set were characterized by very low GC content. 6-mers that were significantly over-represented in the *P. tricornutum* centromeres compared to 1) a randomly selected subset of *P. tricornutum* genomic DNA sequence, 2) randomly generated sequences at approximately 48% GC, and 3) randomly generated sequences at approximately 39% GC were identified in the Myco-Yes and Myco-No inserts sequences. After normalizing 6-mer frequencies for sequence length, the average frequency for each 6-mer per set (i.e. Myco-Yes and Myco-No) was calculated and ranked. The differences in rank between the Myco-Yes and Myco-No sets was then plotted, and the GC content of the sequence was indicated by the bar color.

## Discussion

### Features of predicted diatom centromeres

In this study, we identified native diatom centromere sequences with high resolution. Based on previous studies, we hypothesized that low GC content would be a common characteristic of diatom centromeres. We deconstructed two *Phaeodactylum tricornutum* chromosomes (25 and 26) and found that regions with low GC content appeared to function as centromeres, while adjacent regions did not. We subsequently conducted a genome-wide ChIP-seq screen (confirmed with ChIP-qPCR) and a forward genetics screen to identify centromeres and additional sequences enabling episome maintenance, and used reverse genetics to test for function. We discovered 25 unique *P. tricornutum* centromeric DNA sequences: 24 among the nuclear genome scaffolds and 1 in the non-scaffolded genome assemblies. If our results indicate a true estimate of diatom chromosomes, with one unique centromere sequence each, we would predict that the diatom genome contains fewer chromosomes than the 33 predicted previously (Bowler et al., 2008). Centromere sequences may be erroneously missing from the genome assembly. Additionally, some of the *P. tricornutum* chromosome-scale scaffolds lacking telomere-to-telomere assembly may not be individual chromosomes, but rather partial chromosomes. For example, the putative centromeres identified by ChIP-seq from chromosomes 24 and 29 were nearly identical (99%), and each of these two centromeres was positioned near a scaffold terminus lacking a telomere. Thus, chromosome-scale scaffolds 24 and 29 may be two arms of a single chromosome. In any case, the identification of centromeric DNA sequences will help to develop a better model of *P. tricornutum* genome organization. Another possibility is that the chromosomes not identified by our ChIP-seq analysis do not recruit CENP-A and may employ different mechanisms to assemble the kinetochore; however, this seems unlikely, as previously studied organisms tend to have conserved epigenetic features (i.e., CENP-A recruitment) for centromeres across all chromosomes.

Two *P. tricornutum* chromosomes, 2 and 8, appeared to deviate from the monocentric model by having two sequences identified by the ChIP-seq analysis. The regions adjacent to the centromeres on the chromosome scaffolds are unresolved DNA sequence and both centromere regions contained long direct repeats. Thus, sample processing, sequencing, or assembly error could be responsible for the apparent duplication of the centromere on these chromosomes. Attempts to resolve the structure of the region using PCR were unsuccessful (data not shown). Alternatively, these may be true centromeres that have simply been duplicated. The presence of a nearby retrotransposon may support this theory, and could also confound PCR assays (Figure 3). Dicentric chromosomes have been noted in several organisms; however, typically only one of the centromeres is active and the other is inactivated (Cuacos et al., 2015; Neumann et al., 2012; Sato et al., 2012; Stimpson et al., 2012; Sullivan et al., 2001). The presence of two active centromeres typically leads to chromosomal breakage followed by either cell death or two functional monocentric chromosomes. Chromosomes with multiple functional centromeres have been identified. In human cells two active centromeres were in close proximity, essentially behaving as a single centromere (Sullivan and Willard, 1998). In rice, recombinant centromeres were found to contain 2 repetitive arrays; both recruited CENP-A, while an intervening sequence did not (Wang et al., 2013). Additionally, tricentric chromosomes were identified in wheat where one of the centromeres was large and presumably dominant, and co-occurring centromeres were smaller and weaker (Zhang et al., 2010).

ChIP-seq peaks were typically found only once per chromosome suggesting *P. tricornutum* has small monocentric regional centromeres. This centromere structure is also found in the closest related organisms with identified centromeres: the protist *P. falciparum* and the red alga *Cyanidioschyzon merolae* (Bowman et al., 1999; Iwanaga et al., 2010; Kanesaki et al., 2015; Maruyama et al., 2008). Both organisms have similarly sized centromeric DNA regions (˜2-4 kb) and also share low GC content as a characteristic of their centromeres: ˜3% relative to the genome average GC of 21.8% in the case of *P. falciparum* (Bowman et al., 1999; Iwanaga et al., 2010), and 48.4% relative to the genome average GC of 55% for *C. merolae* (Kanesaki et al., 2015). Interestingly, *C. merolae*, which is the only other alga with well characterized centromeres and the closest relation to *P. tricornutum* of the organisms studied, has centromeres with a GC content that is low compared to the genome average, similar to *P. tricornutum.*

Like *P. tricornutum*, the diatom *Thalassiosira pseudonana* can also utilize the yeast-derived *CEN6-ARSH4-HIS3* sequence to maintain episomes (Karas et al., 2015), which may suggest an overall similarity in DNA maintenance mechanisms. We analyzed the GC content of the *T. pseudonana* genome and found similar regions of low GC content that were often found once per chromosome-scale scaffold (Supplementary Figure 6). Thus, the ability of the yeast *CEN6-ARSH4-HIS3* sequence to support episomal maintenance in both species may be due to similar requirements for low GC sequences to function as centromeres. It is remarkable that these diatoms may have such similar centromere features, to the degree that the same sequence can function as a centromere in both organisms, given the ancient evolutionary divergence of the centric and pennate diatom lineages (˜90 Myr ago, (Bowler et al., 2008)) and the relatively rapid evolution of centromere sequences and structures observed for other groups of organisms (Malik and Henikoff, 2002, 2009). Further CENP-A ChIP-seq experiments in *T. pseudonana* will enable centromere identification and comparison to *P. tricornutum*, including an examination of evolutionary implications.

### Simple centromere requirements permit nuclear maintenance of non-nuclear DNA sequences

In this study, by identifying characteristics of native diatom centromere sequences, we have uncovered a mechanism by which foreign DNA can become part of the nuclear DNA repertoire; non-nuclear DNA can act as a centromere, enabling stable maintenance as an extrachromosomal nuclear episome. Maintaining such plasmids may expand the diatom’s genetic potential, and may also facilitate permanent integration into the nuclear chromosomes. We previously observed that DNA sequences from the yeast *Saccharomyces cerevisiae* could enable episome maintenance in *P. tricornutum* (Diner et al., 2016; Karas et al., 2015), and in this study, we confirmed that this sequence does, in fact, recruit the *P. tricornutum* centromeric histone protein CENP-A. The recruitment of this centromere-specific histone protein and subsequent maintenance of the episome in diatoms suggests the foreign DNA sequence is using native diatom DNA replication machinery, essentially functioning as a diatom centromere. There are very few examples in eukaryotes of foreign DNA recruiting host CENP-A to maintain a chromosome. Human centromeres have previously been shown to function in mouse chromosomes (Hadlaczky et al., 1991), and in a recent example, *Arabidopsis* centromeric repeats were shown to recruit human CENP-A and maintain chromosomes in human cells (Wada et al., 2016). In both cases, the chromosomes maintained by foreign DNA originally derived from chimeric host-donor DNA chromosomes followed by chromosomal breakage and/or rearrangement, resulting in smaller linear chromosomes or “mini-chromosomes.” To our knowledge, there are no examples of immediate nuclear genome establishment (i.e., without chimeric intermediates) and maintenance in the host cell as a plasmid. This contrasts with bacteria, where DNA transfer between bacterial and subsequent plasmid establishment is quite common. Our results suggest that non-nuclear DNA can mimic diatom centromeres and, along with co-localized DNA, can immediately establish circular chromosomes in the diatom genome.

Establishment of centromeres in *P. tricornutum* is apparently governed by the simple rule of having a small length of sequence with a GC content less than ˜33%. Although ChIP-seq peaks for centromeres averaged 39% GC over the entire 2-5 kb sequence, each centromeric ChIP-seq peak contained within it a 500-bp region less than ˜33% GC. While 500-bp is the shortest sequence we tested, it is possible that even smaller sequences could be sufficient to function as centromeres. Notably, 500-bp is a particularly short sequence to enable centromere function compared to previously studied organisms with regional centromeres; most regional centromeres are reported to be thousands of basepairs in length, compared to the relatively small (˜125 bp) point centromeres of some yeast species. The low GC rule we propose here for *P. tricornutum* centromere establishment may also apply to diatom genes not originating in the nucleus, which in terms of diatom evolution represent a major source of diatom nuclear DNA. *P. tricornutum* chloroplast and mitochondrial genomes are low in GC content (32% average GC for the chloroplast, 35% average GC for the mitochondria), and we identified multiple sequences from each that could maintain episomes (Supplementary Figure 3).

Despite the experimental support for the simple average GC rule described above, three *Mycoplasma* sequences with GC less than 33% did not support episomal maintenance. When we examined the frequency of consecutive A+T nucleotides in the sequences, the 30% GC and 28% GC sequences that failed to support episomes had lower frequencies of sequences of 6 or more consecutive A+T bases. This lower frequency of contiguous A+T stretches was also apparent when comparing the set that failed to support episomes to a randomly generated set of 30% GC sequences. Thus, the true signal for centromere establishment in *P. tricornutum* may be the frequency or spacing of longer contiguous A+T sequences, and sequences of <33% GC content usually, but not always, happen to contain these signals. This hypothesis remains to be tested with additional sequences.

Most algal genomes studied to date contain large amounts of foreign DNA in their nuclear genomes, including recently acquired bacterial and viral DNA, transposable elements, and DNA acquired from endosymbionts (e.g., genes originating from chloroplast and mitochondrial genomes or secondary endosymbioses in the case of the Stramenopile algae) (Archibald, 2009). Thus, understanding mechanisms facilitating nuclear gene acquisition can shed light on algal diversity and evolution. For diatoms in particular, lateral gene transfer has played a major evolutionary role on multiple scales (Armbrust, 2009). Diatoms, like other stramenopiles, are the result of serial endosymbiotic events, though the precise details are still debated. Originally, a basal eukaryote engulfed and enslaved a cyanobacterium as a plastid, transferring much of the cyanobacterial genome to the nucleus (Martin, 2003). Later in evolutionary history, a non-photosynthetic eukaryote engulfed and enslaved a photosynthetic eukaryote. Endosymbiotic gene transfer from both the plastid and nuclear genomes of the secondary endosymbiotic event form a large portion of the current diatom nuclear genome. In both these events, the DNA transferred to the recipient nucleus was already inside the host cell. More recent gene transfers from bacteria are also apparent from genome analyses (Bowler et al., 2008), and the recent discovery that diatoms are amenable to bacterial conjugation (Karas et al., 2015) provides a potential mechanism for exogenous DNA transfer. Foreign DNA can also enter diatom cells through viral infection. In all these events, the mechanism by which the endosymbiotic or exogenous DNA becomes incorporated into the nuclear genome is unknown.

It is unclear why maintenance of foreign DNA in the form of episomes appears to be well tolerated in *P. tricornutum.* One possibility is that transfer of foreign DNA into diatoms, or intracellular transfer of previously acquired non-nuclear genetic material, is not common enough for a defense system to have evolved (such as the production of restriction enzymes in bacteria to destroy foreign DNA). In contrast to bacteria-bacteria DNA transfer, non-native genes are unlikely to be expressed from a plasmid transferred to a diatom if they are of bacterial origin. Functional gene expression would only occur in the unlikely event that it acquired diatom transcriptional, translational, and subcellular localization signals through further modification. Thus, it is possible there was not strong selection to evolve defense mechanisms against foreign DNA because they were not detrimental to cell fitness and most events were entirely innocuous. If such permissiveness occurs for maintenance of DNA transferred through extracellular mechanisms, it is likely that it would also apply to DNA transferred to the nucleus intracellularly from organelles to the nucleus.

## Conclusions

Identifying and characterizing centromeres is essential for understanding cellular biology, as these are critical features for stable DNA maintenance during cell division. These sequences can also advance synthetic biology through the development of new molecular tools, including artificial chromosome optimization. Here, we have used multiple approaches to characterize the centromeres of the diatom *P. tricornutum.* We found very simple sequence requirements for DNA to function as a centromere, namely a moderately low GC content of <33% across a small region, with 500-bp being that smallest size examined. While most sequences with GC content of <33% allowed episomal maintenance, a few sequences with this simple characteristic did not, and we predicted that a higher frequency of stretches of contiguous A+T bases may be as important as overall average GC content of a fragment in establishing a centromere and supporting an episome. Based on bioinformatic analyses, we predict that these features of centromere identity may be conserved in the distantly related diatom *Thalassiosira pseudonana.* While low GC content has often been identified as a centromeric DNA feature, the diatom centromeres appear to be unique from many other eukaryotes in that they are not composed of repeat regions or other notable primary structures, and that the functional centromere region may be quite small. We also show that these simple requirements mean foreign and non-nuclear DNA sequences with these characteristics can act as centromeres in diatoms, becoming established as extrachromosomal nuclear episomes. Diatoms possess nuclear genes acquired from many foreign DNA sources including viruses, bacteria, and other eukaryotes, which includes the ancient endosymbiotic acquisition of mitochondria and chloroplasts. Our findings present a host-permissive mechanism by which DNA derived from either external or intracellular genetic compartments can become a part of the nuclear DNA repertoire by utilizing host replication and maintenance machinery. This may ultimately lead to gene integration into diatom genomes and subsequent evolutionary diversification.

## Materials and Methods

### Strains and culturing conditions

*Phaeodactylum tricornutum* strain CCMP 632 cultures were grown in L1 artificial seawater medium (Price et al., 1989) or on ½xL1-agar plates at 18°C under cool white fluorescent lights (50 μE m^−2^ s^−1^) as previously described (Diner et al., 2016; Karas et al., 2015). Ex-conjugants were selected on phleomycin (20 μg mL^−1^). *Escherichia coli* (Epi300, Epicentre, WI, USA) were grown in LB broth or agar supplemented with ampicillin (100 μg mL^−1^), tetracycline (10 μg mL^−1^), chloramphenicol (20 μg mL^−1^), gentamicin (20 μg mL^−1^), or combinations of these as needed.

DNA was obtained from multiple organisms to test sequences for episome maintenance, including *P. tricornutum*, Synthetic *Mycoplasma mycoides* JCVI-syn1.0 (NCBI accession #CP002027), and *Altermonas macleodii* strain U4 containing the pAMDE1 plasmid (NCBI accession #CP004849)(López-Pérez et al., 2013). Because templates for PCR amplification of *Cylindrotheca fusiformis* plasmids pCF1 (NCBI accession #X64302) and pCF2 (NCBI accession #X64303) were not available, we synthesized the DNA sequences by assembling 18 individual oligonucleotides (CF1-1 through 18 and CF2-1 through 18, respectively) using Gibson assembly (Supplementary Table 8)(Gibson et al., 2009; Hildebrand et al., 1992).

### Construction of plasmid pPtPBR1-YFP-CENP-A for native centromere identification

The plasmid pPtPBR1-YFP-CENP-A was constructed to express the CENP-A protein in *P. tricornutum*, tagged with a fluorescent protein for use in cellular localization and ChIP-seq analyses. The plasmid was constructed by Gibson assembly with a PBR322 backbone, and contains an N-terminal YFP fused to *P. tricornutum* CENP-A (Phatr2_16843 but with the wrong start site. See Supplementary Figure 7 for complete sequence, translation, and genome coordinates of *P. tricornutum* CENP-A used for cloning), a design modeled after prior *Arabidopsis thaliana* YFP-CENP-A fusions (Ravi and Chan, 2010) (Supplemental Figure 8).

First, ShBle cassette was added to the pUC19 cassette. Plasmid pUC19 was digested with EcoRI and SmaI and the ShBle Cassette from pAF6 (Falciatore et al., 1999)was amplified using primers Puc-ShBle-F and Puc-ShBle-R. The resulting plasmid, pUC19-ShBle, was digested with HindIII and BamHI and the FcpB promoter (amplified with primers Puc-PromTerm-1 and Puc-PromTerm-2) and FcpA terminator (amplified with primers Puc-PromTerm-3 and Puc-PromTerm-4) regions were assembled in three fragments by Gibson assembly. The resulting plasmid, pUC19-ShBle-PromTerm, contains an AgeI restriction site between the promoter and terminator regions. Next, plasmid pUC19-ShBle-PromTerm was digested with AgeI and it was assembled with YFP (CENPA-YFP-3 + CENPA-YFP-4) and CENPA (CENPA-YFP-5 + CENPA-YFP-8) PCR products to create plasmid pUC19-ShBle-YFP-CENPA. Plasmid pUC19-ShBle-YFP-CENPA contains an N-terminal YFP fused to *P. tricornutum* CENPA and the two domains are separated by a linker consisting of 5 glycines and 1 alanine. Fusion of YFP to the N-terminus CENP-A was modeled after *Arabidopsis* fusions (Ravi and Chan, 2010). The insert FcpBprom-YFP-CENPA-FcpAterm was amplified from Puc19-ShBle-YFP-CENPA using primers PtPBRYFP-CENPA-1 and PtPBRYFP-CENPA-2 and assembled into pPtPBR1 which was amplified in two pieces using primers PtPBRnanoluc5 and PtPBRrev for piece 1 and PtPBRnanoluc4 and PtPBRfor for piece 2. The result of the three-piece assembly was named pPtPBR-YFP-CENPA. The sequence of the YFP-CENP-A insert was verified by Sanger DNA sequencing, and the plasmid was transformed into *E. coli* strain Epi300 containing the plasmid pTA-MOB and mobilized into *P. tricornutum* by conjugation. *P. tricornutum* cells containing plasmid pPtPBR1-YFP-CENP-A were grown in liquid culture to a density of 3 x 10^6^ cells mL^−1^. Cells were harvested simultaneously for microscopy, Western blot, and ChIP (Supplementary Figure 8).!

### Western Blot confirmation of YFP-CENP-A fusion protein expression and microscopy localization

For the Western Blot, 50 mL cultures of late log phase (1 × 10^6^ cells mL^−1^) *P. tricornutum* cells expressing plasmid pPtPBR1-YFP-CENP-A were centrifuged for 10 minutes at 4,000g and resuspended in 1x NuPAGE LDS sample buffer (Life Technologies) containing 50 mM dithiothreitol. Resuspended cells were divided into 300 μL aliquots and sonicated (15 minutes: 1 minute high power, 30 seconds off) in a Diagenode Bioruptor cooled to 4°C. Lysates were cleared by centrifugation at 16,000g for 10 minutes at 4°C, and were stored at −20°C until use. Lysates were diluted approximately four-fold, heated for 5 minutes at 95°C, and centrifuged for 2 minutes at 16,000g to remove precipitates. Proteins were separated by SDS-PAGE on NuPAGE Novex 12% Bis-Tris polyacrylamide gels (Life Technologies) in 1x MOPS running buffer (Life Technologies) for 1 hour at 150 V. Separated proteins were transferred by electroblotting to nitrocellulose membranes for 1 hour at 100 V in 1x Transfer Buffer (Life Technologies) containing 10% methanol. Blots were incubated in blocking solution for 30 minutes at room temperature (WesternBreeze Chemiluminescent Kit, Life Technologies) and incubated with primary antibody (anti-GFP, AB290 from AbCam) diluted 1:500 in prepared blocking solution overnight at room temperature. Blots were then washed 4 times with 1x TBST for 5 minutes per wash and incubated with goat anti-rabbit secondary solution (WesternBreeze Chemiluminescent Kit, Life Technologies). Blots were washed 4 times with 1x TBST for 5 min per wash and incubated with CDP-Star chemiluminescent substrate and imaged with the C-digit (LiCor, NE, USA).

To visualize the expression of the YFP-labeled CENP-A protein in *P. tricornutum*, live ex-conjugant cells were imaged on a Zeiss (Oberkochen, Germany) axioscope epifluorescent microscope with a YFP filter. Strains positive for the YFP construct were washed twice in PBS and incubated with the nuclear counter-stain DAPI (4’,6-diamidino-2-phenylindole), 300 nM, for three minutes. Cells were re-washed, oil mounted and visualized on a Leica (Wetzlar, Germany) TCS SP5 confocal laser scanning microscope using a 405 nm diode laser (emission 440–470 nM) for DAPI and a 514 nm argon laser (emission 530 – 570 nm) for the YFP, while plastid autofluorescence was monitored at 700–740 nm (Supplementary Figure 8).

### Chromatin immunoprecipitation (ChIP) sequencing and qPCR

For ChIP, 2-4 × 10^8^ cells were pelleted. Fixation, extraction, and sonication were performed as previously described (Lin et al., 2012). Immunoprecipitations were performed using the OneDay ChIP kit from Diagenode following the manufacturer’s instructions. For small-scale qPCR experiments 66 μL of sheared chromatin was used, and for large-scale sequencing experiments 2,300 μL of chromatin was used with proportional increases in reagents. ChIP-grade Anti-GFP (AB290, Abcam MA, USA) was used at 1 μL per reaction for small-scale and 30 μL for large-scale; controls omitting antibody during immunoprecipitation were also performed.

For ChIP-seq, libraries for the Illumina sequencing platform were prepared from the large-scale ChIP reactions. ChIP samples were concentrated by vacuum centrifugation followed by purification on a Qiagen PCR cleanup column. Libraries for input DNA, anti-GFP antibody, and no antibody control samples were prepared using the NEBNext Ultra II kit (New England Biosciences, MA, USA). After end preparation, adapter ligation, and cleanup, the libraries were barcoded using 5 cycles of PCR for the input library and 12 cycles of PCR for the ChIP samples performed with antibody and without antibody. After final cleanup with SPRI beads, the samples were sequenced on an Illumina MiSeq DNA sequencer. Approximately 4 million paired end reads were obtained for the input library while 1 million paired ends reads were obtained for each of the ChIP reactions performed with and without antibody. Sequences were mapped onto the *P. tricornutum* genome using CLC. We performed qPCR as described in Karas et al. 2015. Primer sets were designed to amplify regions on the episome including the ARSH4 region which was associated with a ChIP-seq peak (primers: Q-ARS-1 + Q-ARS-2) and the TetR region as a negative control (primers: Q-TETR-1 + Q-TETR-2), which were separated by 3.4 kb. Regions on native *P. tricornutum* chromosome 25 were amplified by primers Q-25HR-1 + Q-25HR-2 for the region that recruited a ChIP-seq peak, and by primers Q-25control-1 and Q-25control-2 for a region with that did not. Product sizes for all qPCR reactions were 100-200 bp. For ChIP-qPCR, Ct values were obtained for input, no-antibody control, and anti-YFP reactions. Data was analyzed as previously described (Lin et al., 2012). First, input Ct values were adjusted to account for dilution. Then ΔCt were calculated by subtracting the corrected input values from the Ct values from no-antibody control and anti-YFP samples (ΔCt = Ct_sample_ – Ct_correctedinput_). Percent input was calculated as 100/(2^ΔCt^).

### Bioinformatic Analyses

Bioinformatic analyses were conducted to map peak reads to the *P. tricornutum* genome assembly, to correlate the GC content of *P. tricornutum* DNA regions with CENP-A ChIP-seq peaks, and to identify sequence features unique to centromere and centromere-like maintenance sequences. The genome was first broken down into various sized windows from 10-kb to 0.5-kb with 20% sequence overlap with each step. We then calculated the GC content of *P. tricornutum* genomic DNA sequence in 100-bp windows that advanced by 50-bp each step. To identify small regions (1-5 kb) with higher densities of low GC sequences, we counted the number of 100-bp windows with GC below a given threshold. We tested regions from 2 kb to 5 kb (advancing 1 kb each step) and GC thresholds less than or equal to values 30-33. Sequences predicted to be centromeres by ChIP-seq and further confirmed by GC analysis were examined for sequence similarity by BLAST. The set of 29 putative centromere sequences on 27 scaffolds was searched against a database constructed from the set itself using blastn with default settings (BLAST v2.2.29). Sequence motifs among the putative centromere sequences were searched using MEME (Bailey et al., 2009), and the presence of repeated sequence in the centromeres was searched with Tandem Repeat Finder (Benson, 1999). The 500-bp regions containing the lowest GC content were tested for centromere function in downstream analysis.

Double-stranded K-mer frequency matrices were generated using sliding window of size K through each sequence. A window size (K) of 6 was chosen as it was the longest string that was still well-represented in a sequence of 500-bp. At each window step, the 6-mer was counted along with its reverse complement. 6-mer count vectors were normalized by the total 6-mers in the sequence to weight sequences of varying length equally. To avoid double-counting of 6-mers and to allow for arbitrary orientation of the centromere sequence, each 6-mer was represented by a forward and reverse complement sequence, and only a single representation was considered in downstream analysis. Each 6-mer at this stage encoded information for both strands and the frequency was multiplied by 2 so each 6-mer vector summed to 1.

Genomic control sequences were selected *in silico* by slicing the *P. tricornutum* genomic DNA sequence at random positions. The number and length of the genomic control sequences mirror the number and length of the centromeric sequences. Null sets of random sequences were generated with a target GC of 47% (similar to non-centromere average) and 39% (similar to centromere ChIP-seq peak average). P-values were calculated using a Wilcoxon signed-rank test for each 6-mer between the centromere group and each of the control groups. If the distribution of a 6-mer was significantly different (P < 0.001) when tested against genomic and null control groups, then it was used in the subsequent downstream analysis.

The frequency of contiguous A+T sequences was calculated as follows. The DNA sequences were recoded using IUPAC ambiguity codes to represent {G,C} as S (Strong) and {A,T} as W (Weak). Consecutive W sequences were counted by choosing a size K and identifying every position in which a window composed entirely of W nucleotides of size K was found in the sequence; counts were normalized by total K-mers in the sequence (i.e., dividing counts by length(sequence) - K + 1). Computational analysis was employed in Python 3.5.2 using the Continuum Analytics Scientific Computing ecosystem for data processing (NumPy v1.11, SciPy v0.18.1, and Pandas 0.19.1), data visualization (Matplotlib v1.5.1 and Seaborn v0.7.1), and sequence handling (Scikit-Bio v0.5.1).

### *P. tricornutum* genomic library construction

Genomic DNA from *P. tricornutum* was prepared using the DNeasy Plant Mini kit (Qiagen, Hilden, Germany) using approximately 100 mL of 1 × 10^6^ cells mL^−1^. Cells were pelleted and resuspended in 400 μL AP1, and DNA was extracted following the manufacturer’s instructions. Fourteen μg DNA in 200 μL TE buffer were sonicated using a Covaris (MA, USA) red miniTUBE to generate fragments centered around 5 kb. Fifty μL was then used for end repair and adapter ligation with the NEB Next Ultra II library kit. Adapter ligated fragments were amplified using Index primer 1 and the Universal primer for 8 cycles. After purification with SPRI beads (Agencourt AMPure XP, Beckman Coulter, CA, USA), fragments were assembled into the vector backbone pPtPBR2, which was PCR amplified using primers PtPBR2-Illumina-F and PtPBR2-Illumina-R (Supplementary Table 8). The assembly reaction was transformed into pTA-MOB-containing *E. coli* cells and colonies were screened for the presence of *P. tricornutum* genomic DNA inserts using flanking primers CAHtest1+InsertR (Supplementary Table 4). The library ultimately consisted of ˜6000 *E. coli* colonies containing 2- 5 kb genomic DNA inserts, with an estimated coverage of ˜0.7x of the *P. tricornutum* genome. Genome size per cell was estimated at 43 Mb, including 27.4 Mb for the nuclear genome, ˜100x of the 0.118 Mb chloroplast genome, and ˜50x of the 0.077 Mb mitochondrion genome. Chloroplast and mitochondrion genome copy numbers were based on prior calculations (Karas et al., 2015). The library was stored and conjugated into *P. tricornutum* in 9 equal pools of approximately 600-700 *E. coli* colonies to preserve diversity. The resulting 20-100 *P. tricornutum* colonies for each pool were patched onto L1 plates containing phelomycin 20 μg mL^−1^ and chloramphenicol 20 μg mL^−1^. After sufficient growth, plates containing patched *P. tricornutum* colonies were flooded with L1 medium, scraped, and pooled, at which point episomes were extracted as previously described (Karas et al., 2015). Transformation of the extracted episomes to *E. coli* via electroporation yielded episomes that had been successfully maintained in *P. tricornutum.* Colony PCR reactions to amplify the inserts were performed using primers CAHtest1 and InsertR (Supplementary Table 4). PCR products were purified followed by Sanger DNA sequencing performed by Genewiz, Inc.

### Plasmid construction, conjugative DNA transfer, and episome maintenance confirmation

Plasmids were constructed to examine the ability of the foreign and endogenous DNA sequences to maintain DNA as episomal vectors in diatoms. All primers used for insert amplification and assembly, along with template sequence coordinates, are listed in Supplementary Tables 8 and 9, along with additional assembly details.

To test large regions of *P. tricornutum* DNA, we used overlapping 100-kb regions from chromosomes 25 and 26 that were previously cloned in in a modified pCC1BAC vector (conferring *E. coli* chloramphenicol resistance) as described previously (Karas et al., 2013). These plasmids contained the *CEN6-ARSH4-HIS3* (CAH) fragment as part of the yeast-based TAR cloning procedure. Plasmids were transferred to *E. coli*, and lambda red recombineering (Datsenko and Wanner, 2000) was used to replace the CAH with a kanamycin resistance gene. The kanamycin resistance cassette was amplified by PCR from the plasmid pACYC177 using primers KO-CAH-1 and KO-CAH-2. The resulting PCR product contained 40-bp homology at the 5’ and 3’ ends to regions flanking the CAH cassette in the vector backbone. The PCR product was purified, adjusted to 100-200 ng μΓ^1^, and introduced by electroporation into *E. coli* cells that contained both the cloned 100-kb fragments and lambda red plasmid pKD46 according to the recommended procedure (Datsenko and Wanner, 2000). Replacement of the CAH region by the kanamycin resistance cassette was confirmed by PCR on the resulting *E. coli* colonies that were resistant to kanamycin and chloramphenicol.

For all other DNA sequences tested for episomal maintenance in this study, the vector pPtPBR2, which is incapable of episomal maintenance in diatoms, was used as a backbone (Diner et al., 2016)(Supplementary Table 9). DNA vector inserts were PCR amplified and confirmed by gel electrophoresis. Vectors were assembled using Gibson assembly, and screened using restriction digest. In addition to the 100-kb *P. tricornutum* DNA regions described above, several smaller regions of chromosomes 25 and 26 were examined for episome maintenance ability including three 10-kb regions (Pt-10kb-6, Pt-10kb-9. Pt-10kb-12) and two 1 kb regions (Pt-1kb-25, Pt-1kb-26). Based on the results of the ChIP-seq analysis an additional 38 constructs (Test1-40) were created to confirm the functionality of the predicted centromere sequences.

Foreign DNA inserts of various insert sizes (500-, 700-, and 1000-bp) and GC contents (15, 21, 28, 30, 40, and 50%) were obtained from the *M. mycoides* JCVI-syn1.0 genome. In total, 24 *M. mycoides-based* insert sequences were tested. Two insert sequences containing ˜500 bp of *A. macleodii* pAMDE1 plasmid DNA were tested for episome maintenance, which contained ˜26 and 29% GC content each. To examine maintenance ability of *C. fusiformis* plasmid DNA, one insert sequence of ˜500-bp was synthesized from each of the two CF plasmids, pCF1 and pCF2 (Supplementary Table 5).

DNA was transferred by conjugation from *E. coli* into *P. tricornutum* following previously described methods (Diner et al., 2016; Karas et al., 2015). Each experiment was run alongside positive (pPtPBR1) and negative (pPtPBR2) control plasmids, and efficiency of conjugation was determined by comparing the number of ex-conjugant colonies obtained after conjugation with a plasmid to the number of ex-conjugant colonies obtained from the negative control plasmid (pPtPBR2) performed simultaneously. Ex-conjugant diatom lines representing each foreign and endogenous DNA source were further examined for the presence of the episome in *P. tricornutum* (as opposed to selective marker genomic integration) and maintenance. These lines were passaged on selection for ˜1 month (transferred weekly) to avoid plasmid carryover, and the absence of *E. coli* donors was confirmed by plating on LB agar medium. Plasmids were isolated using the modified alkaline lysis protocol described in Karas et al. 2015, and were subsequently transformed into *E. coli* by electroporation to confirm plasmid presence. Lines were then passaged semi-continuously in liquid media containing antibiotics for ˜30 days before plasmid extraction, *E. coli* transformation, and confirmation of plasmid presence and size using restriction digest (Diner et al., 2016; Karas et al., 2015).

## Acknowledgements

We thank John McCrow for many helpful discussions concerning bioinformatics and Sarah Smith for assistance with ChIP-seq. Funding for this work was provided by the Gordon and Betty Moore Foundation (GBMF5007 to P. D. W. and C. L. D., GBMF3828 and GBMF5006 to A. E. A.), the U.S. Department of Energy (DE-SC0008593) to A. E. A. and C. L. D., and the National Science Foundation (NSF-MCB-1129303 to C. L. D. and OCE-1136477 to A. E. A.).

---

**Supplementary Figure 1:**
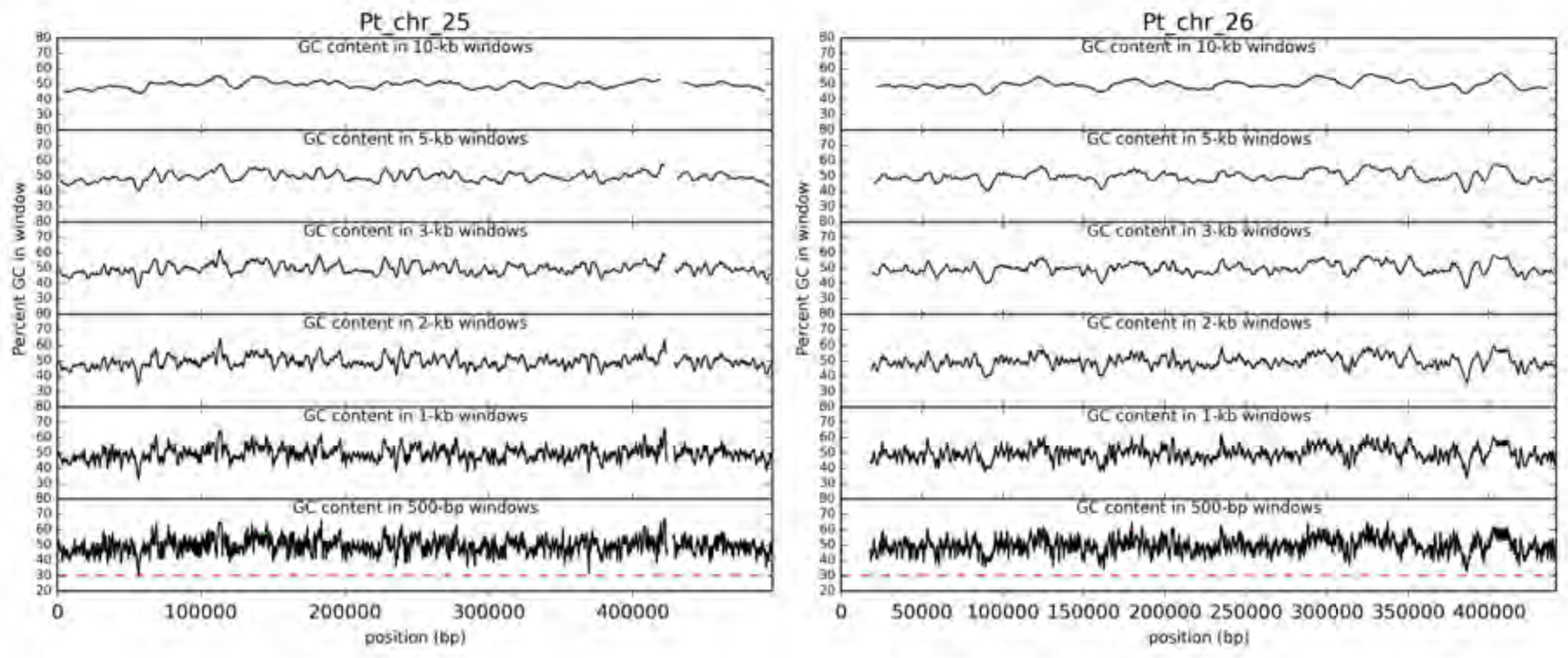
GC content of *P. tricornutum* chromosome 25 and 26 scaffolds using sliding window sizes of 10-, 5-, 3-, 2-, 1-, and 0.5-kb. Breaks in the lines indicate the presence of unknown sequence within the window. A reference dashed line at 30% is drawn for the 0.5-kb plots.

**Supplementary Figure 2:**
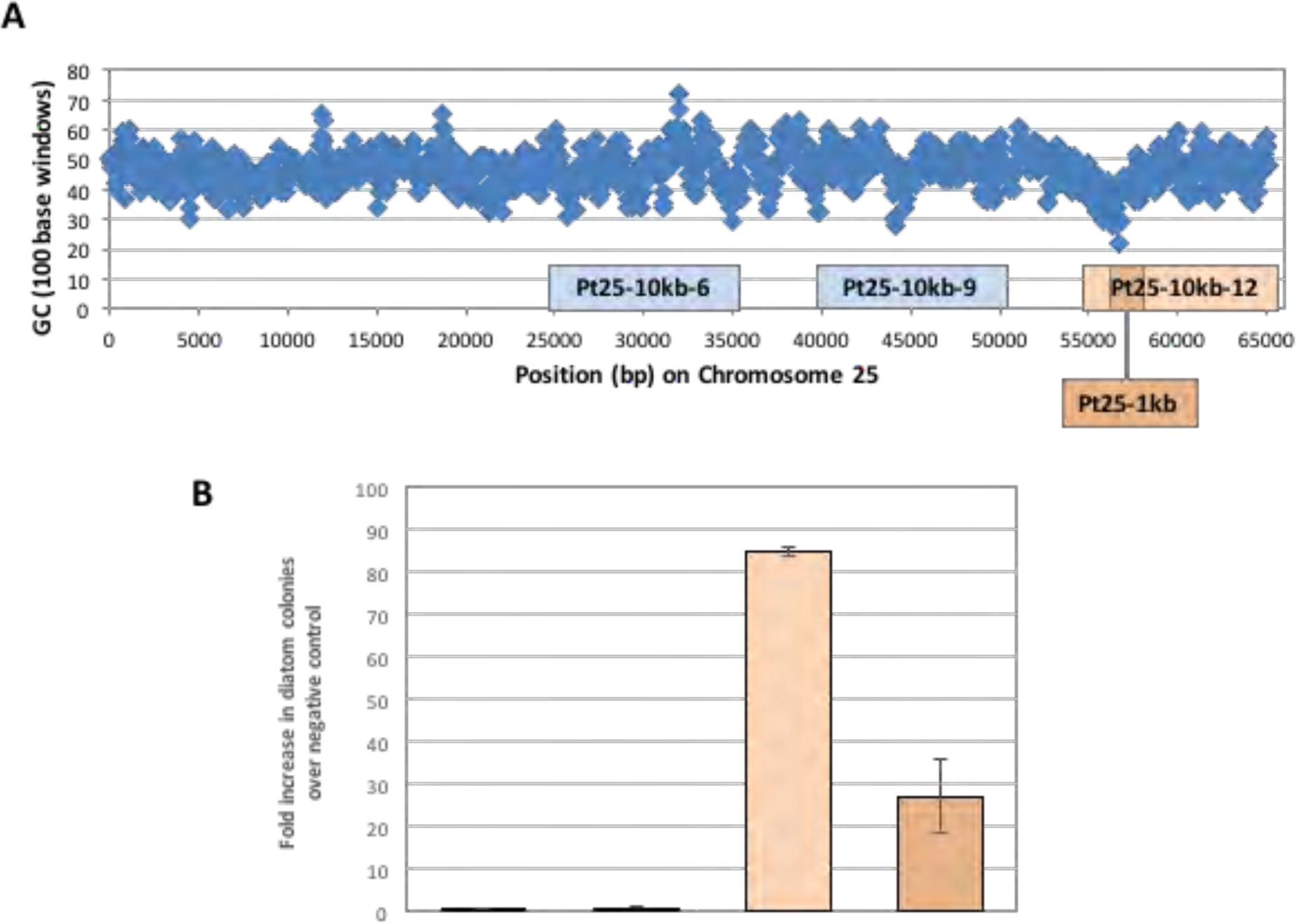
Regions of *Phaeodactylum tricornutum* chromosome 25 that support episomal maintenance. **A.** Enlargement of chromosome 25 from 1-65,000 bp showing GC content in 100-bp windows overlapping by 50-bp, overlayed with the *P. tricornutum* DNA inserts tested (Pt25-10kb-6, Pt25-10kb-9, Pt25-10kb-12, and Pt25-1kb). **B.** Number of diatom ex-conjugant colonies obtained after conjugation, shown as fold increase in colony numbers over the pPtPBR2 negative control.

**Supplementary Figure 3:**
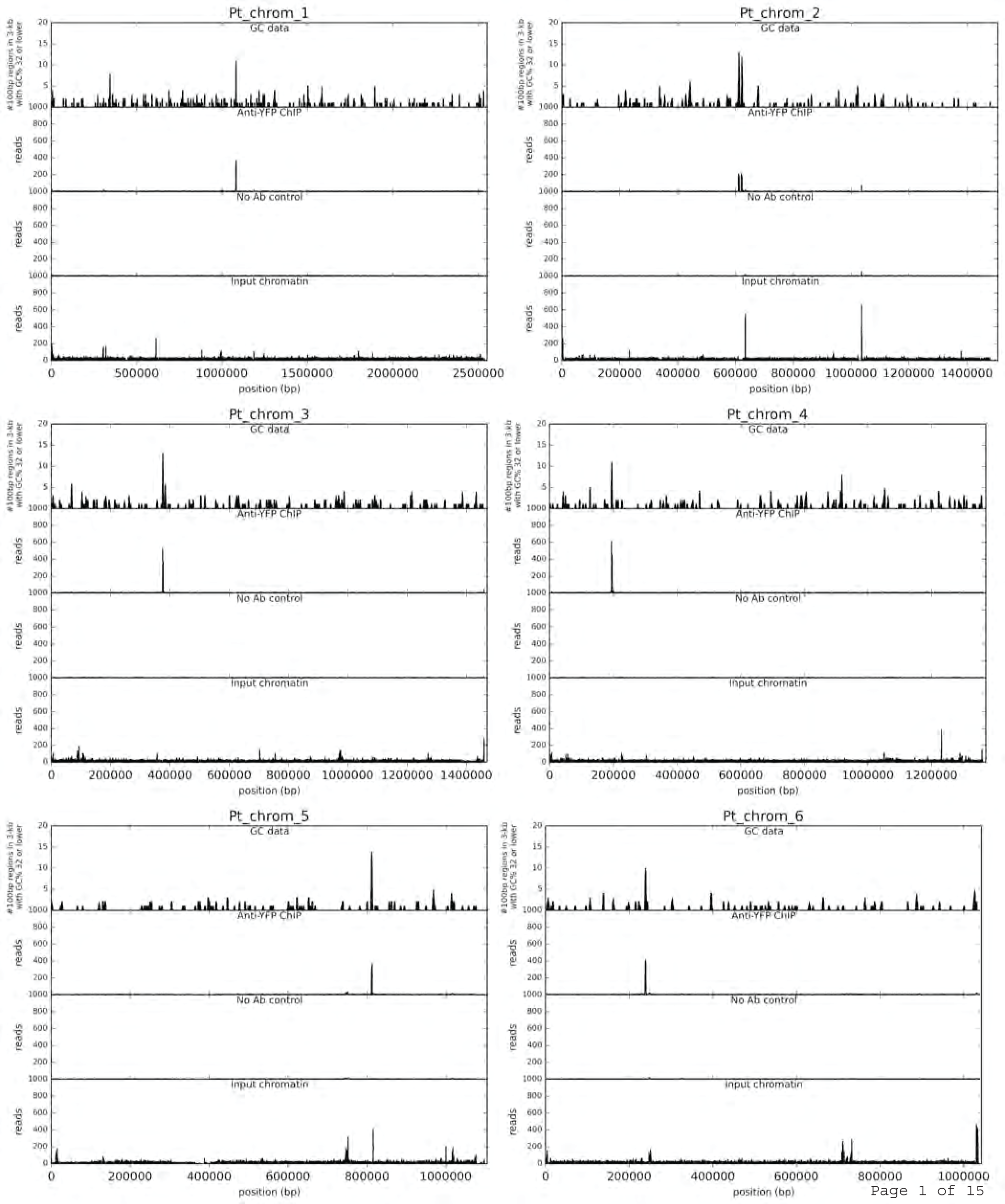
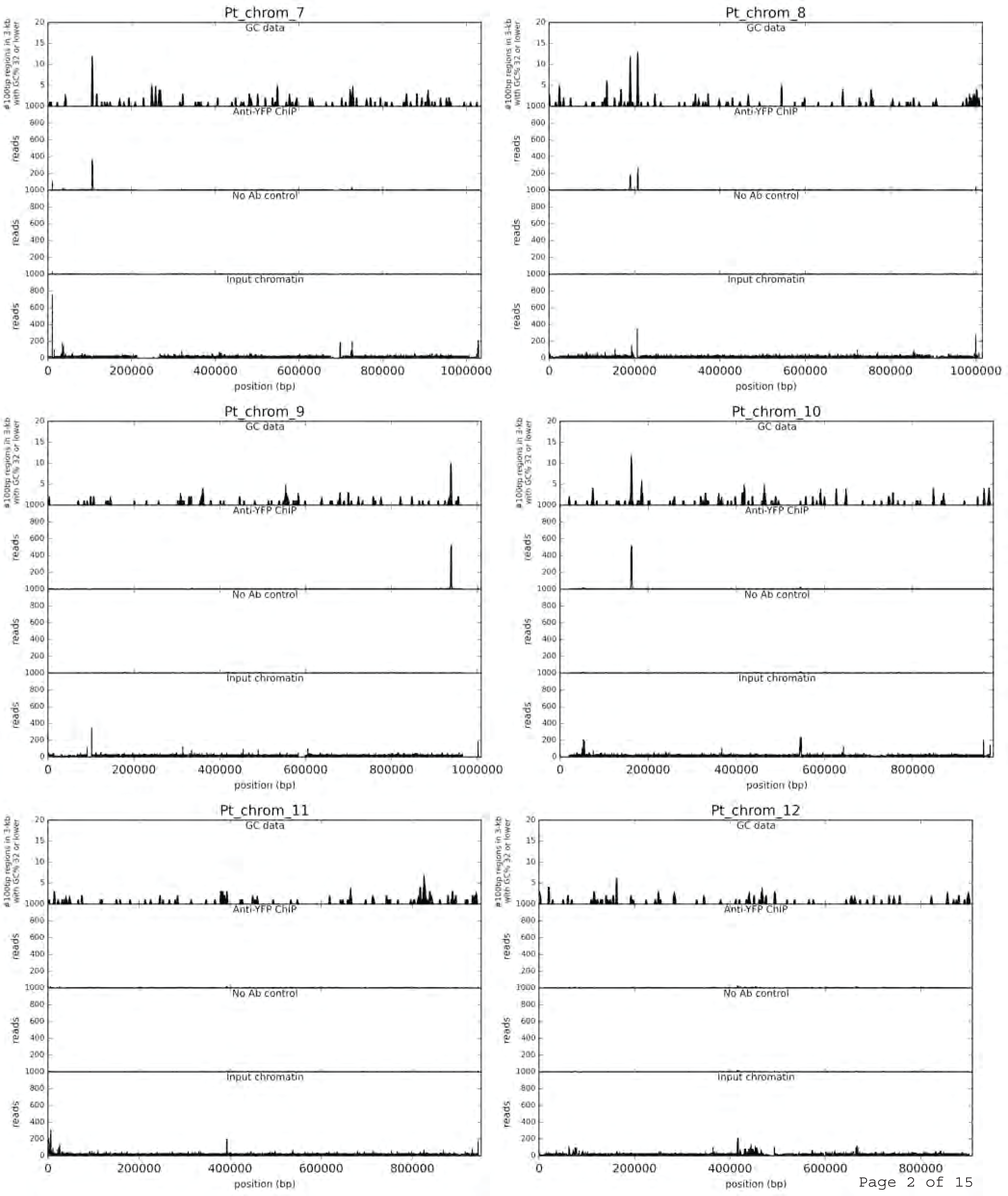
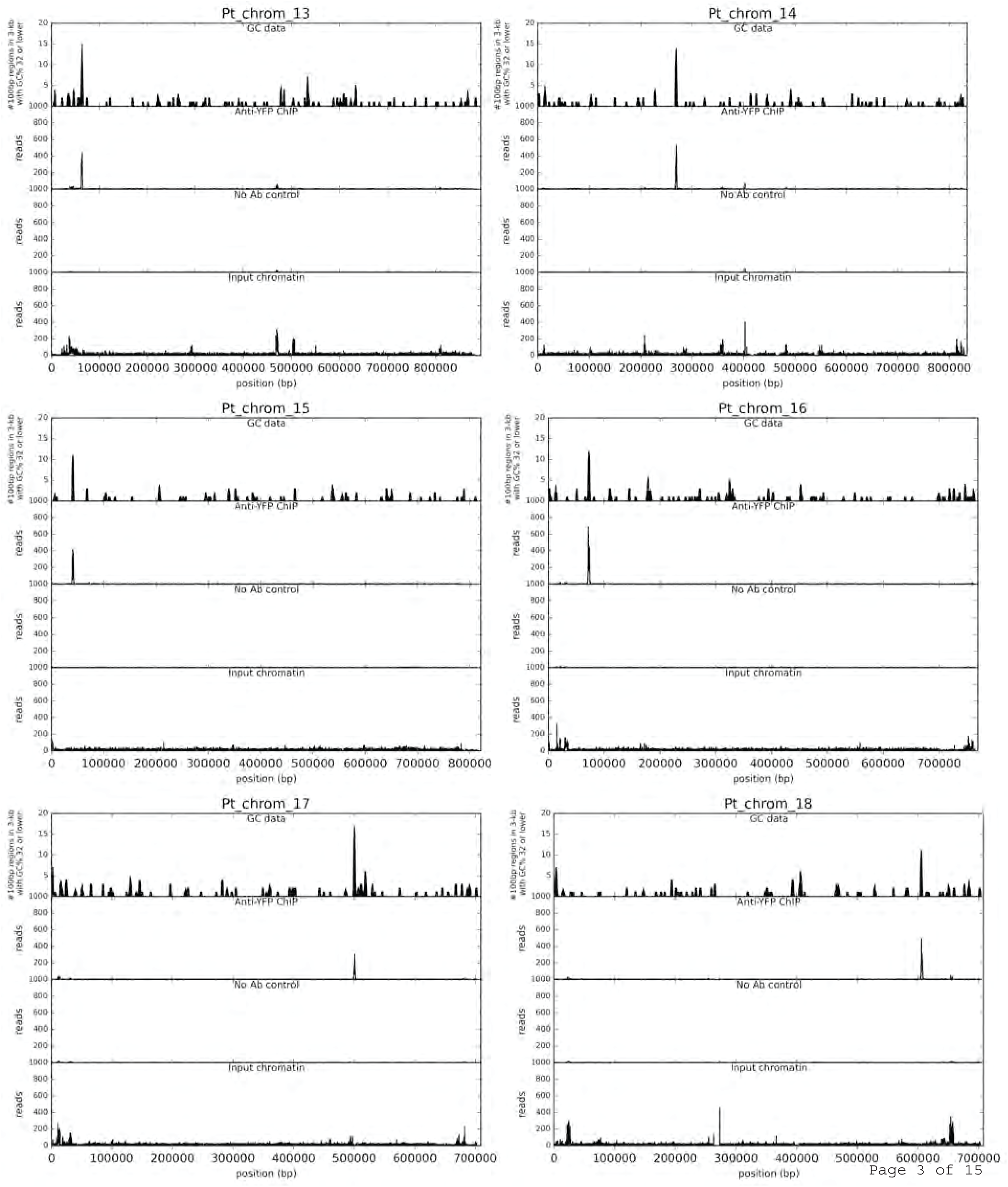
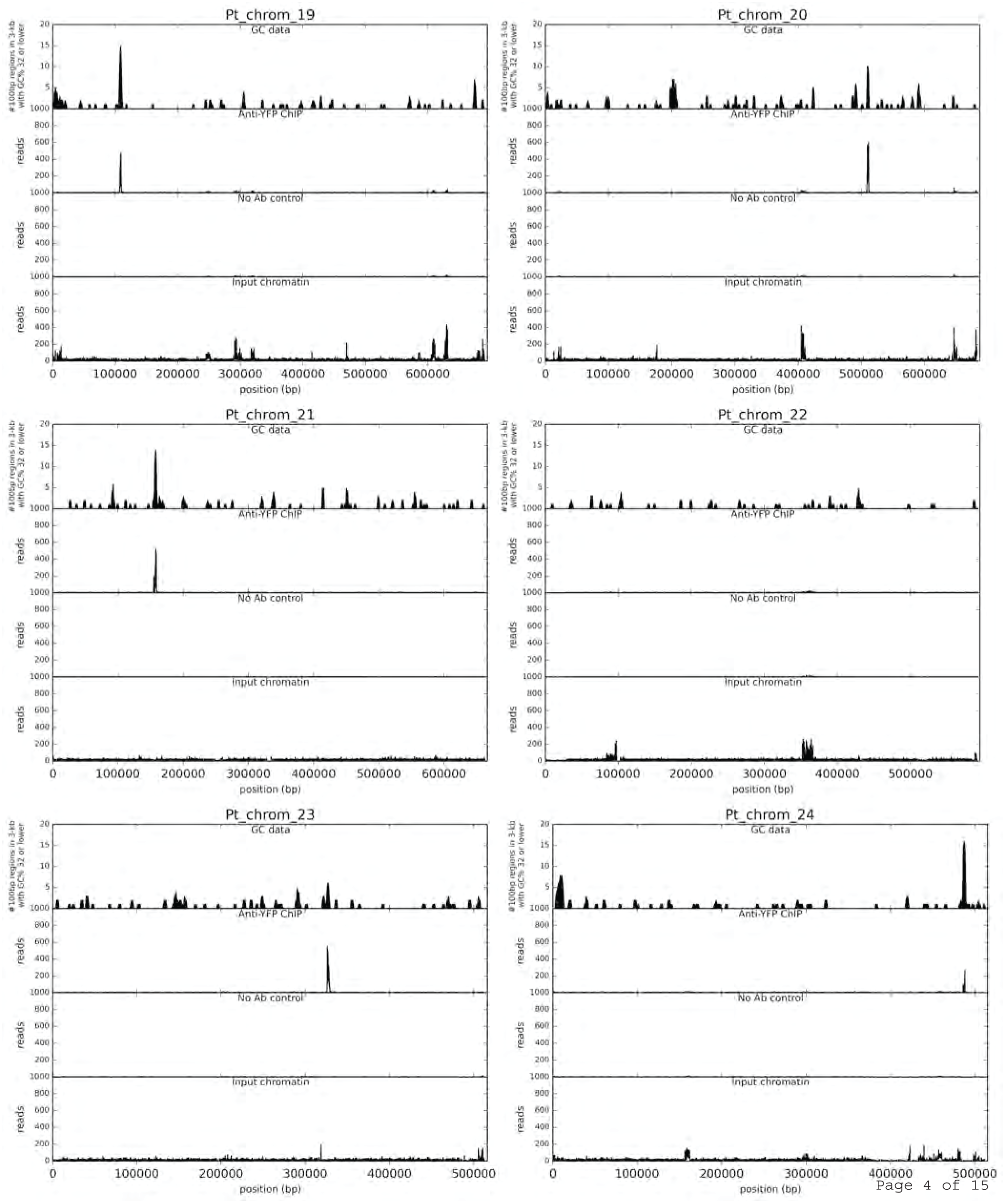
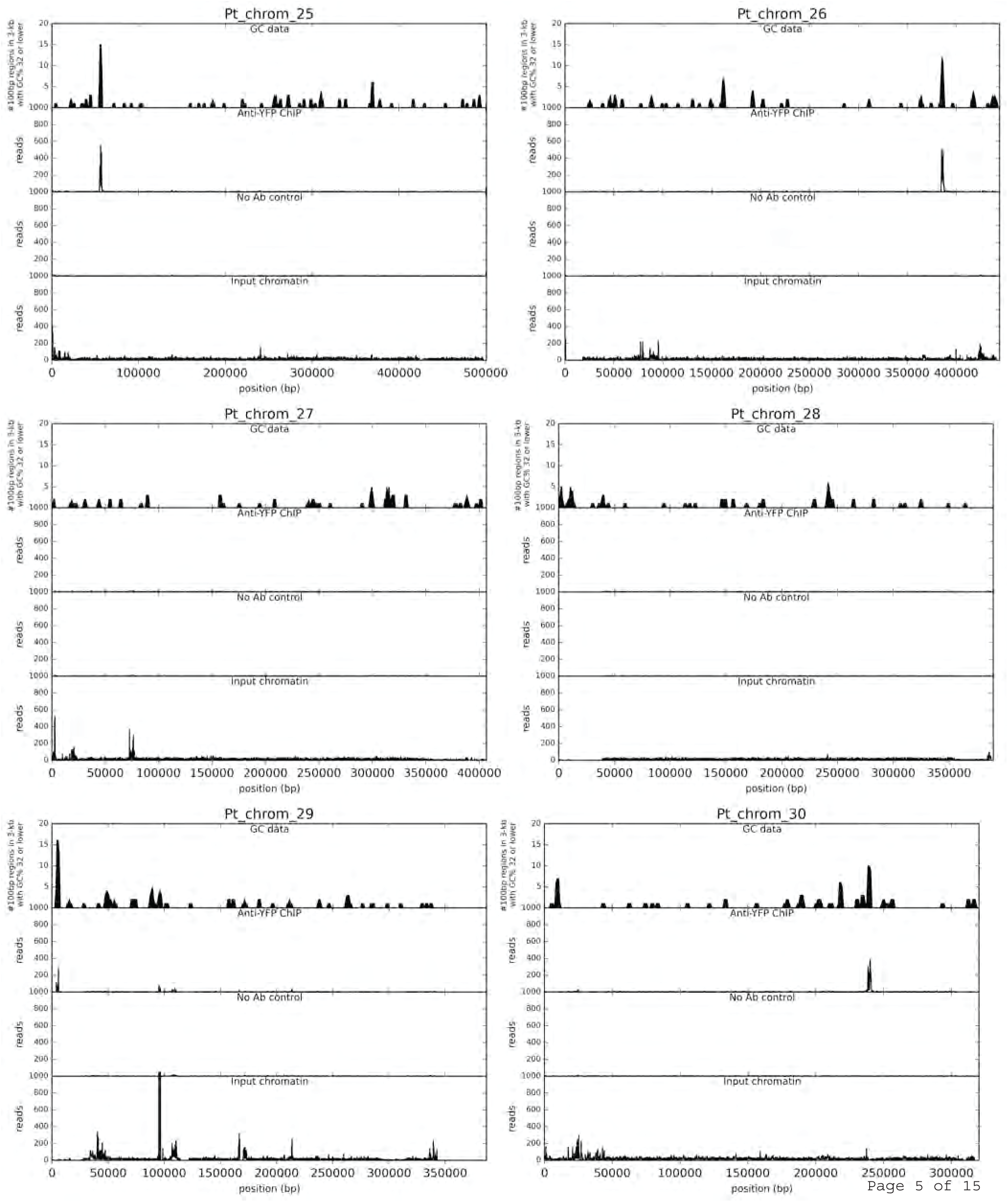
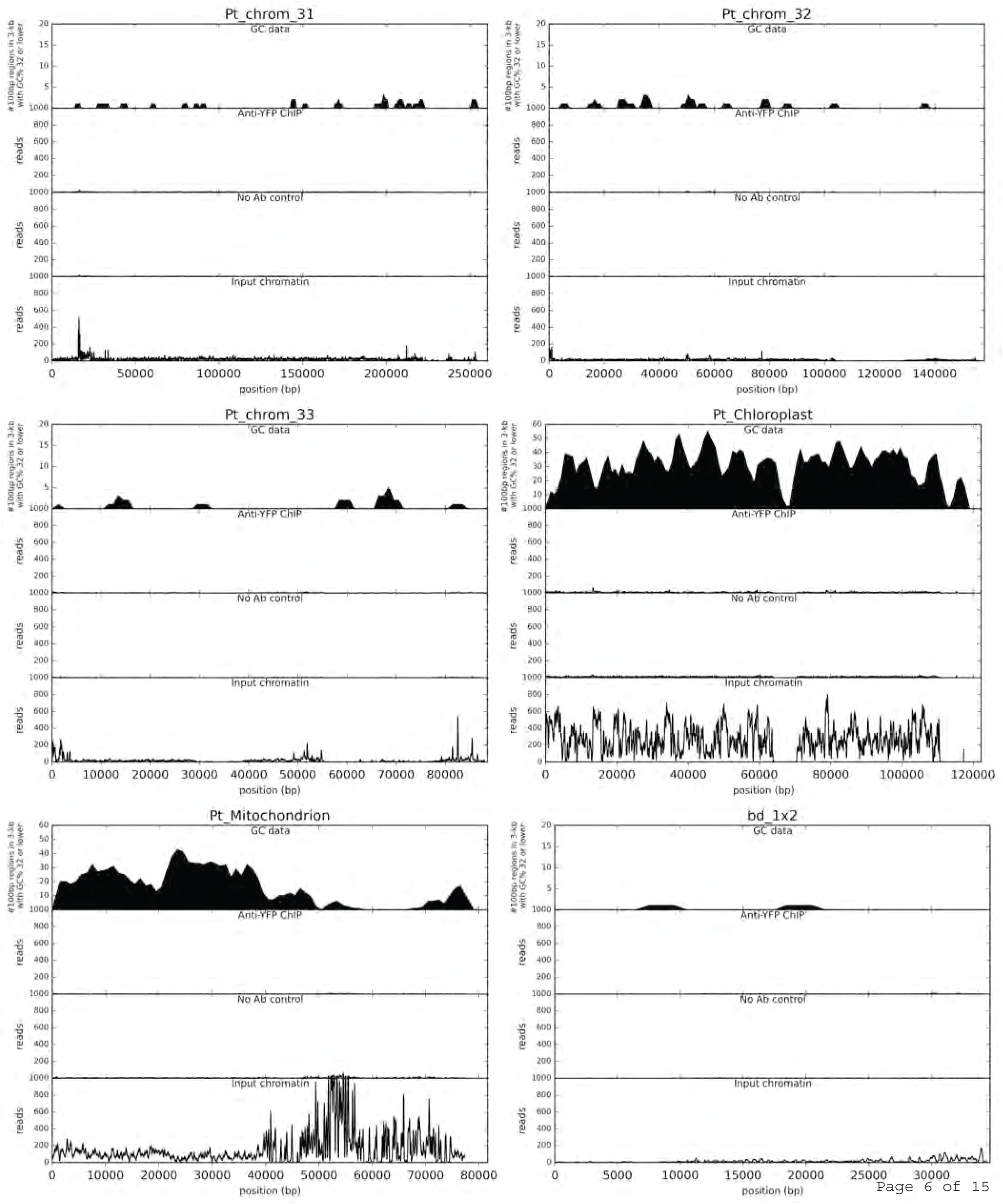
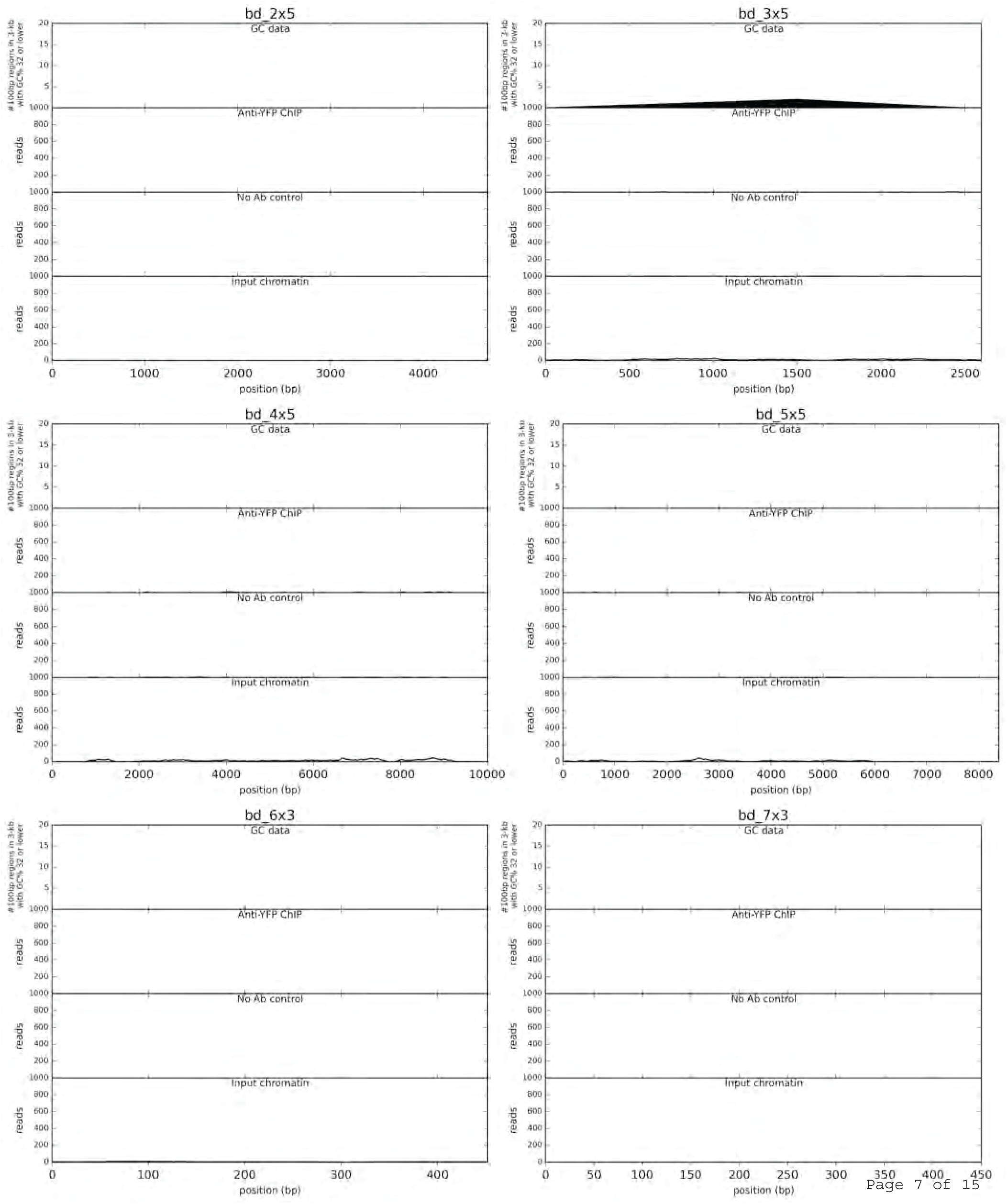
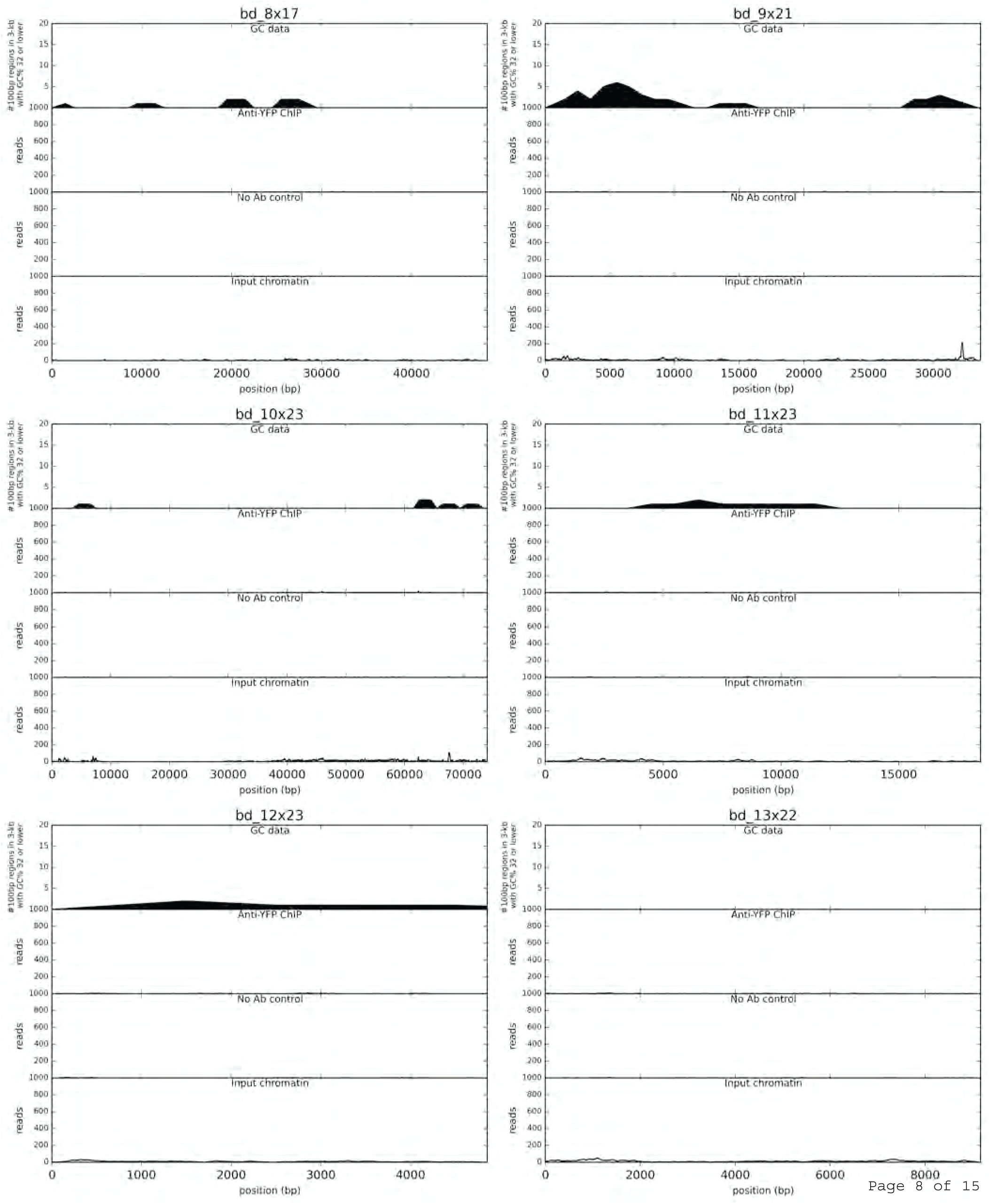
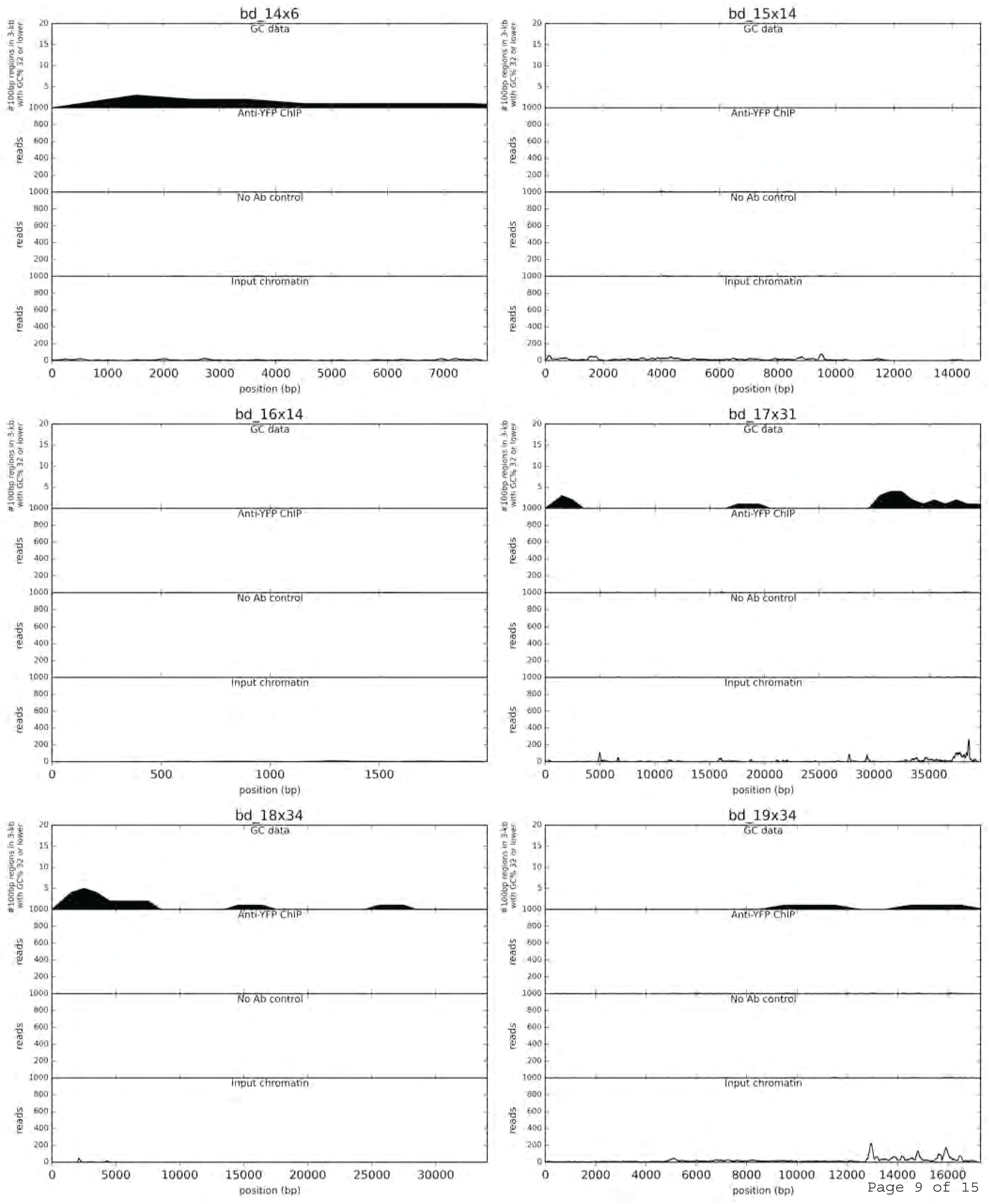
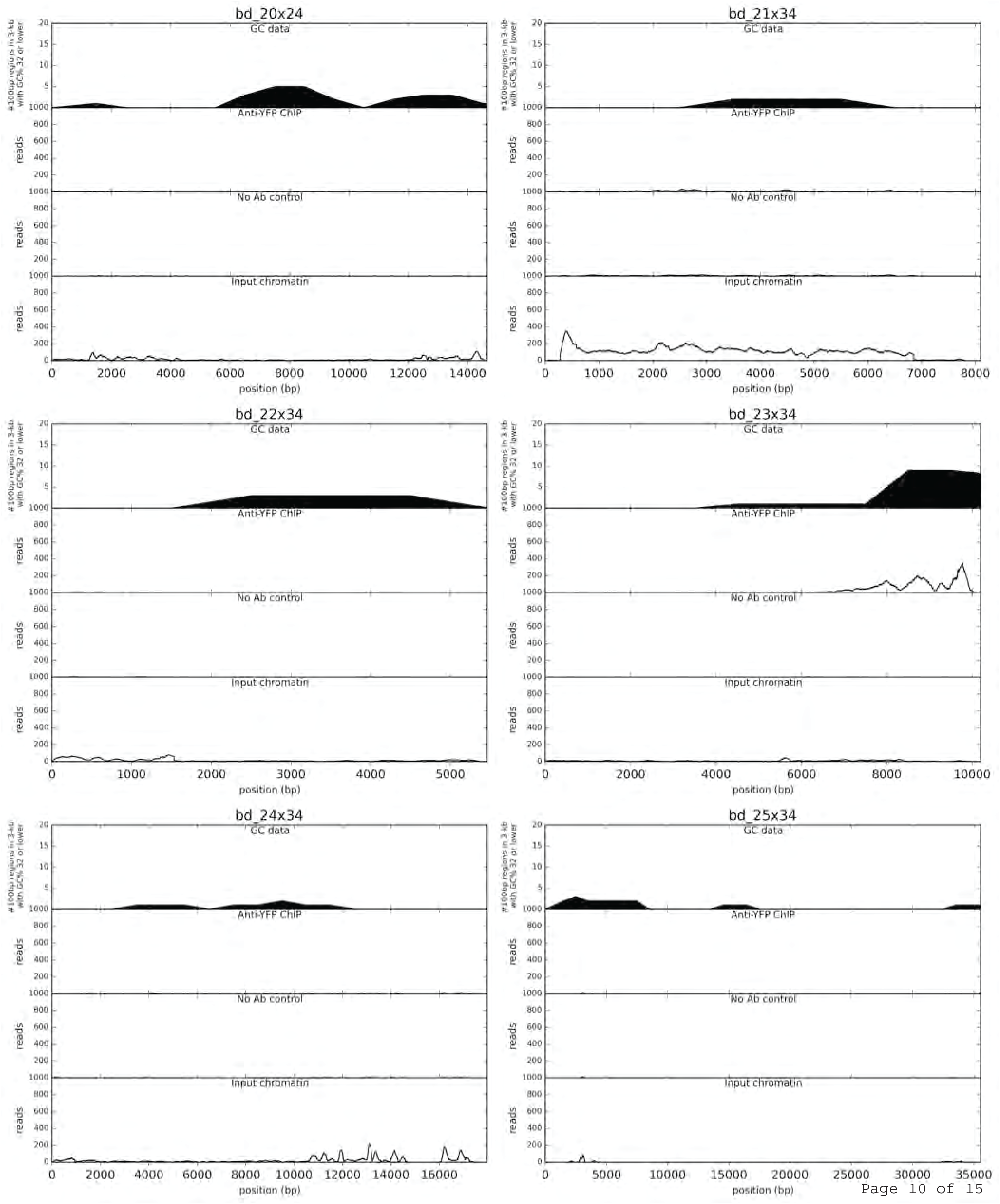
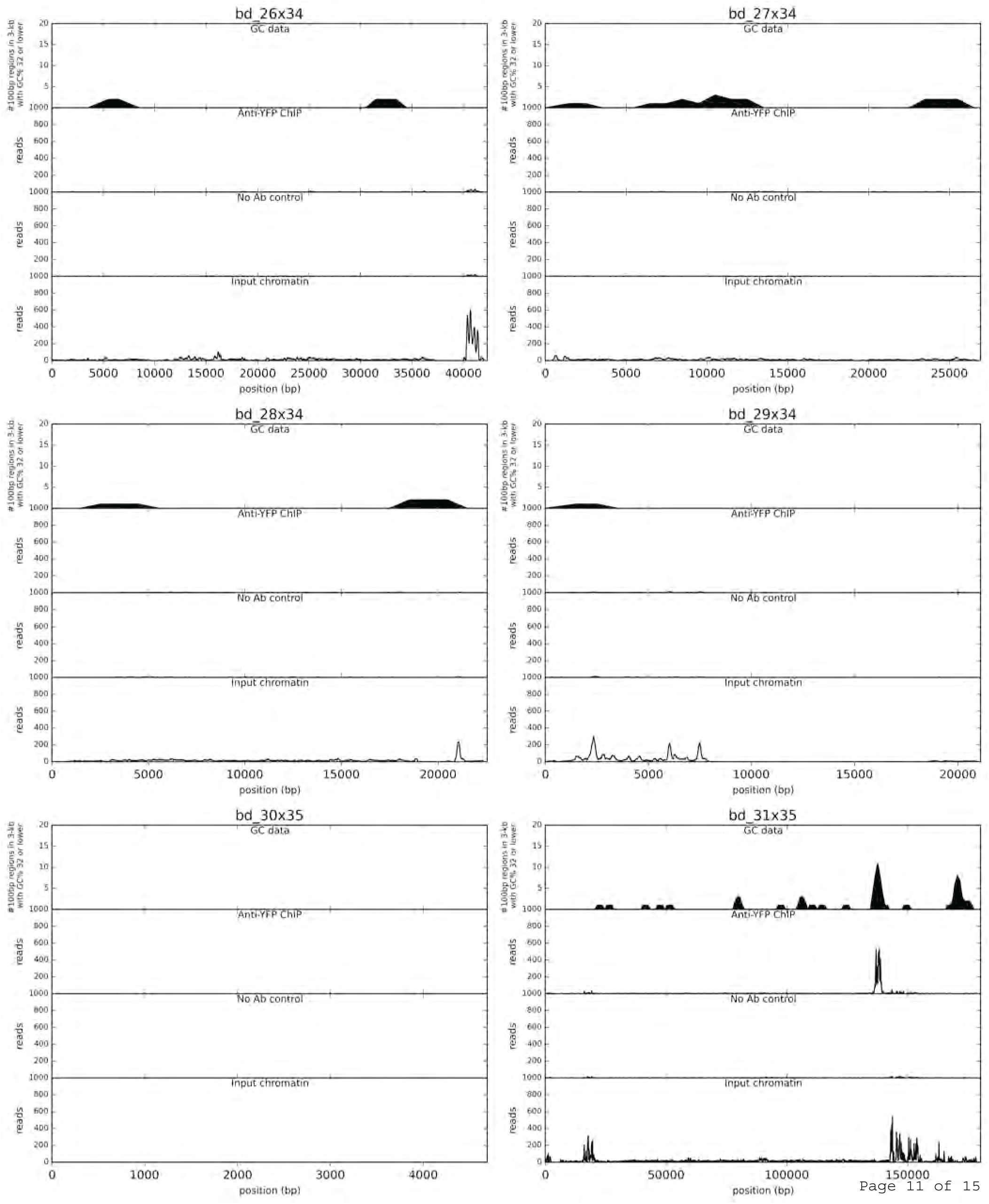
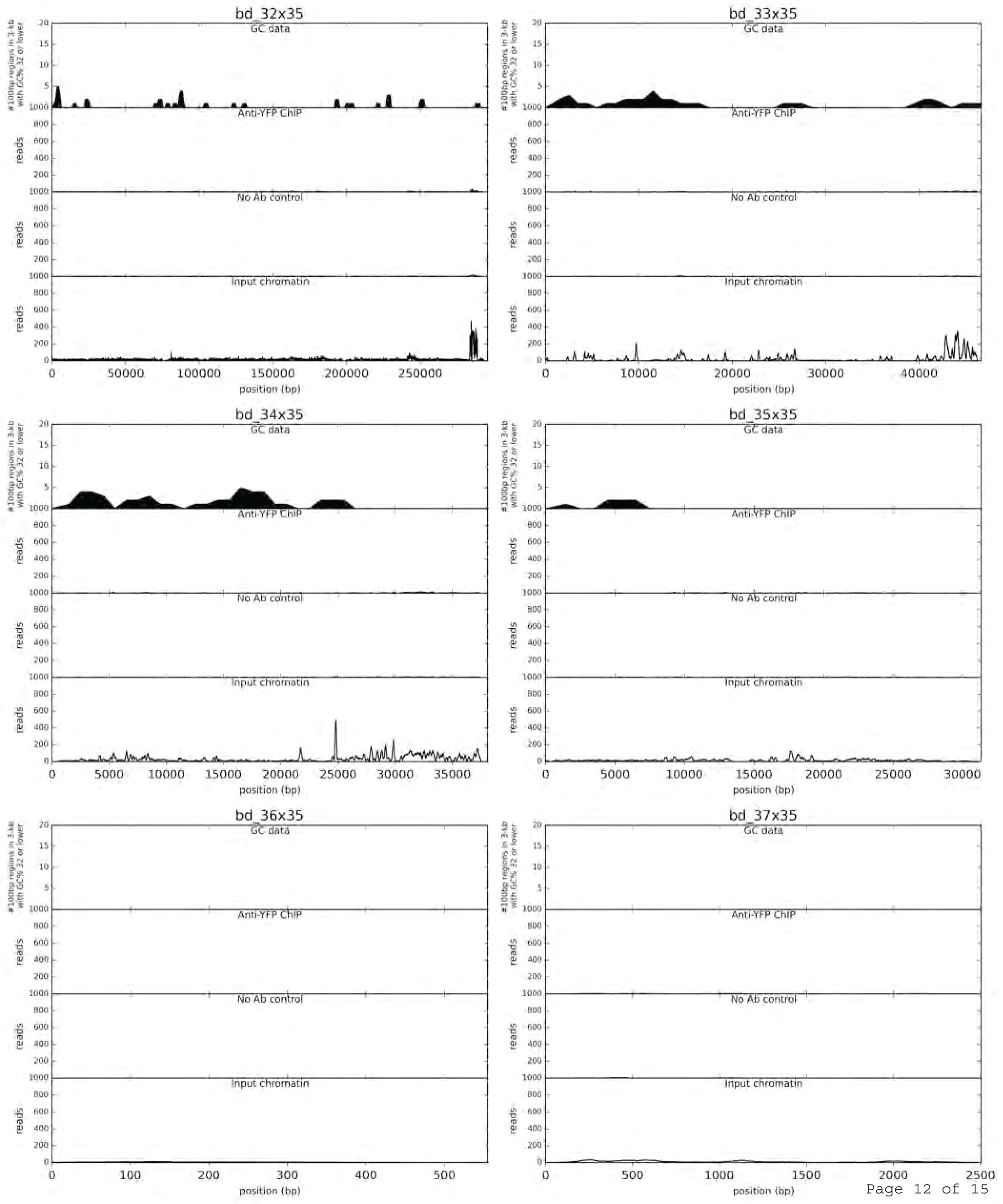
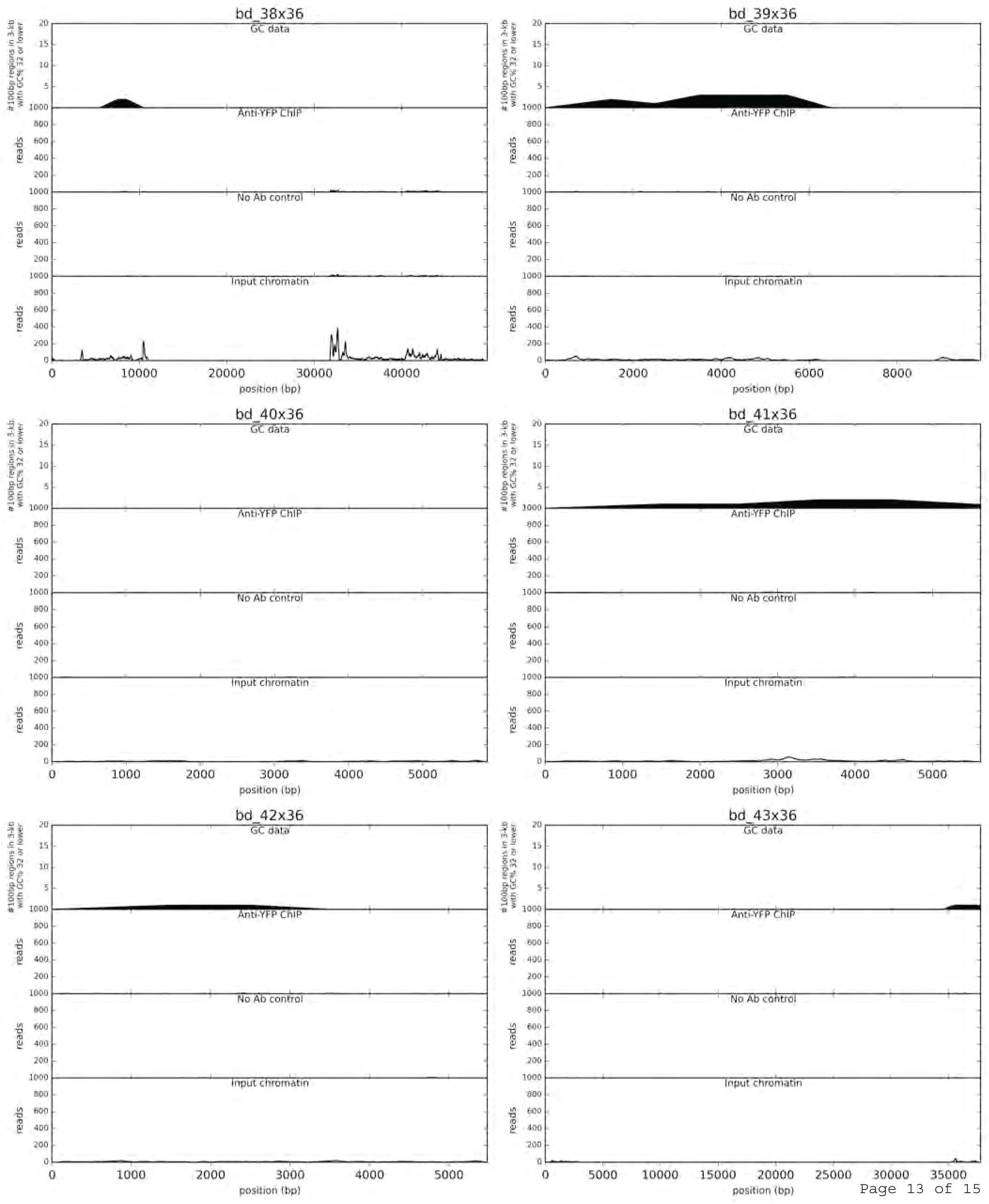
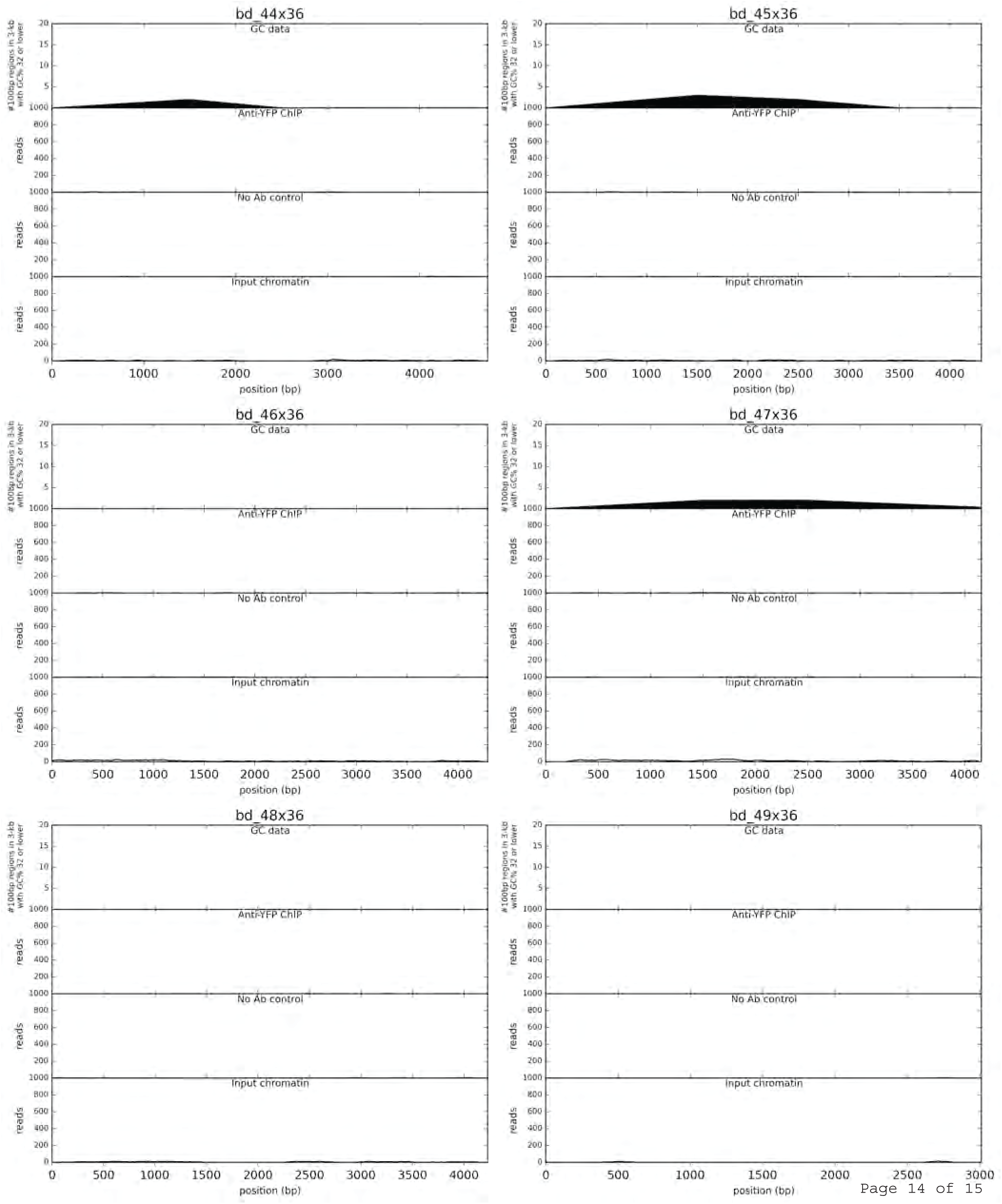
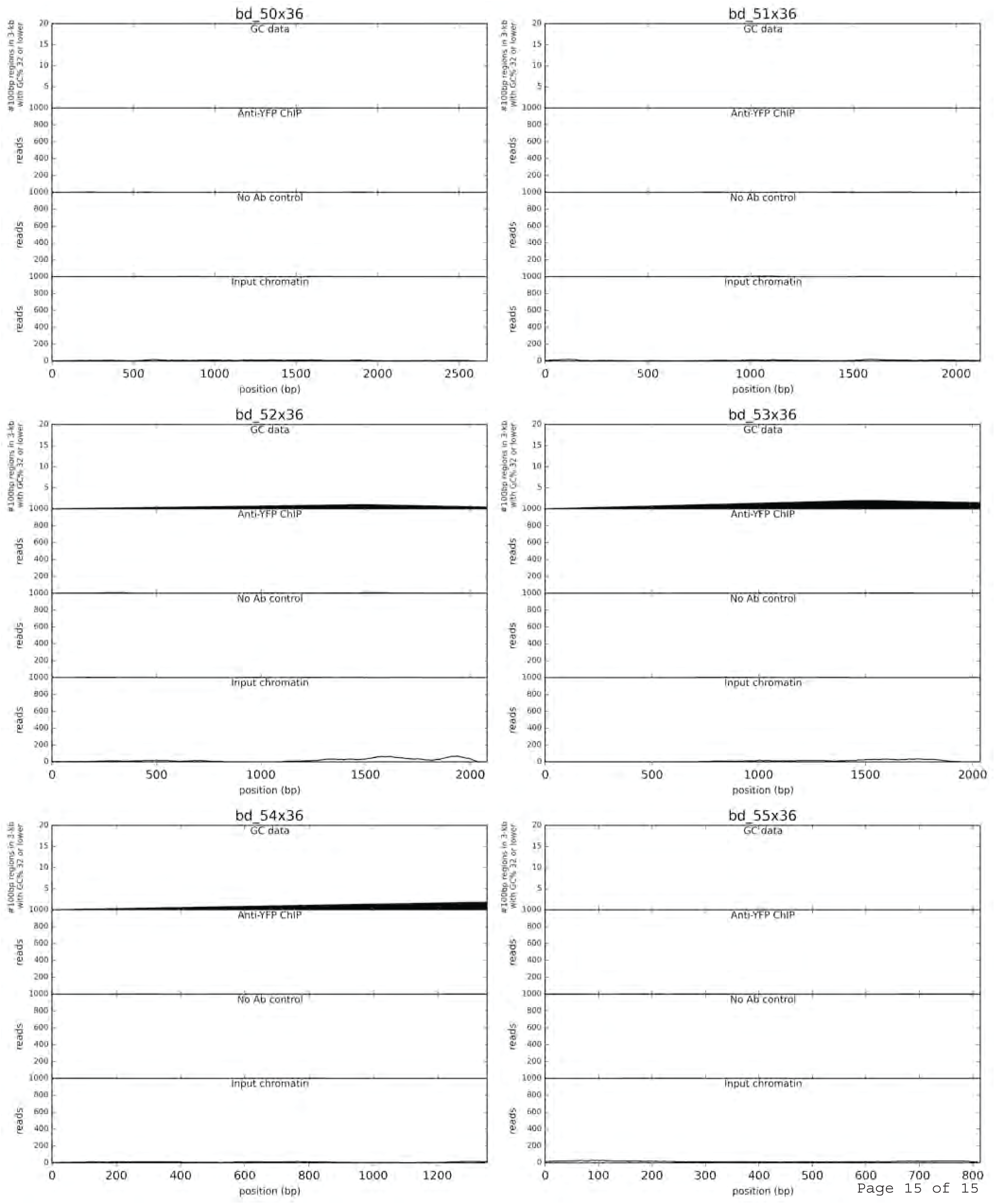
GC data and ChIP-seq data for all chromosome scaffolds, chloroplast and mitochondrion genomes, and bottom drawer non-scaffolded assemblies. The following four data series were plotted for each sequence: 1) Low GC features plotted as the number of 100-bp windows with GC 32% or lower within a larger 3-kb window that advanced by 1-kb each step, 2) ChIP-seq reads for anti-YFP treatment, 3) no antibody ChIP-seq reads, and 4) input chromatin reads for ChIP-seq experiment.

**Supplementary Figure 4:**
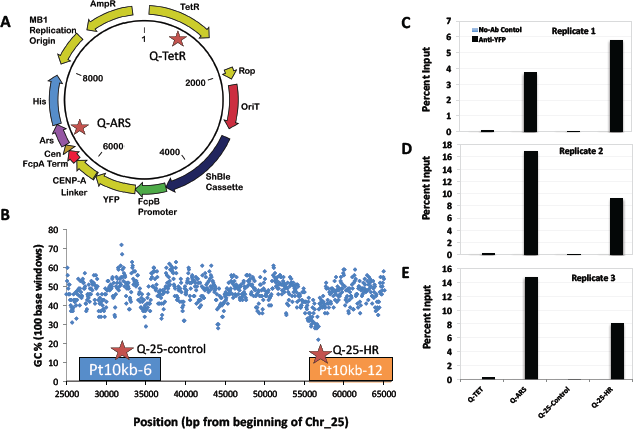
ChIP-qPCR. **A.** Map of the pPtPBR1-YFP-CENP-A with positions of qPCR primer sets indicated by stars. **B.** Map of chromosome 25 with qPCR primer sets indicated by stars. **C-D.** ChIP-qPCR results calculated as percent of input chromatin for each of the two episomal loci and two chromosomal loci. Parts C-D each show a replicate that was performed from a different *P. tricornutum* line expressing YFP-CENP-A.

**Supplementary Figure 5:**
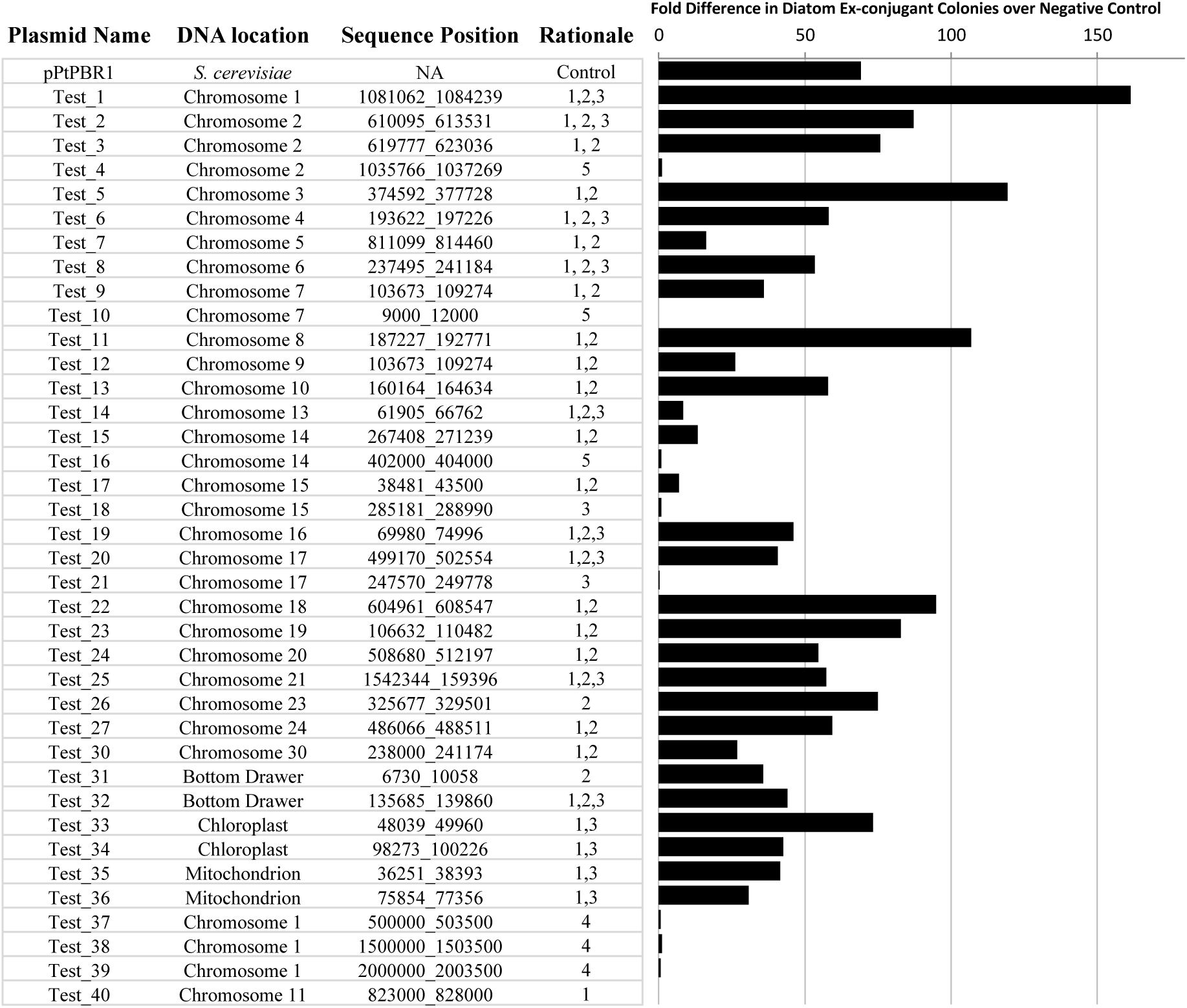
Tests of putative centromeres and other sequences identified by ChIP-seq, bioinformatics analysis, or episome library. Sequences were identified with the following rationales to test for the ability to support episomal maintenance in *P. tricornutum.* 1) Positive for bioinformatic analysis 2) Positive for YFP-CENP-A-ChIP-seq, 3) identified in episome library, 4) designed negative control, and 5) potential YFP-CENP-A ChIP-seq mapping artifact. Plotted are the number of diatom ex-conjugant colonies obtained after conjugation shown as fold increase in colony numbers over the pPtPBR2 empty vector negative control. Each value is the mean of two independent biological replicates with the exception of Test11.

**Supplementary Figure 6:**
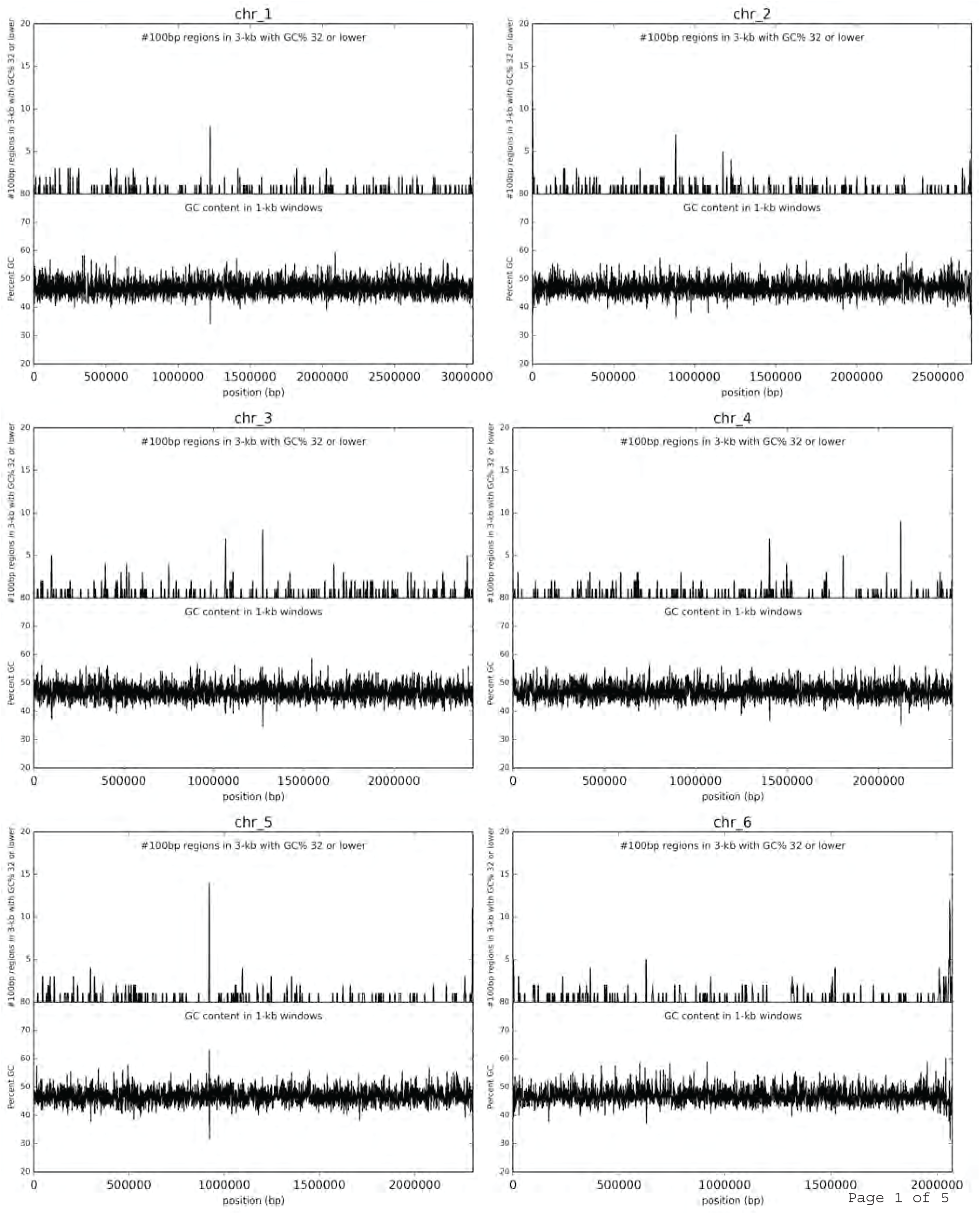
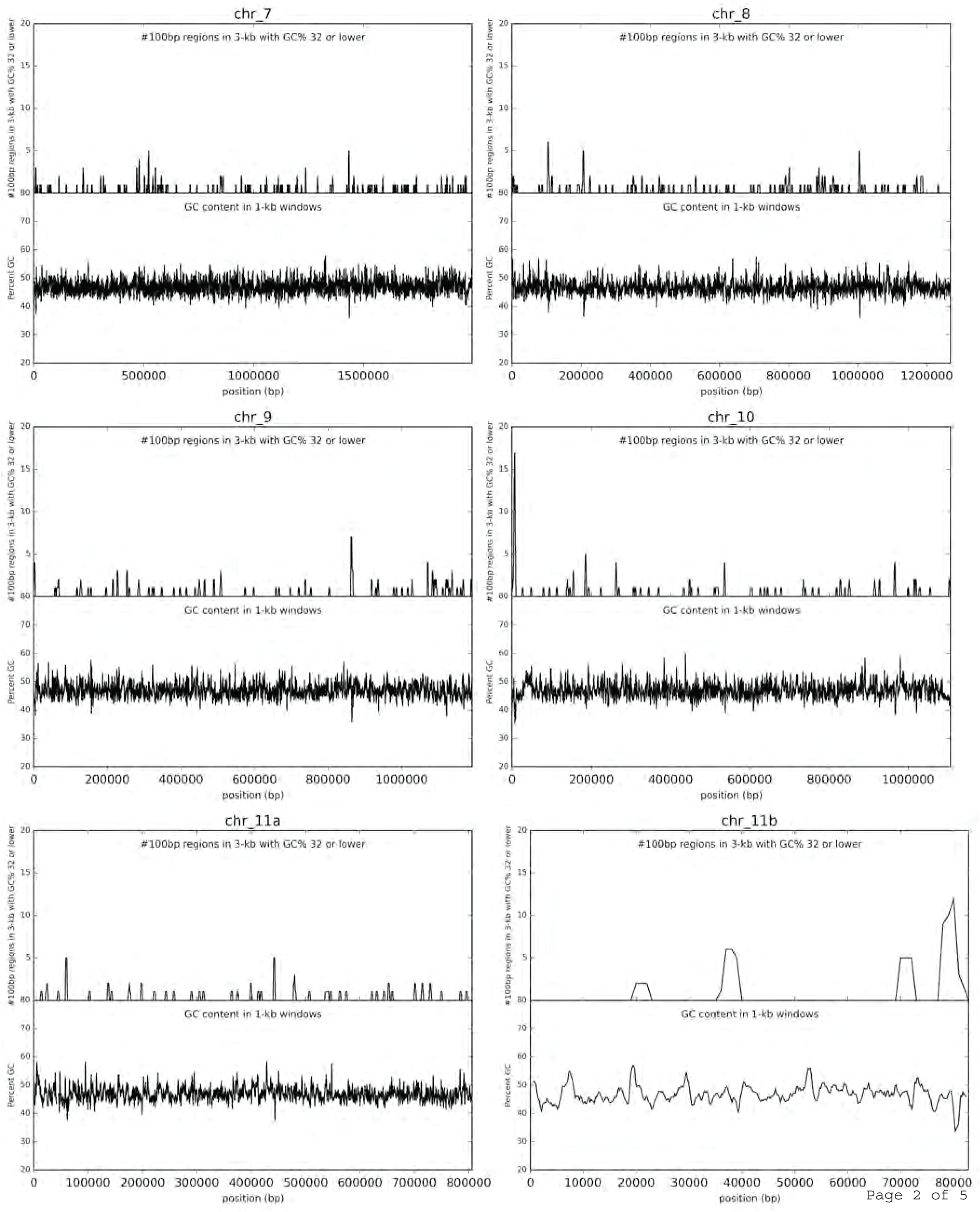
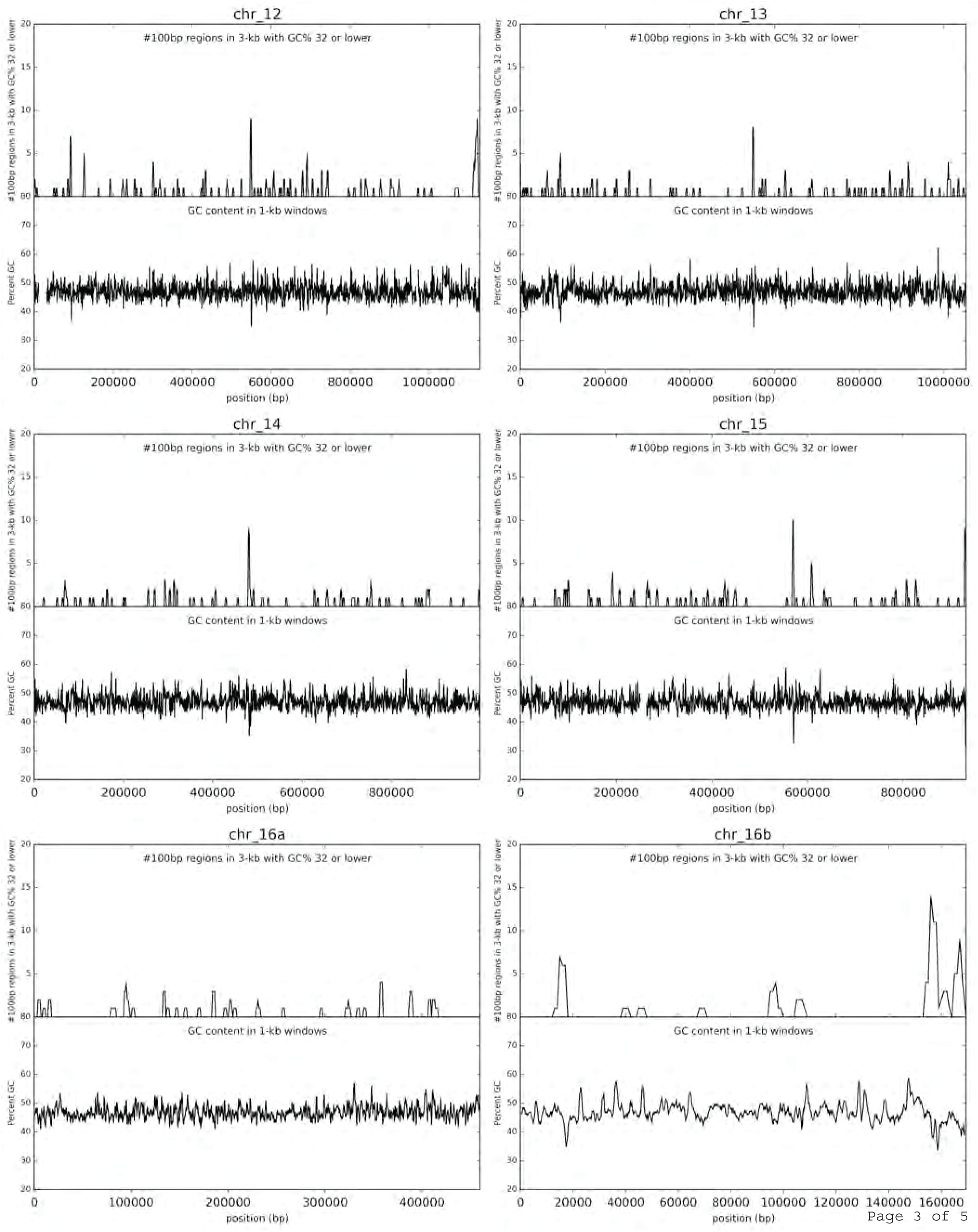
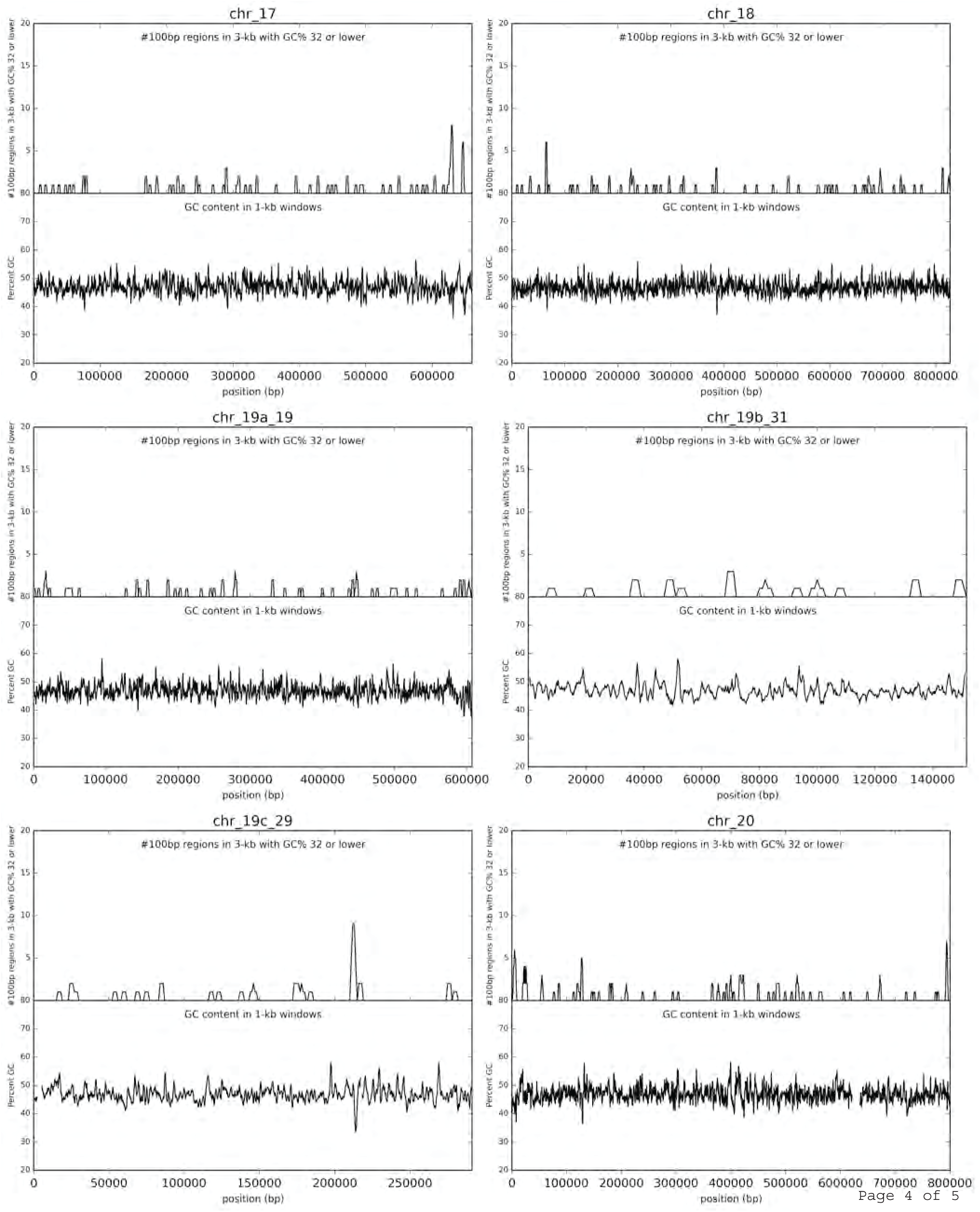
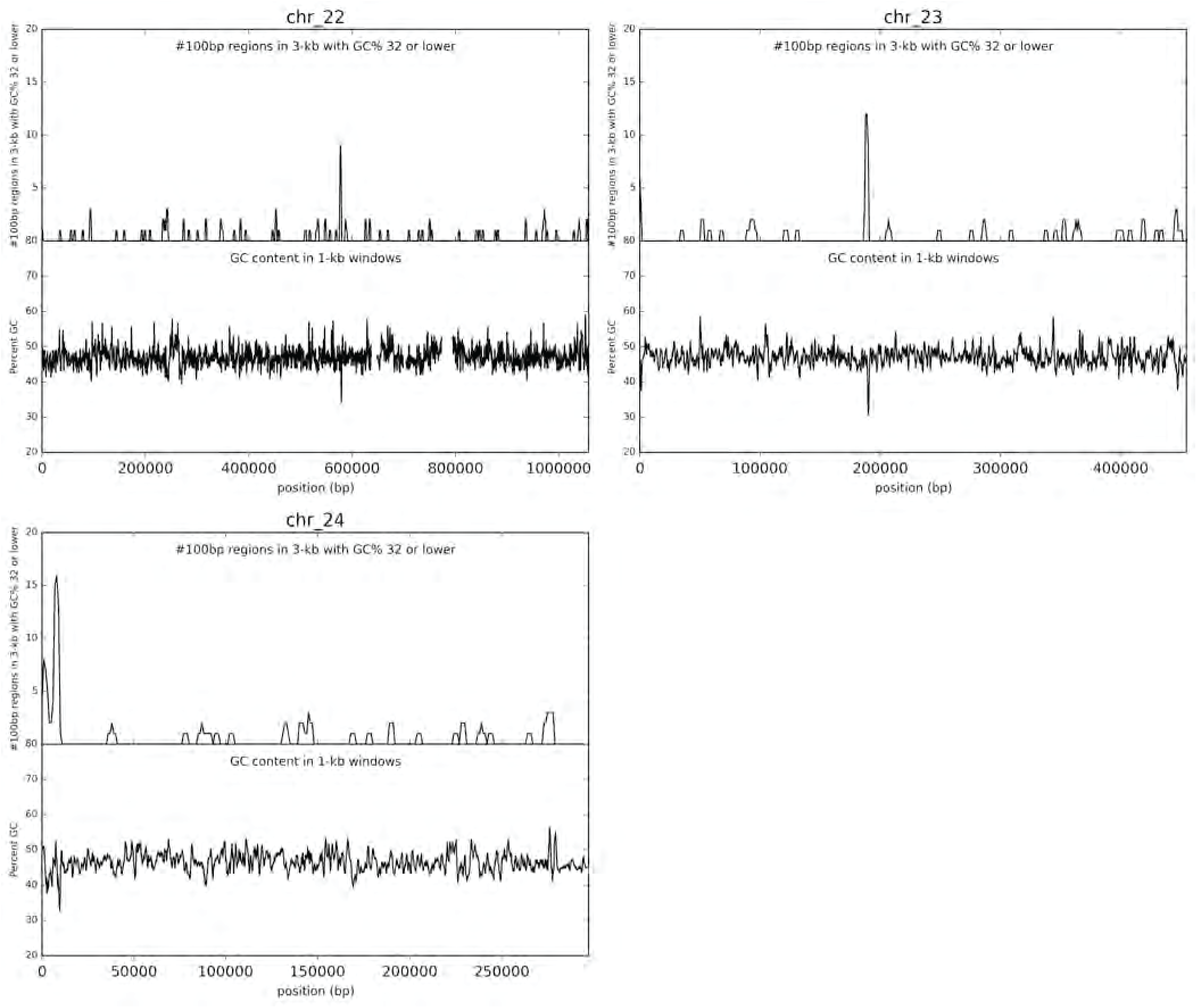
GC analysis of the *T. pseudonana* genome. For each chromosomal scaffold, two data series are plotted: 1) the number of 100-bp windows with GC less than or equal to 32% in a larger 3-kb window that advanced by 1-kb each step, and 2) the GC in a sliding 1-kb window that advanced by 200-bp each step.

**Supplementary Figure 7:**
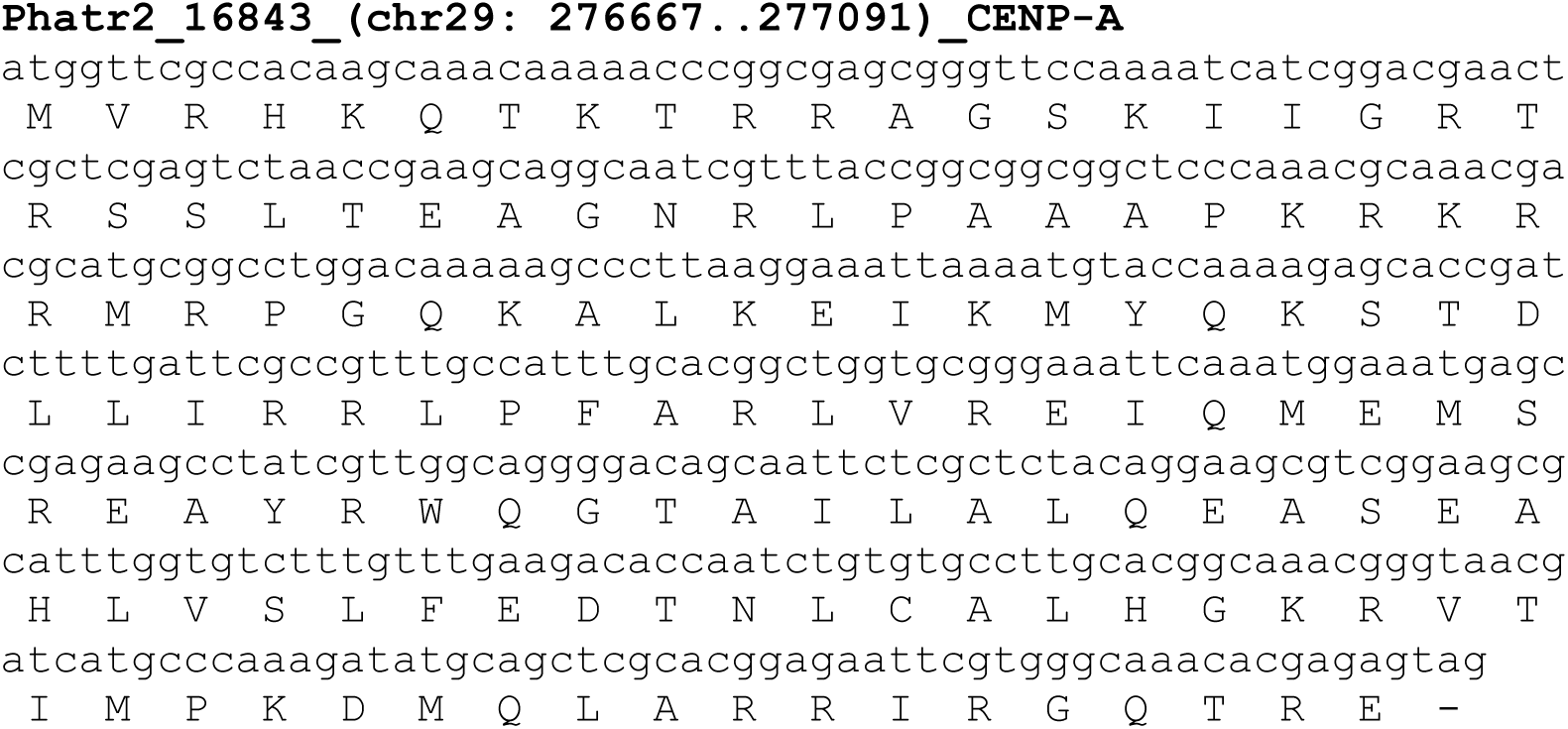
DNA sequence, translation, and genome coordinates of *P. tricornutum* CENP-A used for cloning and expression of YFP-CENP-A fusion protein used for ChIP analyses.

**Supplementary Figure 8:**
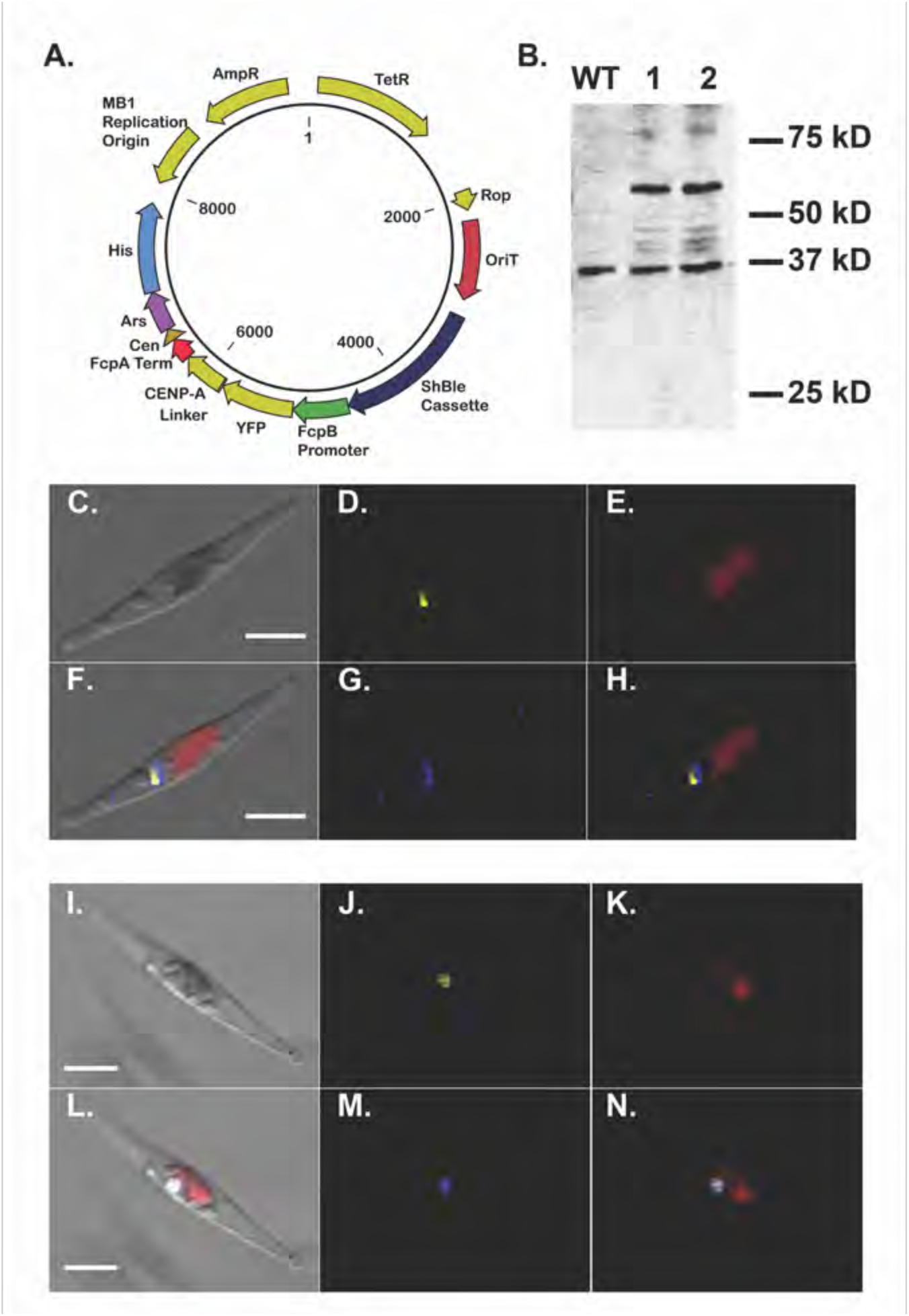
The YFP-CENP-A fusion is expressed in *P. tricornutum* and localized to the nucleus. **A**. Vector map showing plasmid pPtPBR1-YFP-CENP-A expressed under FcpB promoter on the episome. **B.** Western blot showing wild type (WT) and two ex-conjugant lines expressing the YFP-CENP-A fusion (1 and 2). The expected size of the YFP, Gly5-Ala-spacer and CENP-A fusion is expected to be 43.3 kDa. Multiple bands in the 40-45 kDa region are visible in the YFP-CENP-A lines and absent in the wild type and may correspond to posttranslational modifications of the YFP-CENP-A fusion. A prominent band at ˜60 kDa is also differentially observed in the YFP-CENP-A lines. It is unknown if this also corresponds to a modified YFP-CENP-A protein. **C-N**: Scanning electron micrographs showing Brightfield **(C and I)**, YFP **(D and J)**, chlorophyll autofluorescence **(E and K)**, DAPI **(G and M)**, merged fluorescent channels **(H and N)** and all channels merged **(F and L)**. The YFP-CENP-A signal localizes specifically to the nucleus. Scale bar indicates 5 μm.

**Suplementary Table 1:**
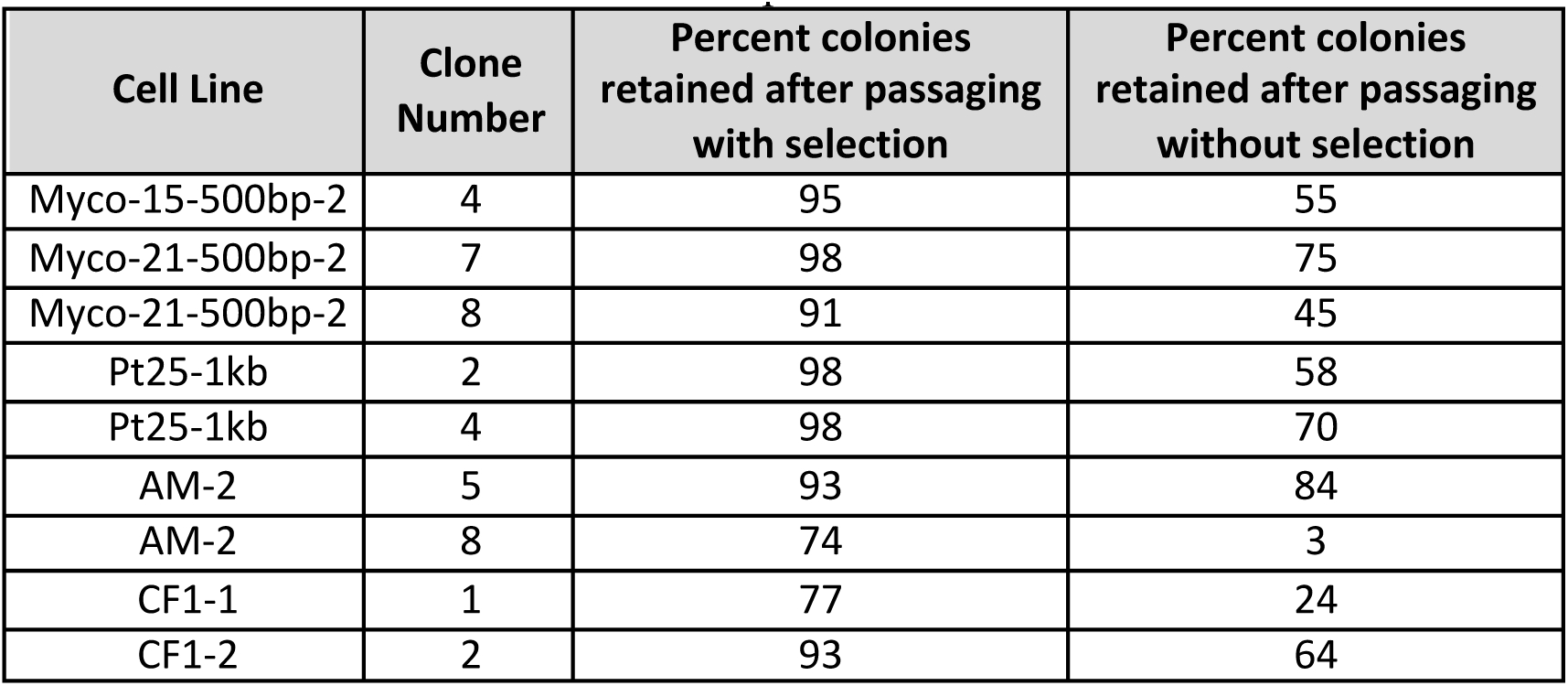
Stable maintenance of episomes in *P. tricornutum* enabled by DNA from foreign and native sources. Each clone examined is presented with the percentage of colonies that retained antibiotic resistance abilities after 30 days of passaging with and without selection. A minimum of 40 colonies were plated for each clone.

**Suplementary Table 2:**
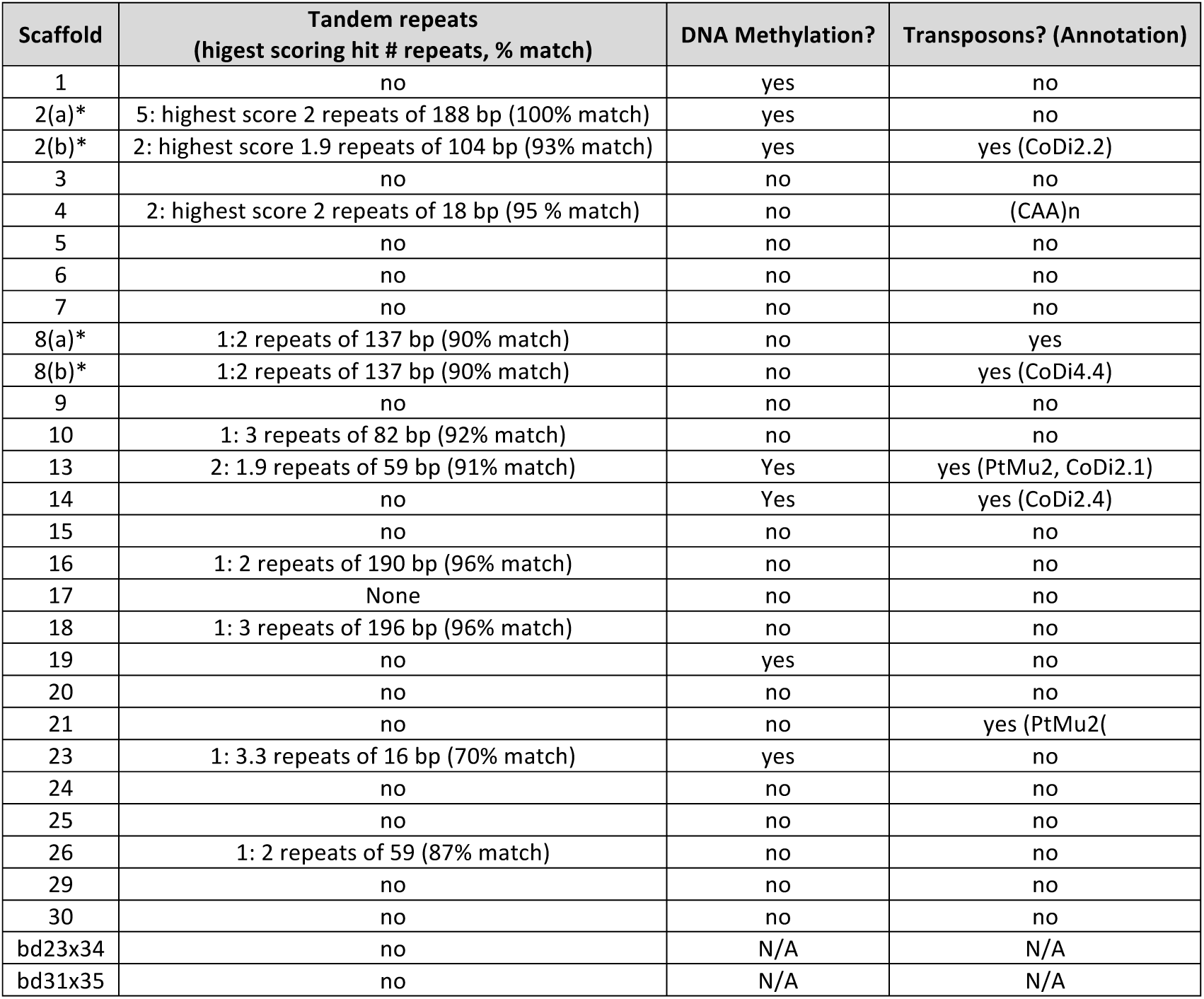
Analysis of genetic features associated with putative centromeres on each chromosome scaffold, including tandem repeats (tandem.bu.edu/trf/trf.html), DNA methylation, and transposon presence (http://ptepi.biologie.ens.fr/cgi-bin/gbrowse/Pt_Epigenome/). Star (*) indicates that two putative centromeres were identified for that chromosome scaffold, and each was analyzed.

**Suplementary Table 3:**
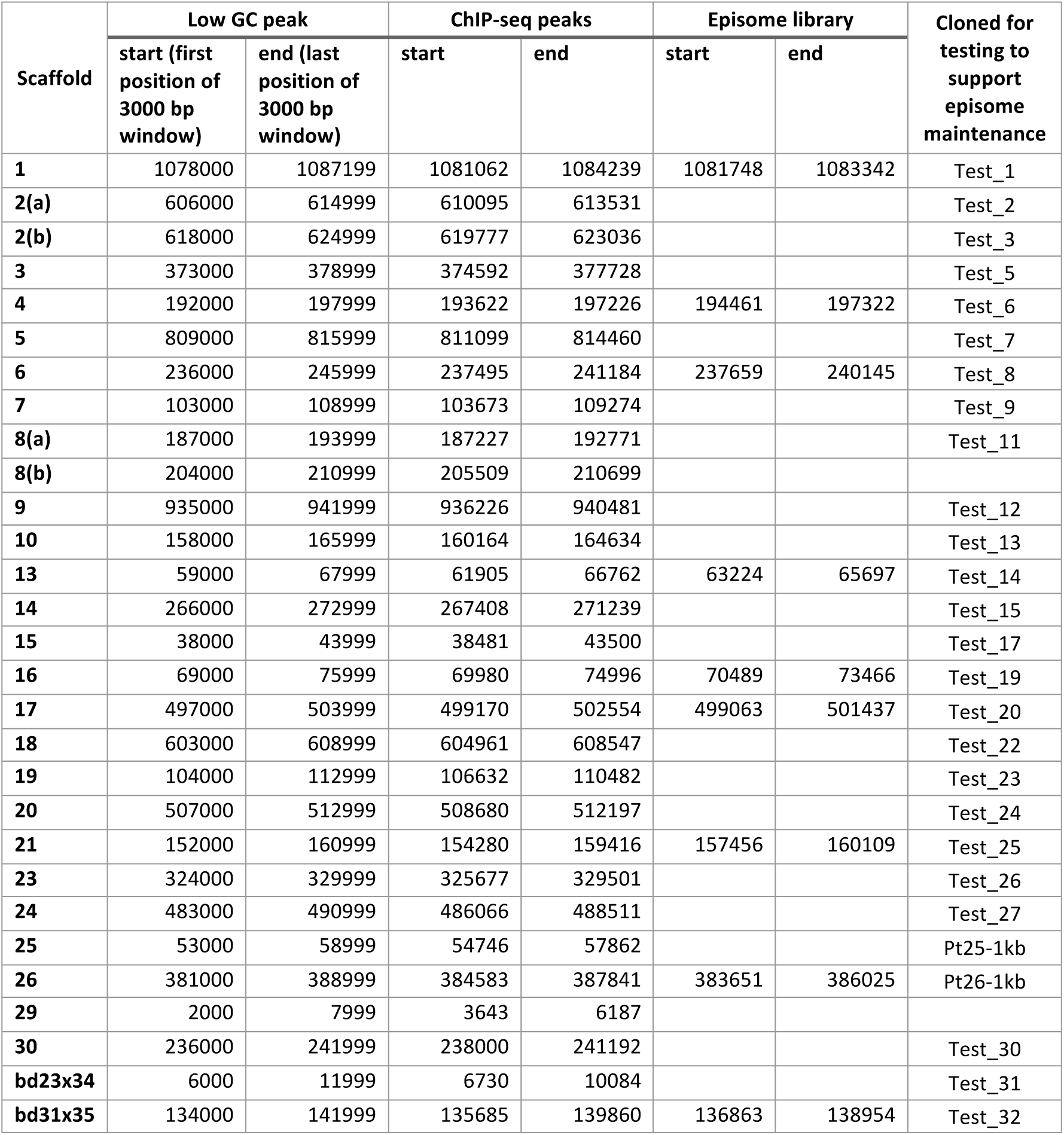
Putative *P. tricornutum* centromeres are identified by co-localization of peaks of low GC DNA, ChIP-seq peaks, and fragments recovered from the episome library. Peaks of low GC content in the scaffold is the region with the largest number of 100-bp windows with GC equal or lower than 32% in a larger 3-kb window that advances 1-kb each step. Presented are the boundaries of the peak above background which spans multiple 3-kb windows. The ChIP-seq peaks are the chromosomal regions containing mapped reads higher than background values for the scaffold and with no substantial peak in the input or no antibody controls. The episome library coordinates describe the boundaries of the fragment cloned in the library that conferred episomal maintenance.

**Suplementary Table 4:**
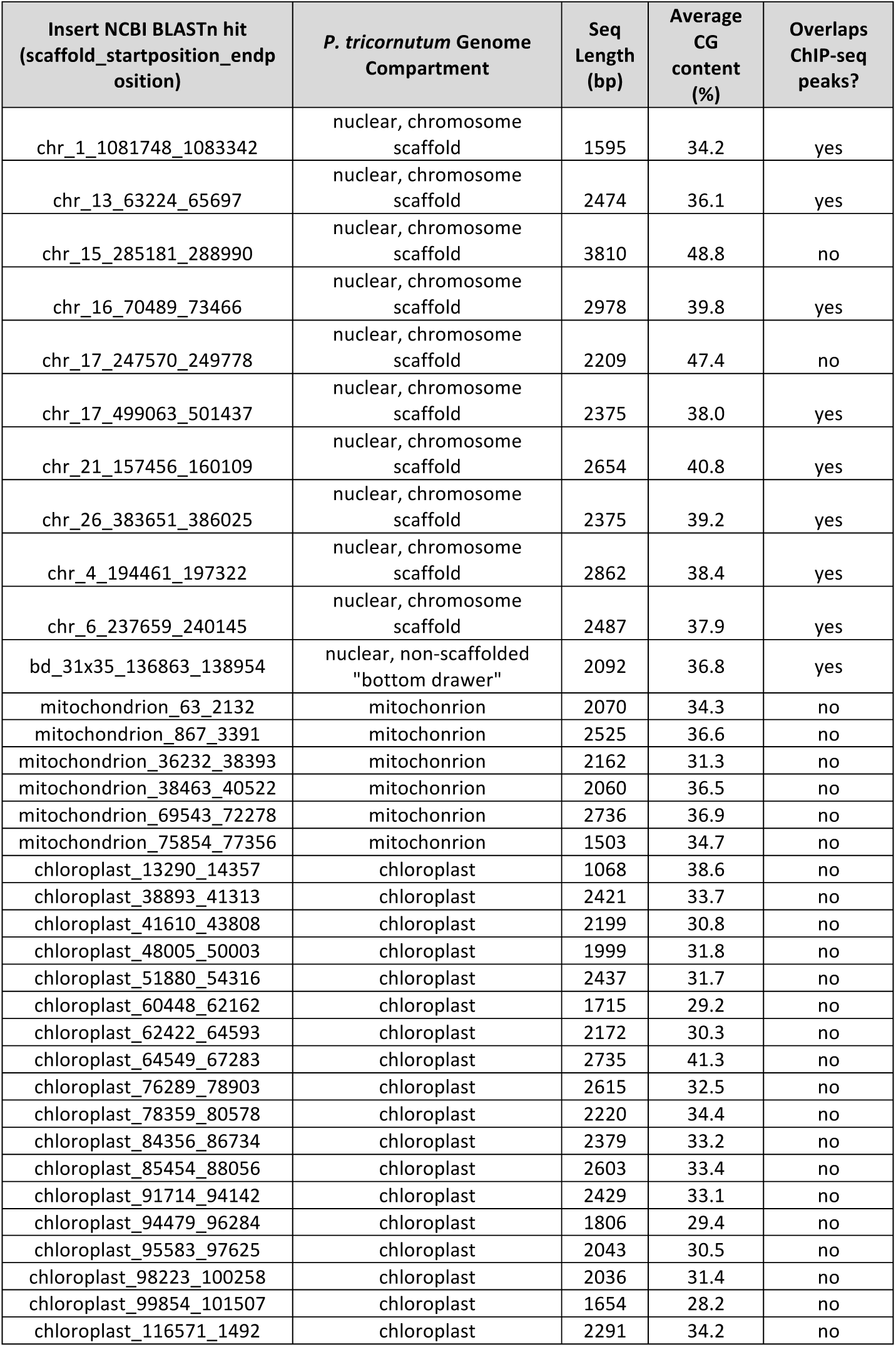
*P. tricornutum* sequences recovered from the forward genetic screen after post-conjugation re-isolation. Sanger DNA sequencing reads for the insert portion of the recovered episome were searched by BLASTn against the set of sequences including the *P. tricornutum* chromosome scaffolds, bottom drawer sequence, and chloroplast and mitochondrion genomes and predicted boundaries of the insert are included for each recovered sequence, as well as the predicted sequence length, average GC content, and whether there were associated ChIP-seq peaks identified in this study.

**Suplementary Table 5:**
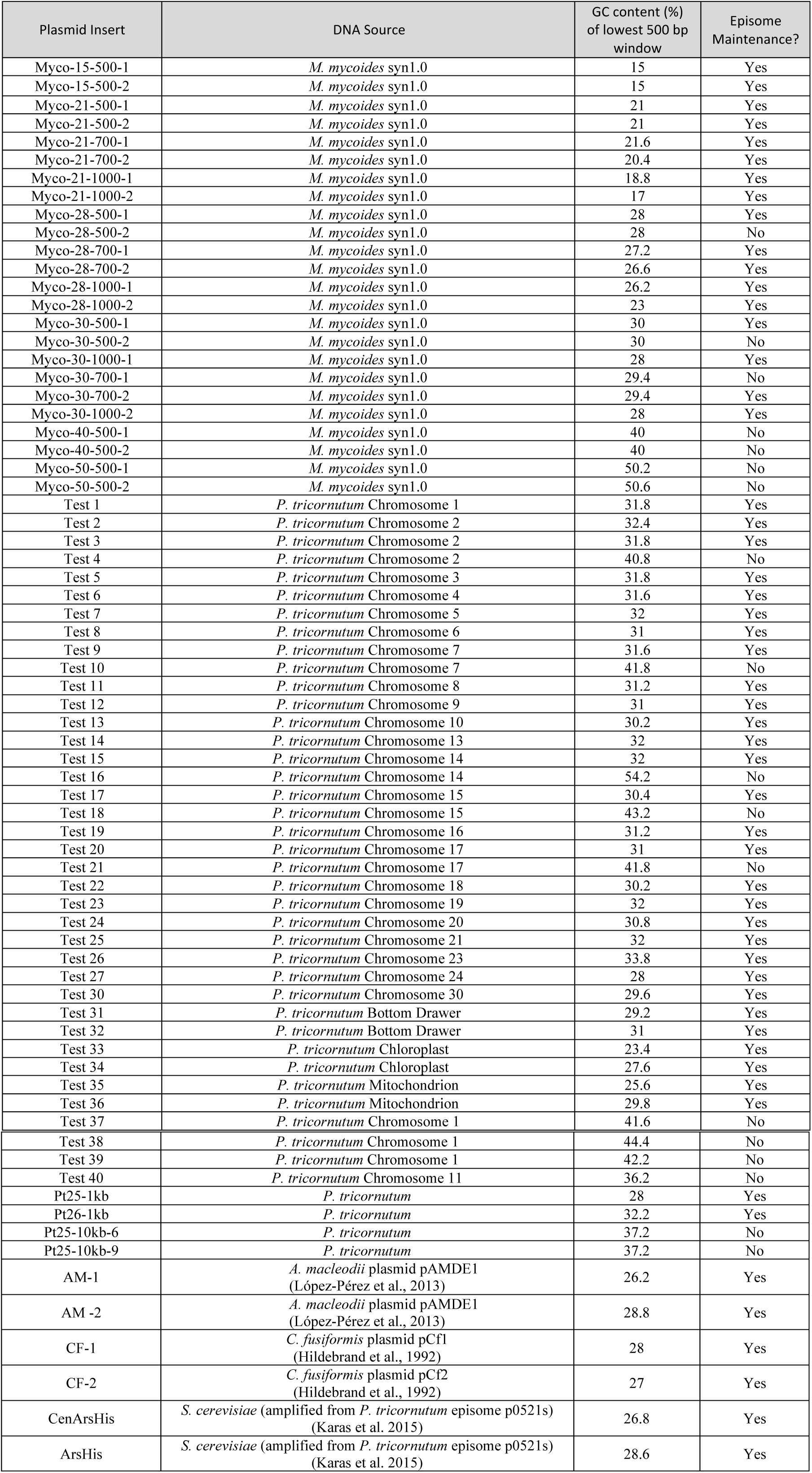
DNA inserts from various sources examined for ability to maintain an episome in *P. tricornutum*, and the GC content of the lowest 500-bp window in each sequence.

**Suplementary Table 6:**
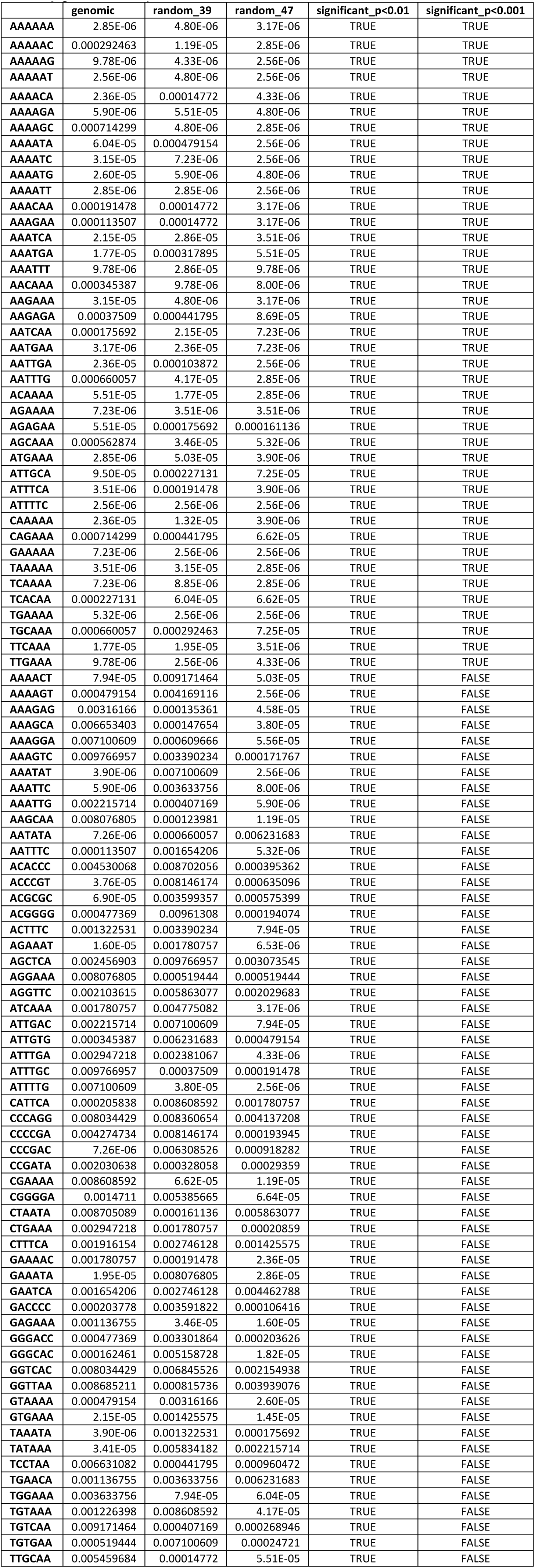
Identification of 6-mer sequences significantly enriched in *P. tricornutum* centromeres. P-values of statistical tests for each 6-mer for centromeres compared to randomly selected genomic sequences, and randomly generated sequences at 39% GC and 47% are shown.

**Suplementary Table 7:**
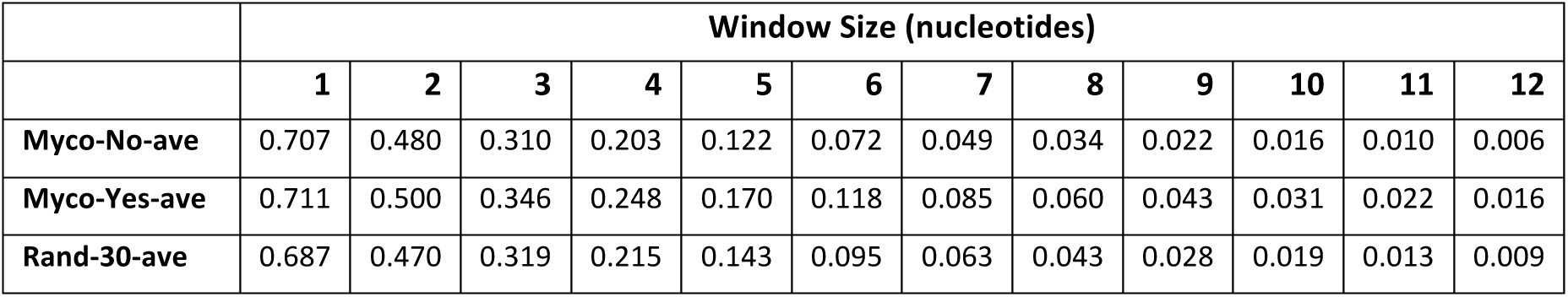
Myco-No set has lower frequency of longer contiguous A+T windows (sized 1-12 nt) composed entirely of A+T bases relative to Myco-Yes and randomly generated sequences with 30% GC content. Windows advanced by 1 nt each step until it reached the end of the sequence. “Myco-No” set consisted of Myco-28-500-2, Myco-30-500-2, and Myco-30-700-2. “Myco-Yes” set consisted of Myco-28-500-1, Myco-28-700-1, Myco-28-700-2, Myco-28-1000-1, Myco-28-1000-2, Myco-30-500-1, Myco-30-700-1, Myco-30-1000-1, Myco-30-1000-2. The “random-30” set consisted of 29 randomly generated sequences of approximately 30% GC. Values are normalized to correct for size differences between the fragments.

**Suplementary Table 8:**
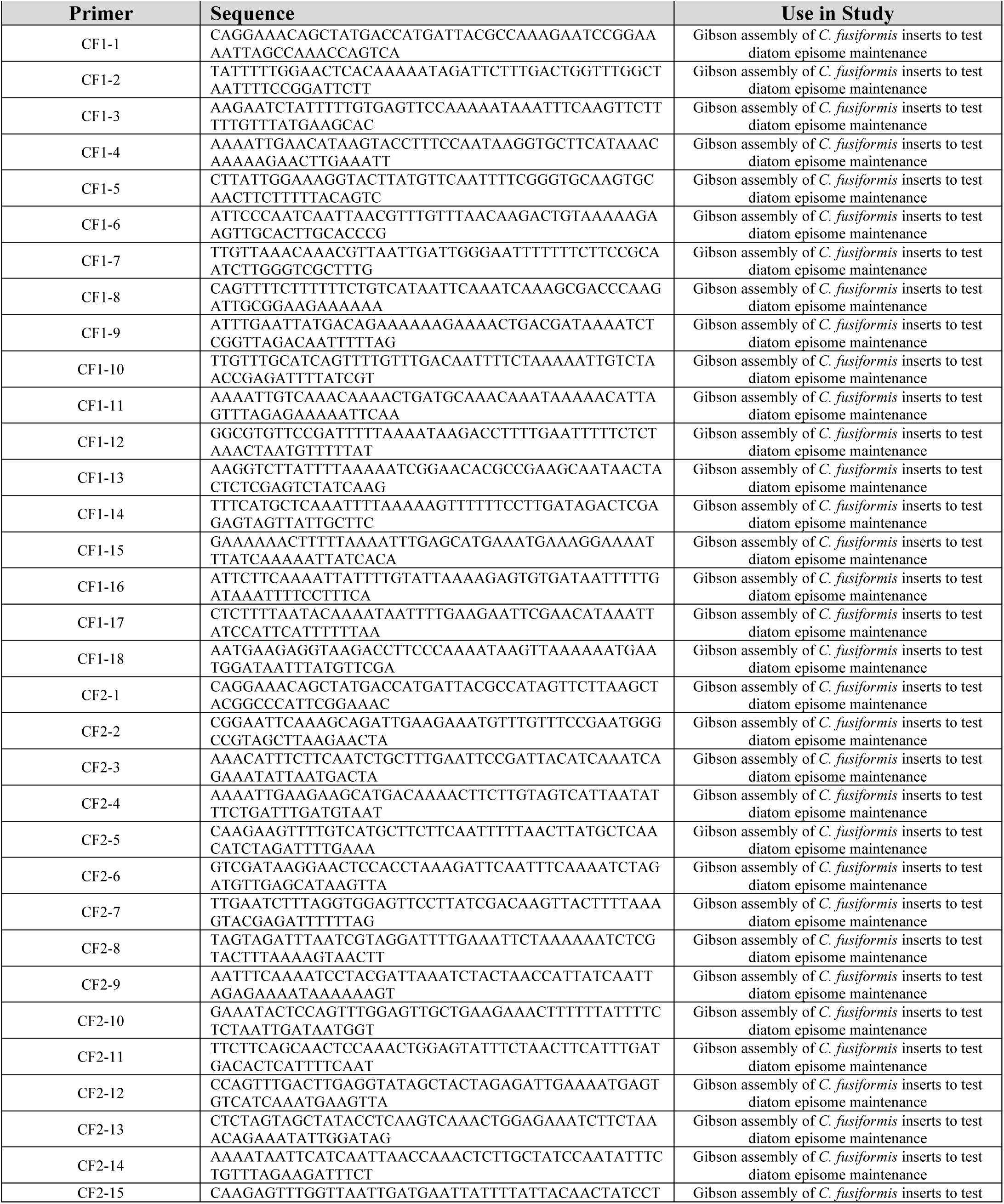
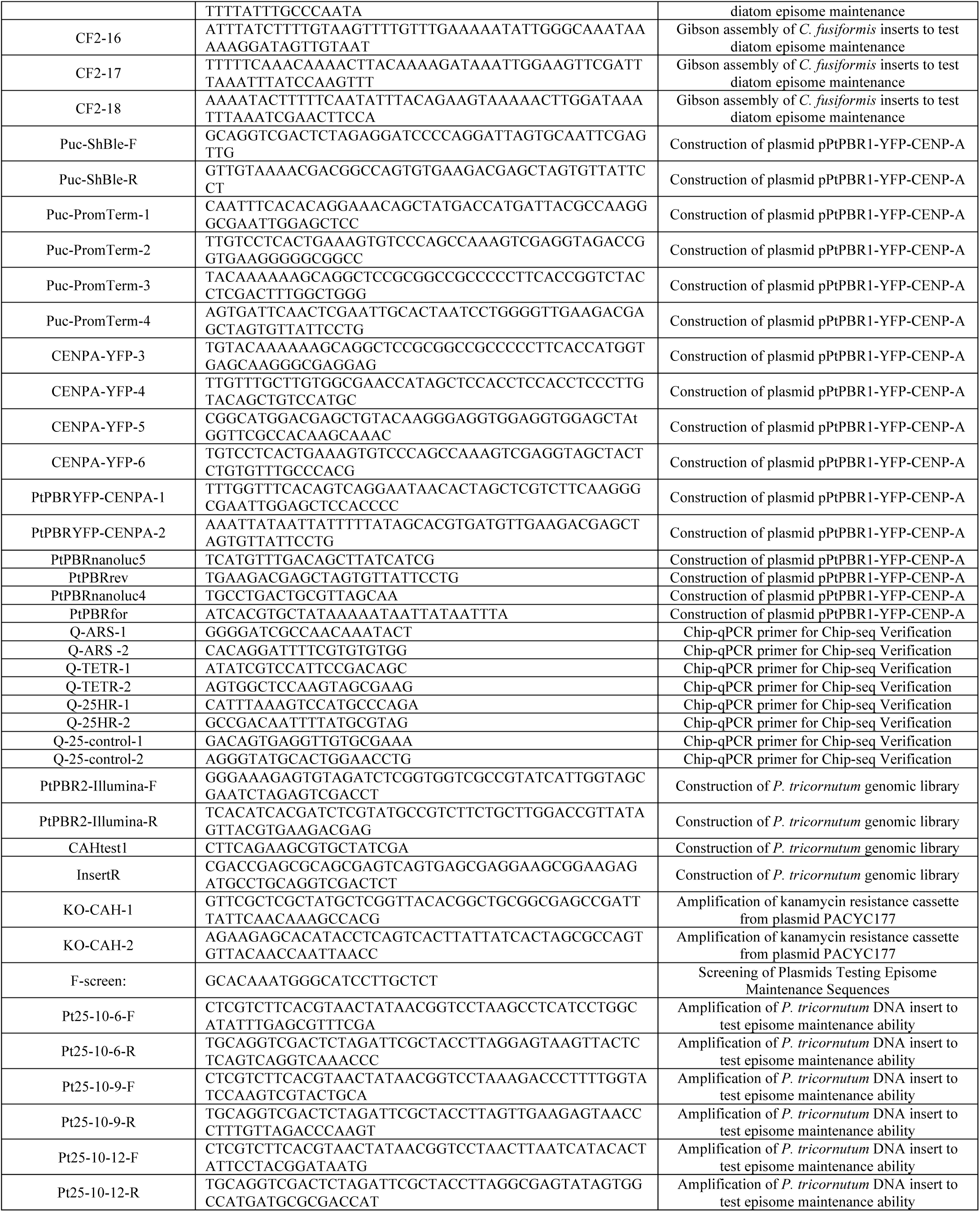
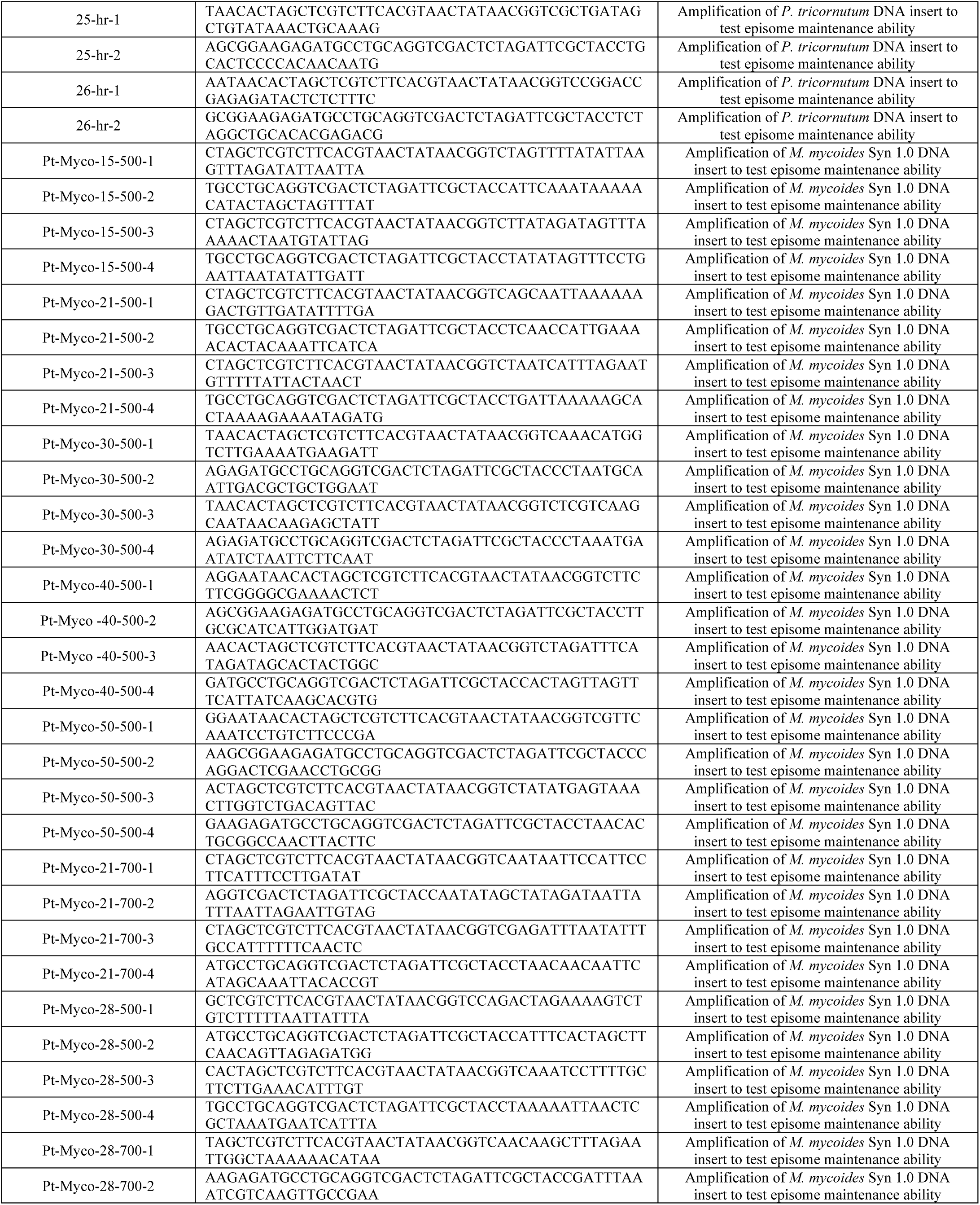
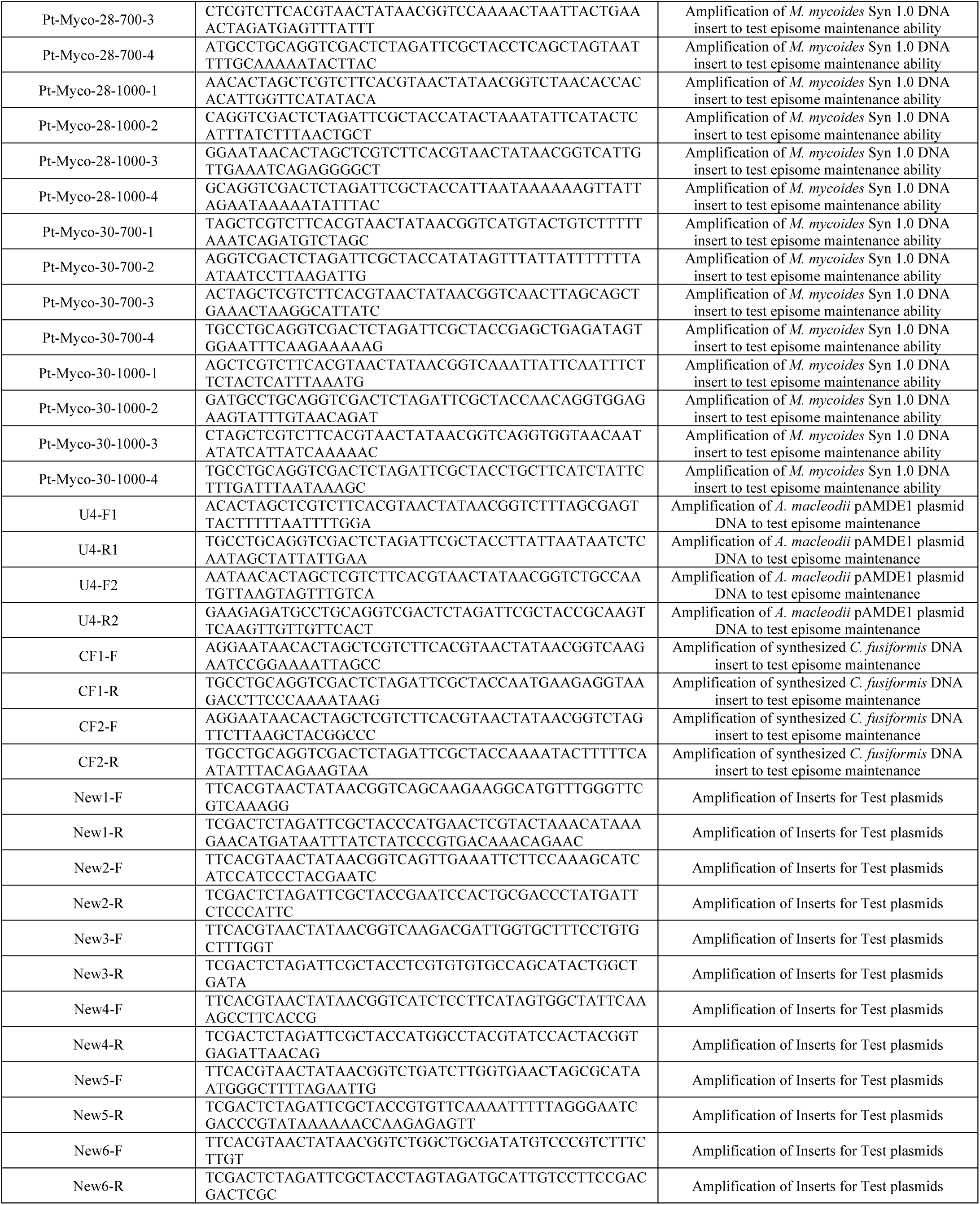
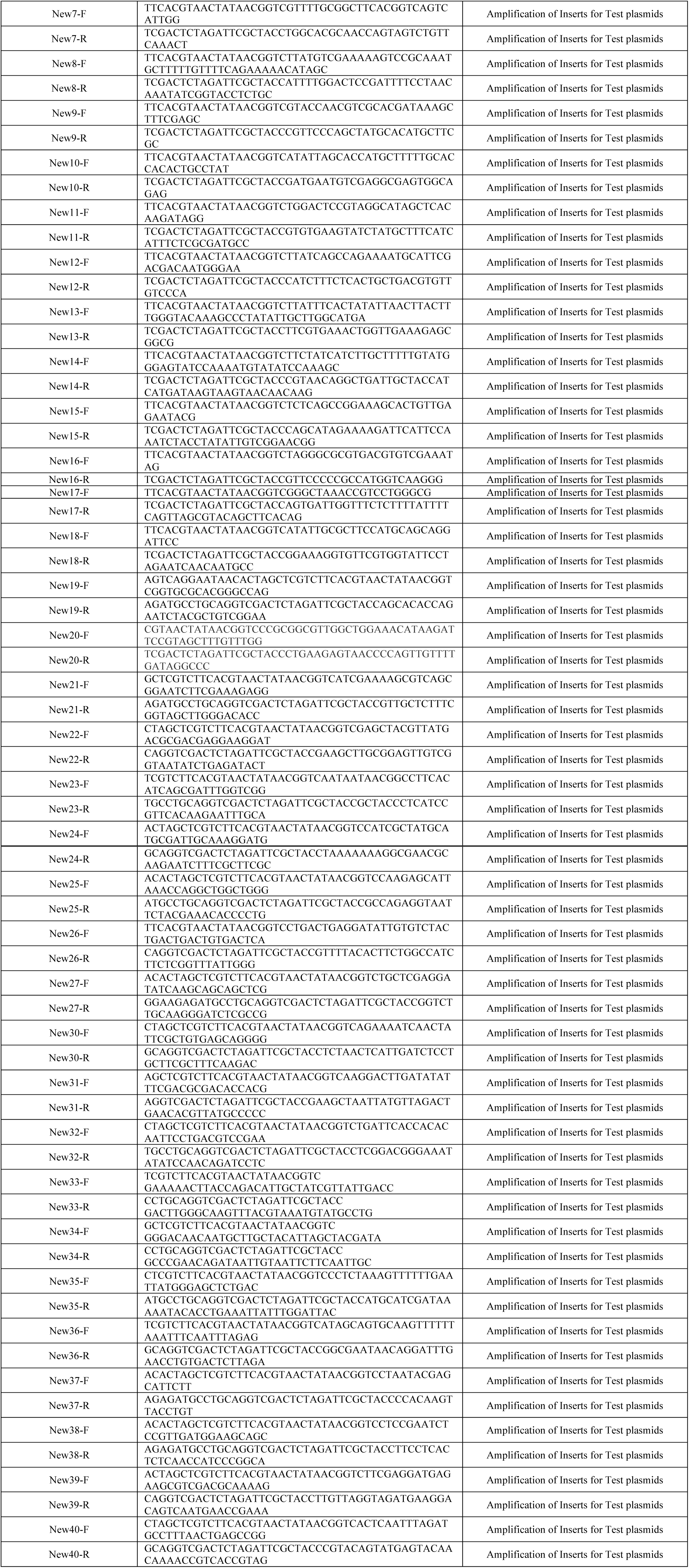
Sequence of primers used in this study

**Suplementl Table 9:**
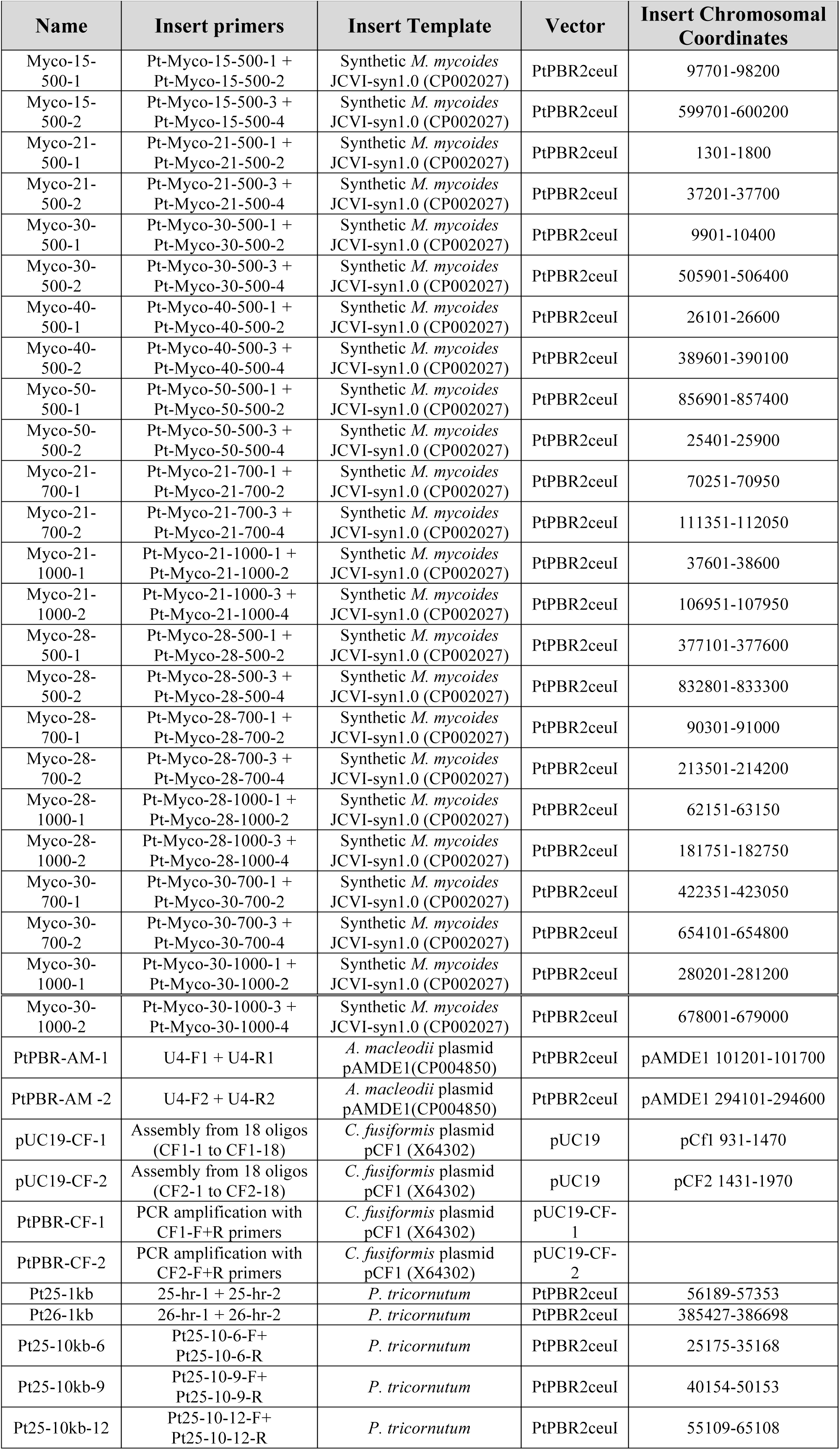
Plasmid characteristics and assembly details: Plasmids constructed to test maintenance ability of DNA sequences, including names of primers and template used to amplify insert plasmid sequences, the backbone vector used, the species the sequence originates from, and the sequence coordinates of the cloned insert. All plasmids were constructed using Gibson assembly.

## References

Albertson, D. G., and Thomson, J. N. (1982). The kinetochores of Caenorhabditis elegans. Chromosoma 86, 409–428. doi:10.1007/BF00292267.

Altschul, S. F., Gish, W., Miller, W., Myers, E. W., and Lipman, D. J. (1990). Basic local alignment search tool. J. Mol. Biol. 215, 403–10. doi:10.1016/S0022-2836(05)80360-2.

Archibald, J. M. (2009). The Puzzle of Plastid Evolution. Curr. Biol. 19, R81–R88. doi:10.1016/j.cub.2008.11.067.

Armbrust, E. V. (2009). The life of diatoms in the world’s oceans. Nature 459, 185–92. doi:10.1038/nature08057.

Armbrust, E. V., Berges, J. a, Bowler, C., Green, B. R., Martinez, D., Putnam, N. H., et al. (2004). The genome of the diatom Thalassiosira pseudonana: ecology, evolution, and metabolism. Science 306, 79–86. doi:10.1126/science.1101156.

Bailey, T. L., Boden, M., Buske, F. A., Frith, M., Grant, C. E., Clementi, L., et al. (2009). MEME Suite: Tools for motif discovery and searching. Nucleic Acids Res. 37, 1–7. doi:10.1093/nar/gkp335.

Benson, G. (1999). Tandem Repeats Finder: a program to analyse DNA sequences. Nucleic Acids Res. 27, 573–578.

Bowler, C., Allen, A. E., Badger, J. H., Grimwood, J., Jabbari, K., Kuo, A., et al. (2008). The Phaeodactylum genome reveals the evolutionary history of diatom genomes. Nature 456, 239–44. doi:10.1038/nature07410.

Bowman, S., Lawson, D., Basham, D., Brown, D., Chillingworth, T., Churcher, C. M., et al. (1999). The complete nucleotide sequence of chromosome 3 of Plasmodium falciparum. Nature 400, 532–8. doi:10.1038/22964.

Bozarth, A., Maier, U. G., and Zauner, S. (2009). Diatoms in biotechnology: Modern tools and applications. Appl. Microbiol. Biotechnol. 82, 195–201. doi:10.1007/s00253-008-1804-8.

Cheeseman, I. M., and Desai, A. (2008). Molecular architecture of the kinetochore–microtubule interface. Nat. Rev. Mol. Cell Biol. 9, 33–46. doi:10.1038/nrm2310.

Clarke, L., Amstutz, H., Fishel, B., and Carbon, J. (1986). Analysis of centromeric DNA in the fission yeast Schizosaccharomyces pombe. Proc. Natl. Acad. Sci. U. S. A. 83, 8253–7. doi:10.1073/pnas.83.21.8253.

Clarke, L., and Carbon, J. (1980). Isolation of a yeast centromere and construction of functional small circular chromosomes. Nature 287, 504–509. doi:10.1038/287504a0.

Cleveland, D. W., Mao, Y., and Sullivan, K. F. (2003). Centromeres and kinetochores: From epigenetics to mitotic checkpoint signaling. Cell 112, 407–421. doi:10.1016/S0092-8674(03)00115-6.

Cottarel, G., Shero, J. H., Hieter, P., and Hegemann, J. H. (1989). A 125 bp CEN6 DNA fragment is sufficient for complete meiotic and mitotic centromere functions in Saccharomyces cerevisiae. Trends Genet. 5, 322–324. doi:10.1016/0168-9525(89)90119-4.

Coudreuse, D. (2009). Insights from synthetic yeasts. Yeast 26, 545–551. doi:10.1002/yea.

Cuacos, M., H. Franklin, F. C., and Heckmann, S. (2015). Atypical centromeres in plants—what they can tell us. Front. Plant Sci. 6, 1–15. doi:10.3389/fpls.2015.00913.

Datsenko, K. A., and Wanner, B. L. (2000). One-step inactivation of chromosomal genes in Escherichia coli K-12 using PCR products. Proc. Natl. Acad. Sci. U. S. A. 97, 6640–5. doi:10.1073/pnas.120163297.

Diner, R. E., Bielinski, V. A., Dupont, C., Allen, A. E., and Weyman, P. D. (2016). Refinement of the Diatom Episome Maintenance Sequence and Improvement of Conjugation-based DNA Delivery Methods. Front. Bioeng. Biotechnol. 4, 65. doi:10.3389/FBIOE.2016.00065.

Earnshaw, W. C., Allshire, R. C., Black, B. E., Bloom, K., Brinkley, B. R., Brown, W., et al. (2013). Esperanto for histones: CENP-A, not CenH3, is the centromeric histone H3 variant. Chromosom. Res. 21, 101–106. doi:10.1007/s10577-013-9347-y.

Falciatore, A., Casotti, R., Leblanc, C., Abrescia, C., and Bowler, C. (1999). Transformation of Nonselectable Reporter Genes in Marine Diatoms. Mar. Biotechnol. 1, 239–251. doi:10.1007/PL00011773.

Fu, W., Wichuk, K., and Brynjólfsson, S. (2015). Developing diatoms for value-added products: Challenges and opportunities. N. Biotechnol. 32, 547–551. doi:10.1016/j.nbt.2015.03.016.

Gibson, D. G., Young, L., Chuang, R.-Y., Venter, J. C., Hutchison, C. a, and Smith, H. O. (2009). Enzymatic assembly of DNA molecules up to several hundred kilobases. Nat. Methods 6, 343–345. doi:10.1038/nmeth.1318.

Hadlaczky, G., Praznovszky, T., Cserpán, I., Keresö, J., Péterfy, M., Kelemen, I., et al. (1991). Centromere formation in mouse cells cotransformed with human DNA and a dominant marker gene. Proc. Natl. Acad. Sci. U. S. A. 88, 8106–10. Available at: http://www.pubmedcentral.nih.gov/articlerender.fcgi?artid=52455&tool=pmcentrez&rendertype=abstract.

Harrington, J. J., VanBokken, G., Mays, R. W., Gustashaw, K., and Willard, H. F. (1997). Formation of de novo centromeres and construction of first-generation human artificial microchromosomes. Nat. Genet. 15, 57–61.

Henikoff, S., Ahmad, K., and Malik, H. S. (2001). The centromere paradox: stable inheritance with rapidly evolving DNA. Science 293, 1098–1102. doi:10.1126/science.1062939.

Hildebrand, M., Hasegawa, P., Ord, R. W., Thorpe, V. S., Glass, C. A., and Volcani, B. E. (1992). Nucleotide-Sequence of Diatom Plasmids - Identification of Open Reading Frames with Similarity to Site-Specific Recombinases. Plant Mol. Biol. 19, 759–770. doi:Doi 10.1007/Bf00027072.

Iwanaga, S., Kato, T., Kaneko, I., and Yuda, M. (2012). Centromere plasmid: A new genetic tool for the study of Plasmodium falciparum. PLoS One 7. doi:10.1371/journal.pone.0033326.

Iwanaga, S., Khan, S. M., Kaneko, I., Christodoulou, Z., Newbold, C., Yuda, M., et al. (2010). Functional Identification of the Plasmodium Centromere and Generation of a Plasmodium Artificial Chromosome. Cell Host Microbe 7, 245–255. doi:10.1016/j.chom.2010.02.010.

Jacobs, J. D., Ludwig, J. R., Hildebrand, M., Kukel, A., Feng, T. Y., Ord, R. W., et al. (1992). Characterization of two circular plasmids from the marine diatom Cylindrotheca fusiformis: plasmids hybridize to chloroplast and nuclear DNA. MGGMol. Gen. Genet. 233, 302–310. doi:10.1007/BF00587592.

Kanesaki, Y., Imamura, S., Matsuzaki, M., and Tanaka, K. (2015). Identification of centromere regions in chromosomes of a unicellular red alga, Cyanidioschyzon merolae. FEBS Lett. 589, 1219–1224. doi:10.1016/j.febslet.2015.04.009.

Kapoor, S., Zhu, L., Froyd, C., Liu, T., and Rusche, L. N. (2015). Regional centromeres in the yeast Candida lusitaniae lack pericentromeric heterochromatin. Proc. Natl. Acad. Sci. U. S. A. 112, 12139–44. doi:10.1073/pnas.1508749112.

Karas, B. J., Diner, R. E., Lefebvre, S. C., McQuaid, J., Phillips, A. P. R., Noddings, C. M., et al. (2015). Designer diatom episomes delivered by bacterial conjugation. Nat. Commun. 6, 6925. doi:10.1038/ncomms7925.

Karas, B. J., Molparia, B., Jablanovic, J., Hermann, W. J., Lin, Y.-C., Dupont, C. L., et al. (2013). Assembly of eukaryotic algal chromosomes in yeast. J. Biol. Eng. 7, 30. doi: 10.1186/1754-1611-7-30.

Kouprina, N., Tomilin, A. N., Masumoto, H., Earnshaw, W. C., and Larionov, V. (2014). Human artificial chromosome-based gene delivery vectors for biomedicine and biotechnology. Expert Opin. Drug Deliv. 11, 517–35. doi:10.1517/17425247.2014.882314.

Lin, X., Tirichine, L., Bowler, C., Round, F., Crawford, R., Mann, D., et al. (2012). Protocol: Chromatin immunoprecipitation (ChIP) methodology to investigate histone modifications in two model diatom species. Plant Methods 8, 48. doi:10.1186/1746-4811-8-48.

Liu, W., Yuan, J. S., and Stewart Jr, C. N. (2013). Advanced genetic tools for plant biotechnology. Nat Rev Genet 14, 781–793. doi:10.1038/nrg3583.

López-Pérez, M., Gonzaga, A., and Rodriguez-Valera, F. (2013). Genomic diversity of “deep ecotype” Alteromonas macleodii isolates: Evidence for pan-mediterranean clonal frames. Genome Biol. Evol. 5, 1220–1232. doi:10.1093/gbe/evt089.

Lopez, P. J., Desclés, J., Allen, A. E., and Bowler, C. (2005). Prospects in diatom research. Curr. Opin. Biotechnol. 16, 180–6. doi:10.1016/j.copbio.2005.02.002.

Lynch, D. B., Logue, M. E., Butler, G., and Wolfe, K. H. (2010). Chromosomal G + C content evolution in yeasts: Systematic interspecies differences, and GC-poor troughs at centromeres. Genome Biol. Evol. 2, 572–583. doi:10.1093/gbe/evq042.

Ma, J., Wing, R. a., Bennetzen, J. L., and Jackson, S. a. (2007). Plant centromere organization: a dynamic structure with conserved functions. Trends Genet. 23, 134–139. doi:10.1016/j.tig.2007.01.004.

Malik, H. S., and Henikoff, S. (2002). Conflict begets complexity: The evolution of centromeres. Curr. Opin. Genet. Dev. 12, 711–718. doi:10.1016/S0959-437X(02)00351-9.

Malik, H. S., and Henikoff, S. (2009). Major Evolutionary Transitions in Centromere Complexity. Cell 138, 1067–1082. doi:10.1016/j.cell.2009.08.036.

Martin, W. (2003). Gene transfer from organelles to the nucleus: frequent and in big chunks. Proc. Natl. Acad. Sci. U. S. A. 100, 8612–4. doi:10.1073/pnas.1633606100.

Maruyama, S., Matsuzaki, M., Kuroiwa, H., Miyagishima, S.-Y., Tanaka, K., Kuroiwa, T., et al. (2008). Centromere structures highlighted by the 100%-complete Cyanidioschyzon merolae genome. Plant Signal. Behav. 3, 140–1. doi:10.1186/1741-7007-5-28.140.

McKinley, K. L., and Cheeseman, I. M. (2016). The molecular basis for centromere identity and function. Nat Rev Mol Cell Biol 17, 16–29. doi:10.1038/nrm.2015.5.

Monaco, A. P., and Larin, Z. (1994). YACs, BACs, PACs and MACs: Artificial chromosomes as research tools. Trends Biotechnol. 12, 280–286. doi:10.1016/0167-7799(94)90140-6.

Murray, A. W., and Szostak, J. W. (1983). Construction of artificial chromosomes in yeast. J. Chem. Inf. Model. 305, 189–193. doi:10.1017/CBO9781107415324.004.

Nakaseko, Y., Adachi, Y., Funahashi, S., Niwa, O., and Yanagida, M. (1986). Chromosome walking shows a highly homologous repetitive sequence present in all the centromere regions of fission yeast. EMBO J. 5, 1011–21. Available at: http://www.pubmedcentral.nih.gov/articlerender.fcgi?artid=1166895&tool=pmcentrez&rendertype=abstract.

Neumann, P., Navrátilová, A., Schroeder-Reiter, E., Koblížková, A., Steinbauerová, V., Chocholová, E., et al. (2012). Stretching the rules: Monocentric chromosomes with multiple centromere domains. PLoS Genet. 8. doi:10.1371/journal.pgen.1002777.

Pluta, A. F., Mackay, A. M., Ainsztein, I. G., Goldberg, I. G., and Earnshaw, W. C. (1995). The Centromere: Hub of Chromosomal Activities. Science (80-.). 270, 1591.

Price, N. M., Harrison, G. I., Hering, J. G., Hudson, R. J., Nirel, P. M. V., Palenik, B., et al. (1989). Preparation and Chemistry of the Artificial Algal Culture Medium Aquil. Biol. Oceanogr. 6, 443–461. doi:10.1080/01965581.1988.10749544.

Ravi, M., and Chan, S. W. L. (2010). Haploid plants produced by centromere-mediated genome elimination. Nature 464, 615–8. doi:10.1038/nature08842.

Rochaix, J. D., van Dillewijn, J., and Rahire, M. (1984). Construction and characterization of autonomously replicating plasmids in the green unicellular alga Chlamydomonas reinhardii. Cell 36, 925–931. doi:10.1016/0092-8674(84)90042-4.

Sato, H., Masuda, F., Takayama, Y., Takahashi, K., and Saitoh, S. (2012). Epigenetic inactivation and subsequent heterochromatinization of a centromere stabilize dicentric chromosomes. Curr. Biol. 22, 658–667. doi:10.1016/j.cub.2012.02.062.

Smith, K. M., Galazka, J. M., Phatale, P. A., Connolly, L. R., and Freitag, M. (2012). Centromeres of filamentous fungi. Chromosom. Res. 20, 635–656. doi:10.1007/s10577-012-9290-3.

Stimpson, K. M., Matheny, J. E., and Sullivan, B. A. (2012). Dicentric chromosomes: Unique models to study centromere function and inactivation. Chromosom. Res. 20, 595–605. doi:10.1007/s10577-012-9302-3.

Sullivan, B. a, Blower, M. D., and Karpen, G. H. (2001). Determining centromere identity: cyclical stories and forking paths. Nat. Rev. Genet. 2, 584–596. doi:10.1038/35084512.

Sullivan, B. a, and Willard, H. F. (1998). Stable dicentric X chromosomes with two functional centromeres. Nat. Genet. 20, 227–228. doi:10.1038/3024.

Timmis, J. N., Ayliffe, M. a, Huang, C. Y., and Martin, W. (2004). Endosymbiotic gene transfer: organelle genomes forge eukaryotic chromosomes. Nat. Rev. Genet. 5, 123–35. doi:10.1038/nrg1271.

Torras-Llort, M., Moreno-Moreno, O., and Azorín, F. (2009). Focus on the centre: the role of chromatin on the regulation of centromere identity and function. EMBO J. 28, 2337–2348. doi:10.1038/emboj.2009.174.

Tyler-Smith, C., Oakey, R., Z., L., Fisher, R., Crocker, M., Affara, N., et al. (1993). Localisation of DNA sequences requires for human centromere function through an analysis of rearranges Y chromosomes. Nat. Genet. 5, 368–375.

Wada, N., Kazuki, Y., Kazuki, K., Inoue, T., Fukui, K., and Oshimura, M. (2016). Maintenance and Function of a Plant Chromosome in Human Cells. ACS Synth. Biol., acssynbio.6b00180. doi:10.1021/acssynbio.6b00180.

Wang, G., Li, H., Cheng, Z., and Jin, W. (2013). A novel translocation event leads to a recombinant stable chromosome with interrupted centromeric domains in rice. Chromosoma 122, 295–303. doi:10.1007/s00412-013-0413-1.

Westermann, S., Cheeseman, I. M., Anderson, S., Yates, J. R., Drubin, D. G., and Barnes, G. (2003). Architecture of the budding yeast kinetochore reveals a conserved molecular core. J. Cell Biol. 163, 215–222. doi:10.1083/jcb.200305100.

Westhorpe, F. G., Straight, A. F., Varetti, G., Pellman, D., David, J., and Cheeseman, I. M. (2014). The Centromere: Epigenetic Control of Chromosome Segregation during Mitosis. Cold Spring Harb. Perspect. Biol. 7, 1–26. doi:a015818.

Willard, H. F. (1998). Centromeres: The missing link in the development of human artificial chromosomes. Curr. Opin. Genet. Dev. 8, 219–225. doi:10.1016/S0959-437X(98)80144-5.

Yu, W., Yau, Y., and Birchler, J. A. (2016). Plant artificial chromosome technology and its potential application in genetic engineering. Plant Biotechnol. J. 14, 1175–82. doi:10.1111/pbi.12466.

Zhang, W., Friebe, B., Gill, B. S., and Jiang, J. (2010). Centromere inactivation and epigenetic modifications of a plant chromosome with three functional centromeres. Chromosoma 119, 553–563. doi:10.1007/s00412-010-0278-5.

